# Intervention-consistent causal-source recovery from covariance-response geometry reveals upstream organisation in sporadic ALS

**DOI:** 10.64898/2026.05.16.716261

**Authors:** Shunya Kaneko, Makoto Urushitani

## Abstract

Sporadic amyotrophic lateral sclerosis (sALS) lacks longitudinal molecular measurements, making it difficult to distinguish early disease-organising changes from downstream consequences. We present a training-free framework that extracts directional structure from static single-nucleus RNA-seq by applying discrete Hodge decomposition to gene co-expression dynamics across pseudotime-ordered donor states. The framework separates irreversible co-expression cascades from circular feedback structure and regresses out the component explained by the healthy co-expression network, allowing disease-specific organisation to be examined in isolation. Perturbation benchmarks show that experimentally imposed sources are recoverable from control-normalised off-diagonal covariance-response fields, whereas marginal variance and diagonal covariance controls do not recover the source.

Applied to sALS primary motor cortex (24 donors, 10 cell types), the framework identifies oligodendrocytes as the most structurally upstream cell type and upper-motor-neuron-containing layers as the most structurally downstream (Oligo cell-type *φ* = 0.900, with glial cell types preserving the healthy co-expression network topology, whereas neuronal cell types show collapse-dominant deformation). Cytoplasmic translation is the only pathway with reproducible cross-cell-type upstream enrichment. At the gene level, the ribosome-associated quality-control factor NEMF — which appends C-terminal alanine-threonine tags (“CATylation”) to nascent chains on stalled ribosomes — shows disease-specific loss of co-expression coherence in seven of ten cell types despite essentially unchanged mRNA expression; the disease signal is decoupling from collision-response partners (GCN2, PKR), not expression-level change.

Cross-cohort validation across three BA4 motor cortex cohorts (including two external cohorts; total N=107) reproduced the oligodendrocyte-upstream / upper-motor-neuron-downstream structural architecture (Oligo-preserved / ET-sink) in all three cohorts, with NEMF co-expression coherence loss replicated in two of three cohorts. These data support a brain-side, circuit-distal structural model in which oligodendrocyte-lineage stress occupies an upstream-like preserved compartment, while upper-motor-neuron-containing excitatory populations form a downstream sink. The pattern is consistent with — but does not directly establish — a cascade architecture in which oligodendrocyte stress structurally precedes motor neuron TDP-43 pathology, and would produce a clinical phenotype resembling dying-back (the conventional view of ALS, in which motor neuron pathology appears to begin at distal axons and spread retrogradely toward the cell body) yet originating centrally and glially. NEMF/CATylation network disruption is identified as a candidate intermediate structural node bridging oligodendrocyte stress and motor neuron TDP-43 pathology.

## Introduction

Amyotrophic lateral sclerosis (ALS) is a fatal neurodegenerative disease characterised by progressive motor neuron loss, and the causal sequence linking initial cellular triggers, intermediate molecular events, and final neuron loss remains unresolved. Post-mortem tissue provides only a single snapshot per donor over a 2–5 year disease course, with no longitudinal molecular measurements at the cellular level, making it intrinsically difficult to distinguish early disease-organising programmes from downstream consequences. Existing analytical methods address parts of this challenge but not all of it. Differential expression analysis reports magnitude without direction. Gene regulatory network inference (SCENIC+, Bravo González-Blas,. 2023; GRNBoost2, Moerman et al. 2019; CellOracle, Kamimoto et al. 2023) requires chromatin accessibility or perturbation training data. Trajectory inference (RNA velocity, La Manno et al. 2018; Bergen et al. 2020; CellRank, Lange et al. 2022) reconstructs pseudotemporal orderings of cells but does not identify causal relationships between genes. Perturbation-prediction methods (GEARS, Roohani et al. 2024) answer “what happens if gene X is perturbed?” rather than “which gene is structurally upstream?”, and recent evaluations show no deep-learning perturbation predictor significantly outperforms linear baselines (Ahlmann-Eltze et al. 2025). Hodge-theoretic methods have begun to appear in trajectory inference on cell-state graphs (PHLOWER; Cheng et al. 2025), but none offers a training-free decomposition of gene co-expression changes into directional and feedback components.

Here we introduce an analytical instrument that transforms snapshot single-nucleus RNA-seq data into a structural upstream readout in five stages: per-donor symmetric positive-definite (SPD) covariance estimation with Ledoit–Wolf shrinkage (Ledoit & Wolf 2004), log-Euclidean tangent-space mapping (Arsigny et al. 2006), donor pseudotime by diffusion map on inter-donor covariance distances (Coifman & Lafon 2006; Haghverdi et al. 2016), antisymmetric edge-weight flow construction on per-window covariance changes, and discrete Hodge decomposition (Eckmann 1944; Lim 2020) of the resulting edge flow into orthogonal gradient (irreversible cascade), curl (oscillatory feedback), and harmonic components. The variance-asymmetry premise underlying the construction is validated using a published RNA time-stamping system (TimeVault; Chao et al. 2026), Norman Perturb-seq for causal-source recovery (Norman et al. 2019), spatial CRISPR perturbation (Shen et al. 2026), and cross-disease reproducibility in glioma (TCGA + GTEx; Vivian et al. 2017; Goldman et al. 2020). The instrument is applied to a sALS discovery cohort (NYGC primary motor cortex, 24 donors; Pineda et al. 2024) and tested in two independent cohorts (Takeuchi et al. 2025; Ruf et al. 2026; total N=107 donors).

The central empirical thesis of this work is that intervention-consistent structural cascade direction is not contained in static covariance itself, but becomes recoverable in the control-normalised covariance-response field. Perturbation benchmarks show that experimentally imposed sources appear as centres of off-diagonal covariance-response aggregation, whereas marginal variance and diagonal-multivariate residual controls do not recover the source (Section 2; Supplementary Note 6). We use the Hodge decomposition to give this response geometry source/sink semantics, to recover the latent gradient potential on sparse graphs via Laplacian smoothing, and to define a topology-residual (3φ) framework for transferring the validated readout to static sALS cohorts. Mathematically, natural-gradient relaxation on the log-Euclidean tangent space yields a directed response field Δ, from which an antisymmetric edge flow is constructed and decomposed via Hodge theory into orthogonal gradient (cascade), curl (feedback), and harmonic components; throughout this paper, φ is reported under a source-oriented convention in which larger φ denotes more upstream/source-like position (see Methods; Supplementary Note 7). Norman Perturb-seq then tests whether these structural sources coincide with externally imposed CRISPR perturbation sources. A broader cone–manifold principle-theoretic derivation of this construction is provided in a companion theoretical manuscript.

During the preparation of this manuscript, an independent preprint reported a related Riemannian covariance-tensor approach for Parkinson’s disease (Choi et al. 2026), representing single-cell transcriptomic states as covariance objects on the SPD manifold and using Log-Euclidean geometry, von Neumann entropy, and spectral eigendecomposition of differential covariance networks to identify a bifurcation state and tensor hub drivers in PD prefrontal cortex. The present study addresses a distinct problem at a different methodological layer: whether control-whitened off-diagonal covariance-response fields can recover experimentally imposed perturbation sources (Norman Perturb-seq CRISPR ground truth), and whether this intervention-validated response geometry can be residualised against healthy co-expression topology to reveal disease-specific structural cascade organisation in sALS. Where Choi et al. extract a Riemannian hub-collapse readout from SPD covariance tensors, the present framework operates on the covariance-response field and grounds its source/sink semantics in CRISPR-validated structural causality.

We find that oligodendrocytes are the sole structurally upstream cell type in sALS primary motor cortex and that the ribosome-associated quality-control factor NEMF, unchanged in mRNA expression, is the single gene with the strongest disease-specific loss of co-expression coherence across cell types, identifying central oligodendrocyte translational stress and NEMF/CATylation network disruption as candidate upstream nodes in a brain-origin, dying-back–like cascade converging on motor-neuron TDP-43 pathology.

## Results

### 1. A Hodge-decomposition framework for inferring structural upstream organisation

The framework (Fig. 1) transforms static single-nucleus RNA-seq data into a per-gene upstream readout in five stages. ***Stage 1: Covariance on the SPD manifold***. Per-donor gene–gene correlation matrices (Ledoit–Wolf shrinkage, k = 30 PCs) are mapped into the log-Euclidean tangent space via matrix logarithm. ***Stage 2: Donor pseudotime***. Since post-mortem data lack true time-series, donors are ordered by diffusion pseudotime (DPT) computed on the inter-donor log-Euclidean distance matrix; the 24 NYGC donors (14 sALS + 10 PN, after joint cell-count QC) are partitioned into six windows of four donors each. PN donors have lower mean pseudotime than sALS (−0.016 vs +0.016) but the difference is not significant (Mann–Whitney p = 0.232) and the two groups are interleaved along the axis, confirming that pseudotime captures a continuous covariance-structural axis rather than a binary disease/control separation. ***Stage 3: Edge-weight flow***. For consecutive windows, the delta matrix Δ encodes how every gene pair’s correlation shifts, and an antisymmetric edge flow is constructed as f(i, j) = |Δ_ij_| × sign(d_i_ − d_j_), where d_i_ = □Δ_i,·_□_2_. ***Stage 4: Hodge decomposition***. The edge flow is decomposed into three orthogonal components: f = grad(φ) + curl + harmonic; on complete graphs the harmonic component vanishes. ***Stage 5: Dual-mode architecture***. The complete graph K_N_ provides stable rankings and multi-transition integration, while k-NN sparse graphs amplify per-transition signal and enable curl analysis.

**Figure 1.**
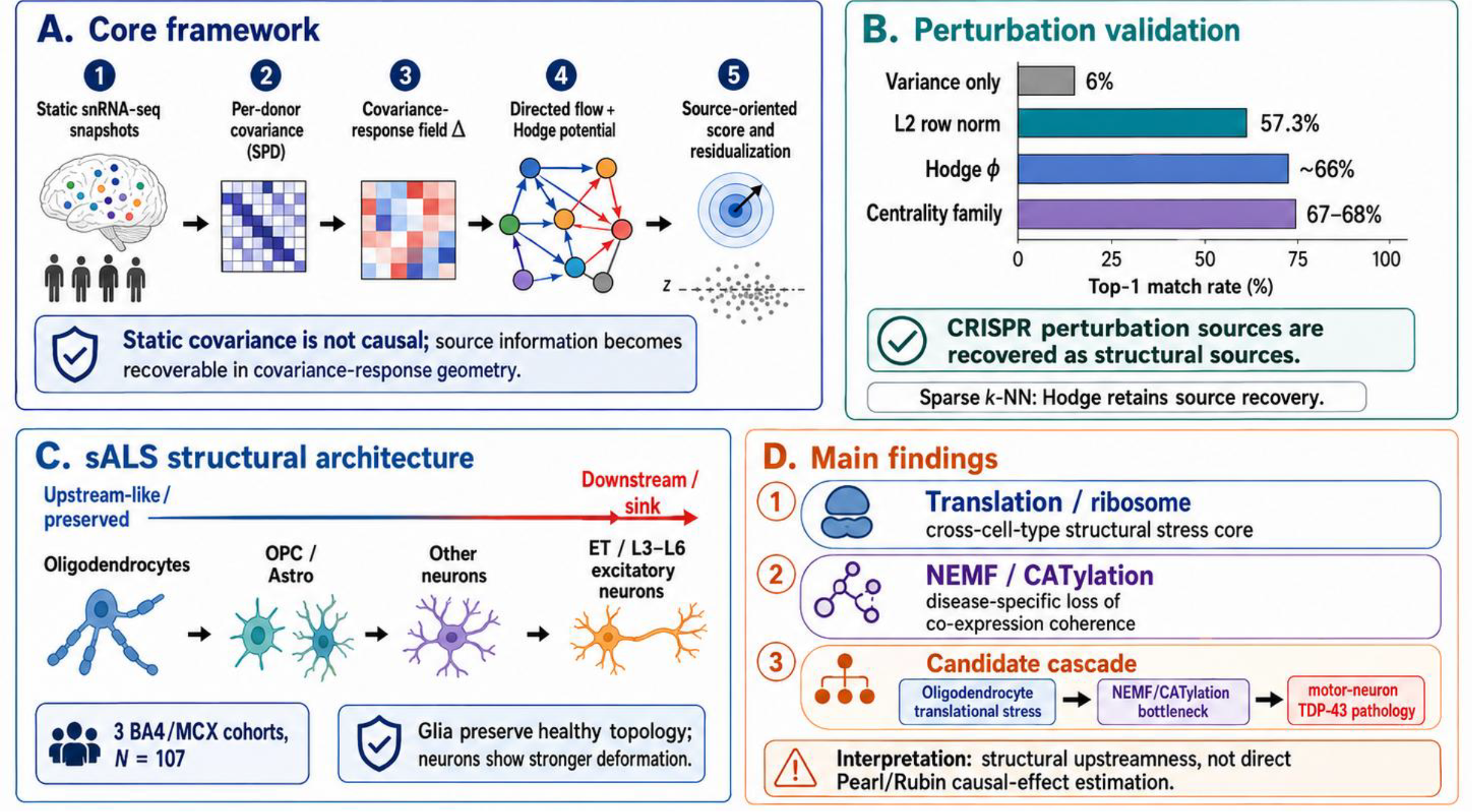
Intervention-consistent causal-source recovery from covariance-response geometry — simplified overview of framework, validation, and biological findings. (A) Core framework: five-stage pipeline transforms static single-nucleus RNA-seq snapshots into a source-oriented structural readout. Stage 1: per-donor symmetric positive-definite (SPD) covariance estimation. Stage 2: log-Euclidean tangent mapping. Stage 3: construction of the control-normalised covariance-response field Δ. Stage 4: antisymmetric directed edge flow and Hodge decomposition (gradient potential φ, curl, harmonic). Stage 5: source-oriented score and topology-residual (3φ) framework for disease-specific interpretation. Central thesis: static covariance is symmetric and does not encode direction; source information becomes recoverable in the covariance-response geometry. (B) Perturbation validation on Norman Perturb-seq (CRISPR ground truth; 232 perturbation groups after cell-count QC): variance-only baseline recovers 6.0% Top-1, L2 row norm 57.3%, Hodge φ ∼65.9%, and the broader structural-centrality family (L1 node strength, PageRank, eigenvector centrality) 67–68% Top-1. CRISPR-imposed sources are recovered as structural sources by any member of the covariance-response structural-centrality family; on sparse k-NN graphs, Hodge retains source-recovery capacity through Laplacian smoothing (Supplementary Note 6). (C) sALS structural architecture across three independent BA4 motor cortex cohorts (NYGC + Takeuchi 2025 + Ruf 2026, total N = 107): oligodendrocytes occupy the upstream-like preserved compartment; OPC and astrocytes lie intermediate; ET / L3–L6 upper-motor-neuron-containing excitatory populations form the downstream sink. Glia preserve the healthy co-expression topology, whereas neuronal cell types show stronger structural deformation. (D) Main biological findings: (1) translation / ribosome as the sole cross-cell-type primary structural stress core (10/10 cell types under 3φ matched-null z-scores); (2) NEMF, the CATylation factor of ribosome-associated quality control, shows disease-specific loss of co-expression coherence in 7 of 10 cell types despite essentially unchanged mRNA expression; (3) candidate cascade — oligodendrocyte translational stress → NEMF/CATylation bottleneck → motor-neuron TDP-43 pathology. Interpretation: the framework infers structural upstream-ness, not direct Pearl/Rubin causal-effect estimation. Overall: the readout is validated on perturbation data, then transferred to static sALS cohorts to infer disease-specific structural upstream organisation.

A closed-form null baseline for the gradient fraction GF = □grad □^2^/ □f □^2^ was established through 316 random matrix simulations and analytical derivation: **GF**_**0**_ **= 2 / [3(1 + CV**^**2**^)] (r = 0.994; Supplementary Note 4), where CV is the coefficient of variation of edge weights. This baseline operates as a built-in diagnostic: deviations of GF without corresponding CV change signal genuine biological structure beyond what edge-weight heterogeneity alone predicts.

### 2. Perturbation benchmarks validate covariance-response causal-source recovery

The premise that variance dynamics in co-expression encode recoverable information about temporal precedence was tested with TimeVault RNA time-stamping (Chao et al. 2026), which physically separates “Recorded” (past, 7 days prior) from “Present” RNA via vault-particle encapsulation and RNase treatment. A label-free expression-space pseudotime constructed from PCA on top 2,000 variable genes separated all 12 control-condition samples into their true temporal categories (AUC = 1.0, p = 0.001) while treatment effect remained non-significant (p = 1.0). A per-gene Temporal Asymmetry Score (TAS, defined in this work) discriminated TimeVault-defined upstream from downstream genes with AUC = 0.9475 (Cohen’s d = 2.06). TAS is a per-gene univariate score that does not involve gene–gene covariance or Hodge decomposition; it supports the variance-asymmetry premise rather than the SPD/Hodge instrument itself.

For causal-source recovery, we benchmarked the instrument on Norman Perturb-seq (Norman et al. 2019; 111,122 K562 cells, 235 perturbation classes, 232 groups after cell-count QC). Global whitening against control-cell covariance was decisive: Top-1 source identification accuracy was 16.4% without whitening and rose to 66.4% with whitening (median rank = 1.0; Table 1). The key finding is not that Hodge φ uniquely outperforms all alternatives on the complete graph. Rather, perturbation-source information is encoded in the control-whitened off-diagonal covariance-response field itself. On the complete graph KN, Hodge φ, L1 node strength, PageRank, and eigenvector centrality applied to the same whitened Δ matrix all recover perturbation sources at 65.9–68.1% Top-1 (Supplementary Note 6; pairwise McNemar p = 1.0 within the L1/PR/EV cluster; per-group Spearman ρ > 0.99 in 231 of 232 groups), whereas raw variance recovers only 6.0% and random ranking 0.98%. Differential-expression ranking reaches 81.0% but uses perturbation group labels — precisely the information unavailable in the observational disease setting IDS is designed for.

**Table 1.** Norman Perturb-seq upstream-ranking benchmark.

| Method | Top-1 | Median rank | Training | Perturbation labels |
| --- | --- | --- | --- | --- |
| Random | 0.98% | 51.5 | — | — |
| Variance ratio (no $\Delta$)* | 6.0% | 25.5 | None | Not used |
| d_corr (whitened) | 57.3% | 1.0 | None | Not used |
| **IDS (whitened + Hodge)** | **66.4%** | **1.0** | **None** | **Not used** |
| **L1 node strength** | **67.7%** | **1.0** | **None** | **Not used** |
| **Eigenvector centrality** | **67.7%** | **1.0** | **None** | **Not used** |
| **PageRank** | **68.1%** | **1.0** | **None** | **Not used** |
| DE ranking | 81.0% | 1.0 | None | **Required** |

\*Variance ratio: per-gene $|\log(\text{var}_{\text{pert}} / \text{var}_{\text{ctrl}})|$ applied without whitening or $\Delta$ construction, included as a label-light per-gene control to confirm that cross-gene structural aggregation — not per-gene magnitude — is the decisive operation. L1 node strength, eigenvector centrality, and PageRank are computed on the same whitened $\Delta$ matrix as IDS and d\_corr; these four 'structural family' methods (IDS $\phi$ , L1, eigenvector, PageRank) are mutually rank-equivalent on $K_N$ (median per-group Spearman $\rho > 0.99$ within the L1/PR/EV cluster; full numerical decomposition in Supplementary Note 6).

A variance-decomposition control shows that this signal is not a whitening or marginal-variance artifact (Supplementary Note 6). Raw per-gene variance reaches 6.0% Top-1; per-gene scale correction (control-normalised diagonal z) reaches 28.4%; control-conditioned residual variance via Schur complement on the control precision matrix, which preserves gene identity while subtracting linear predictability from all other genes, saturates the diagonal-multivariate limit at 19.8% Top-1 (63.8% Top-10). Off-diagonal covariance-response aggregation on the whitened SPD log-correlation Δ matrix reaches 65.9–68.1% Top-1 — a +48 percentage-point gain over the diagonal-multivariate limit. Source recovery therefore specifically requires off-diagonal covariance-response aggregation, not per-gene magnitude or whitening alone.

The Hodge framework’s distinctive ranking contribution emerges in the sparse-graph regime (Supplementary Note 6). After k-NN sparsification, unsigned graph centralities lose source-recovery capacity, while signed-flow Hodge φ remains informative. At k = 10, Hodge φ achieves 44.0% Top-1 while every centrality measure collapses to 0.4–1.3% (effectively the random baseline), with McNemar p ≤ 10 □^22^ in every comparison. A signed-divergence control on the same sparse graph, computed with the same signed flow without Laplacian inversion, shows that Hodge φ strictly dominates raw divergence at k = 10 and k = 20 (+18.6 pp and +13.4 pp Top-1; phi-only wins = 43/0 and 31/0 in discordant pairs), whereas dense graphs reconverge to exact rank equivalence (k = 100: per-group Spearman ρ = 1.000 in all 232 groups). The Hodge framework is therefore not a uniquely superior K_N_ ranking heuristic; its distinctive role is to provide source/sink semantics, sparse-graph Laplacian recovery, gradient/curl decomposition, multi-transition integration, and the 3φ residual framework — operations that no unsigned centrality measure on the same Δ matrix can supply. This is closely related to the classical HodgeRank framework (Jiang et al. 2011); the novelty of the present finding is its specific behaviour on a control-whitened covariance-response field validated against CRISPR-imposed sources.

For functional importance, the instrument was applied to spatial CRISPR data (Shen et al. 2026; 41,749 mouse hippocampal cells, 17 single-gene KOs targeting ALS/PD/AD risk genes plus mSafe). Gene Hodge was run independently within each KO condition’s cells, and each KO target’s φ percentile was measured within its own perturbation network. The 16 evaluated targets occupied a mean 21.8th percentile (source-rank percentile, where 0 denotes the most upstream position after source orientation; Wilcoxon p = 0.002), with ALS targets ranking most upstream (12.8th percentile). For cross-disease reproducibility, the identical pipeline applied to TCGA glioma + GTEx bulk RNA-seq (723 samples, 5 clinical grades) confirmed GF□ invariance (0.4248 ± 0.0004, matching sALS 0.4250). The data-driven pseudotime, constructed from covariance structure without using clinical-grade labels, recapitulated clinical staging at Spearman ρ = −0.755 with grade (AUC = 0.998 for grade discrimination), and gene-level φ rankings derived from pseudotime-based windows agreed with those from clinical-grade-based windows at ρ = 0.891. Transition-specific upstream programmes matched established glioma biology (ECM remodelling Normal → LGG; proliferation G2 → G3; synaptic acquisition G3 → IDHwt; immune remodelling IDHwt → GBM).

In sALS, whitening is not applied because the pseudotime delta construction serves an analogous role: the log-Euclidean delta log(C_2_) − log(C_1_) approximates log(C_1_^−1/2^ C_2_ C_1_^−1/2^), isolating transition-specific changes. The approximation is exact when C_1_ and C_2_ commute and holds to first order in the Baker– Campbell–Hausdorff expansion otherwise; under Ledoit–Wolf shrinkage toward the identity, the commutator is small and the approximation is empirically accurate.

### 3. Oligodendrocytes are the sole structurally upstream cell type in sALS

Among ten major cortical cell types analysed in the discovery cohort, oligodendrocytes alone emerged as upstream (φ = 0.900; 100% cell-type-level bootstrap support; gene-level bootstrap 64/100). All 24 leave-one-donor-out (LODO) iterations retained oligodendrocytes as top-ranked, and gene rankings were stable across the same leave-one-donor folds (Spearman ρ = 0.981; Top-50 Jaccard = 0.954). From 922 candidate genes, 135 stable-High genes (top quantile in ≥ 4/5 transitions) were enriched for ribosomal biogenesis and cytoplasmic translation (p = 9.7 × 10 □^1^ □).

Two sensitivity analyses confirmed robustness. First, pseudotime axes constructed without oligodendrocytes (four configurations: excitatory-neuron-only, all-neuron, glia-free) retained oligodendrocytes within the top two cell types in all configurations (Spearman ρ ≥ 0.952 versus the original axis). Second, substituting d_corr with d_var or d_cov as the pairwise distance metric did not alter the ranking (d_corr vs d_cov: ρ = 1.000). Microglia ranked second in the full-cohort analysis but showed φ = 0 in the sALS+PN cohort across all three metrics, consistent with the known difference in upstream cell type between sALS and C9orf72-ALS. The same instrument applied to the C9ALS subset of the same NYGC cohort yielded a qualitatively different cell-type ranking, providing an internal contrast that supports cohort-specific rather than uniform-technical interpretation.

### 4. The 3φ residual framework separates topology from disease-specific signal

A central finding of the manifold-level analysis is that raw *φ* on K_N_ is largely determined by healthy-state co-expression topology. On the complete graph, *φ* is practically equivalent to a monotone transformation of the row-engagement scalar d_i_ (Spearman ρ ≈ −1.000), and 82–92% of φ variance in glia reflects the weighted strength of each gene in the healthy co-expression network. The biological signal therefore does not sit in raw *φ* position but in the residuals after topology regression.

We computed *φ* three times per cell type, differing only in the donor subset used: *φ*_static_ from the 10 PN donors (zero disease information; captures the weighted strength of each gene in the healthy network), *φ*_disease_ from the 14 sALS donors, and *φ*_condition_ from the pooled panel. Per cell type, *φ*_disease_ is regressed on *φ*_static_ via cubic polynomial fit, and per-gene residual z-scores quantify topology-independent disease effects (z > +2: rewiring; z < −2: collapse). Matched-null z-scores are computed by 1,000 ALS/PN label permutations and used for the pathway- and gene-level tests reported below.

The cell-type R^2^ of the polynomial fit reveals a glia-to-neuron gradient in topology preservation (Table 2): glial cell types preserve the healthy manifold (Astro R^2^ = 0.95, OPC 0.95, Oligo 0.92), whereas neuronal cell types show progressive deformation (L2/3 0.85; L6 0.58; PV 0.68). The same comparison repeated on 16 C9orf72-ALS donors from the same NYGC cohort collapses this gradient: all cell types show R^2^ < 0.30 (Fig. 2). At the differential-expression level sALS and C9ALS appear convergent (Pineda et al. 2024); at the co-expression-structure level they are distinct.

**Table 2.** Cell-type gradient in manifold preservation (Spearman ρ^2^ between disease-state φ and PN-only φ).

| Cell type | Type | $R^2$ (disease ~ static) | Disease architecture |
| --- | --- | --- | --- |
| Astro | glia | **0.950** | Manifold preserved |
| OPC | glia | **0.946** | Manifold preserved |
| Oligo | glia | **0.922** | Manifold preserved |
| L2/L3 | neuron | 0.853 | Glia-like (high preservation) |
| L4/L6 | neuron | 0.752 | Mixed |
| L5/L6 | neuron | 0.730 | Mixed |
| L3/L5 | neuron | 0.728 | Mixed |
| PV | neuron | 0.677 | Mixed-deformed |
| L6 | neuron | 0.575 | Deformed |
| Endo | endo | 0.557 | Deformed |

**Figure 2.**
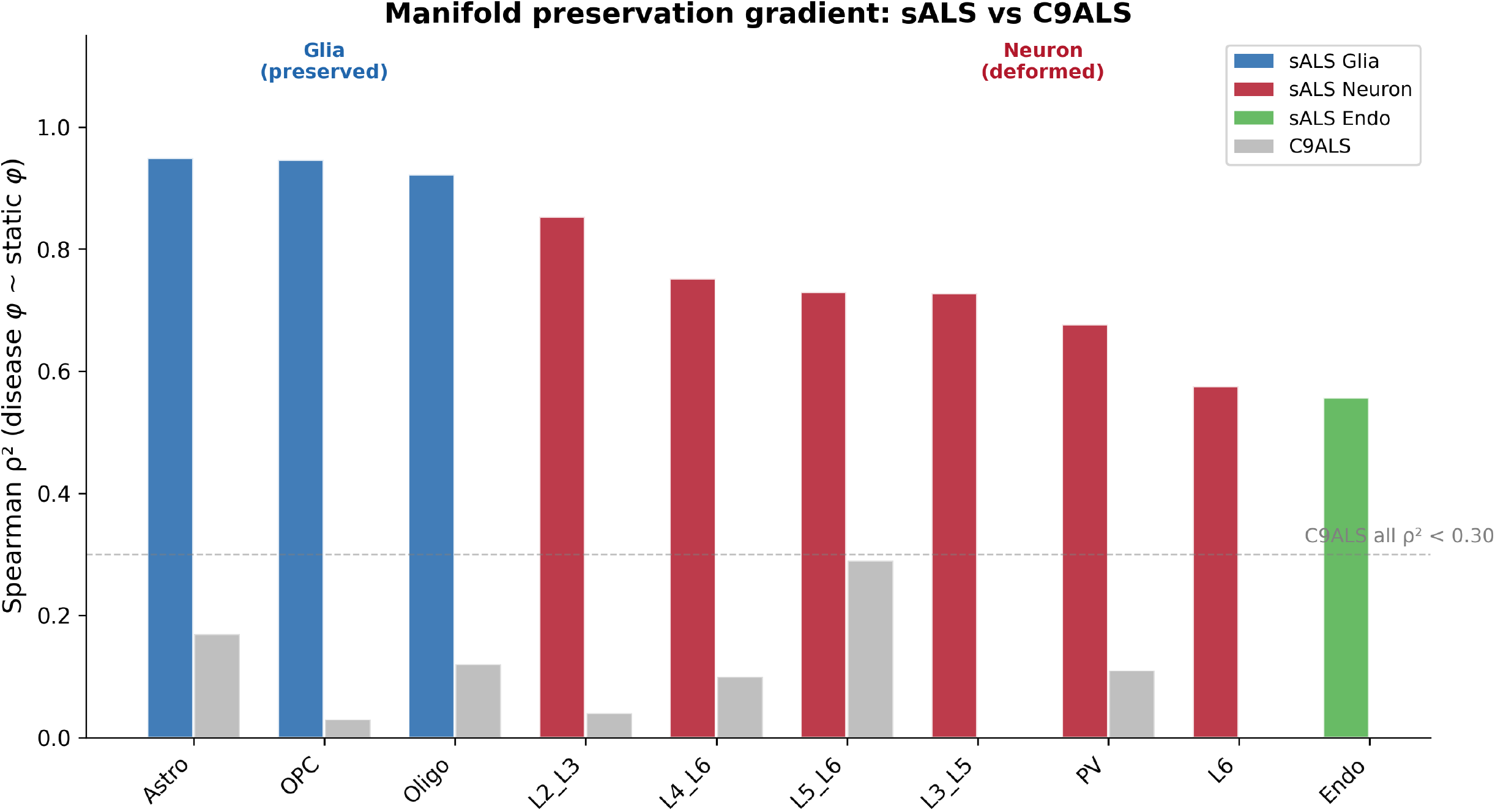
Cell-type gradient in manifold preservation distinguishes sALS from C9orf72-ALS. Bar plot of Spearman ρ^2^ between disease-state φ and static (PN-only) φ across all detected protein-coding genes for ten cell types, comparing sALS (coloured) and C9orf72-ALS (grey). sALS glial cell types (Astro 0.95, OPC 0.95, Oligo 0.92) largely retain the healthy manifold; sALS neuronal cell types (L2/3 0.85, L4/6 0.75, L5/6 0.73, L3/5 0.73, PV 0.68, L6 0.58) show progressive deformation; endothelial cells (0.56) occupy an intermediate position. In C9orf72-ALS, all cell types show ρ^2^ < 0.30 (horizontal dashed line), consistent with a uniform manifold-deformation architecture distinct from the topology-preserving mode of sALS.

**Figure 3.**
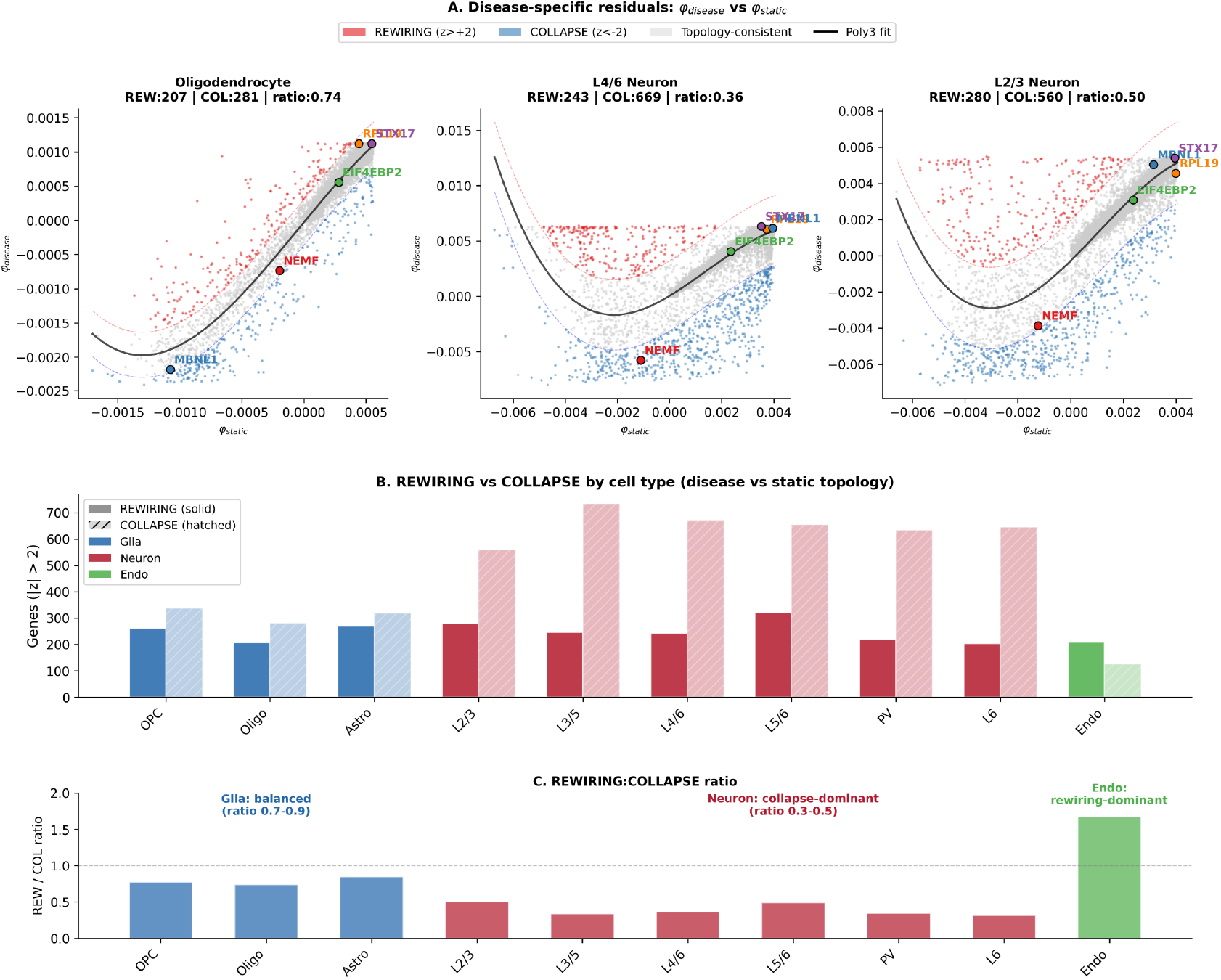
Rewiring versus collapse — cell-type-specific disease programmes revealed by 3φ residual analysis. (A) Scatter plots of disease-state φ versus static (PN-only) φ for all protein-coding genes, shown for three representative cell types (Oligo, L4/6, L2/3). The black curve is the cubic polynomial fit; red/blue dashed curves demarcate z > +2 (rewiring) and z < −2 (collapse) thresholds. NEMF (red), EIF4EBP2 (green), and representative topology-consistent genes (grey) are highlighted. (B) Rewiring (solid) and collapse (hatched) gene counts across all ten cell types. (C) Rewiring-to-collapse ratio: glia cluster at 0.7–0.9 (balanced); neurons at 0.3–0.5 (collapse-dominant); Endo alone exceeds 1.0 (1.67, rewiring-dominant). Horizontal dashed line at ratio = 1.0.

### 5. Translation/ribosome is the only cross-cell-type primary structural stress core

Among twelve candidate pathways tested with 3*φ* matched-null z-scores, translation/ribosome was the only programme satisfying the cross-cell-type universality criterion: ribosome subunits significantly upstream in 10/10 cell types and the broader translation class (RPL+RPS+EIF+EEF) in 9/10 cell types (Fig. 5). Four secondary pathways showed translation-independent significance in a subset of cell types: chaperones (4/10), ATP synthase (3/10), 26S proteasome (3/10), and ETC complex I (2/10). Seven pathways previously implicated in ALS pathology (TBK1/IKK→IRF signalling, nonsense-mediated decay, stress granule dynamics, autophagy core, calcium signalling, PI3K/AKT/mTOR, sphingolipid synthesis) failed to reach cross-cell-type significance under residual analysis (0/10 cell types), indicating that their apparent upstream enrichment in raw *φ* reflects co-expression-topology hub position rather than disease-specific perturbation.

Per-cell-type differential expression (PyDESeq2 ∼ condition + sex, sALS vs PN; Wald test) showed modest per-gene effects (|mean log_2_FC| < 0.4 in every cell type) but a highly significant coordinated upward shift of the translation gene class in most cell types under a one-sided Mann–Whitney rank test (the same GSEA-style approach used by Pineda et al. 2024; pooled-neuron p = 5.6 × 10^−21^; Table 3). The pooled-neuron top-decile enrichment was 2.14-fold (translation genes 21.4% of the top 10% genome-wide log_2_FC distribution versus the expected 10%).

**Table 3.**
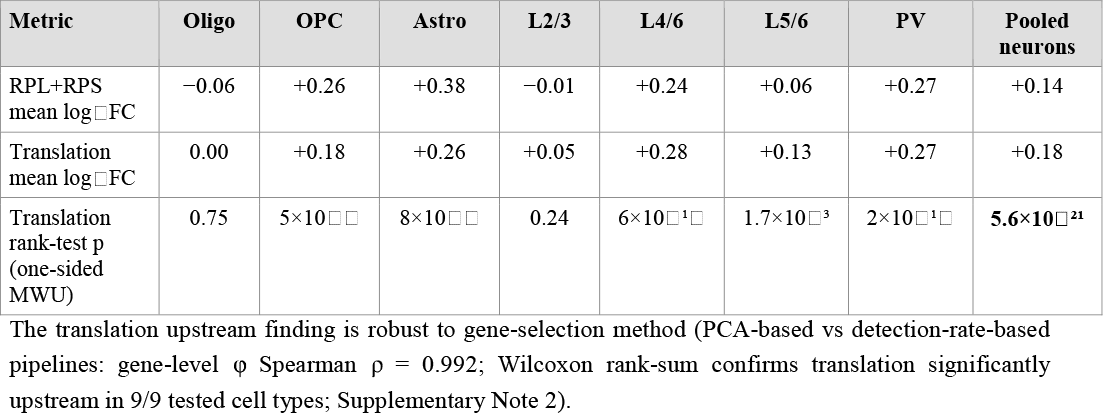
Cell-type-resolved differential expression of ribosomal/translation genes.

### 6. NEMF shows cross-cell-type disease-specific loss of co-expression coherence

At the single-gene level, the 3φ residual framework revealed that NEMF — the ribosome-associated quality-control factor that catalyses CATylation (C-terminal alanine-threonine tagging of nascent chains on stalled ribosomes) — was the only gene in the protein-coding genome satisfying z < −2 in ≥ 7 of 10 cell types in the NYGC cohort (Fig. 4). NEMF showed direction-consistent negative residuals in all ten cell types, passing the z < −2 threshold in seven (L3/L5, L5/L6, L6, Astro, L4/L6, OPC, PV). The strongest effect occurred in L3/L5 (z = −6.95), the cortical layer containing upper motor neurons in BA4 primary motor cortex; this z-score survives Bonferroni correction for the 8,874 detectable protein-coding genes (p_adj = 3.3 × 10 □ □). Under the binomial null with per-cell-type P(z < −2) = 0.0228, the probability of any gene satisfying z < −2 in ≥ 7 of 10 cell types by chance is 3.62 × 10 □^1^ □. Critically, NEMF mRNA expression was not significantly changed at the individual-cell-type level (sALS-only DESeq2 mean log □FC = −0.11, range −0.06 to −0.19; all p_adj > 0.05); the signal is the loss of co-expression coupling, not the expression level.

**Figure 4.**
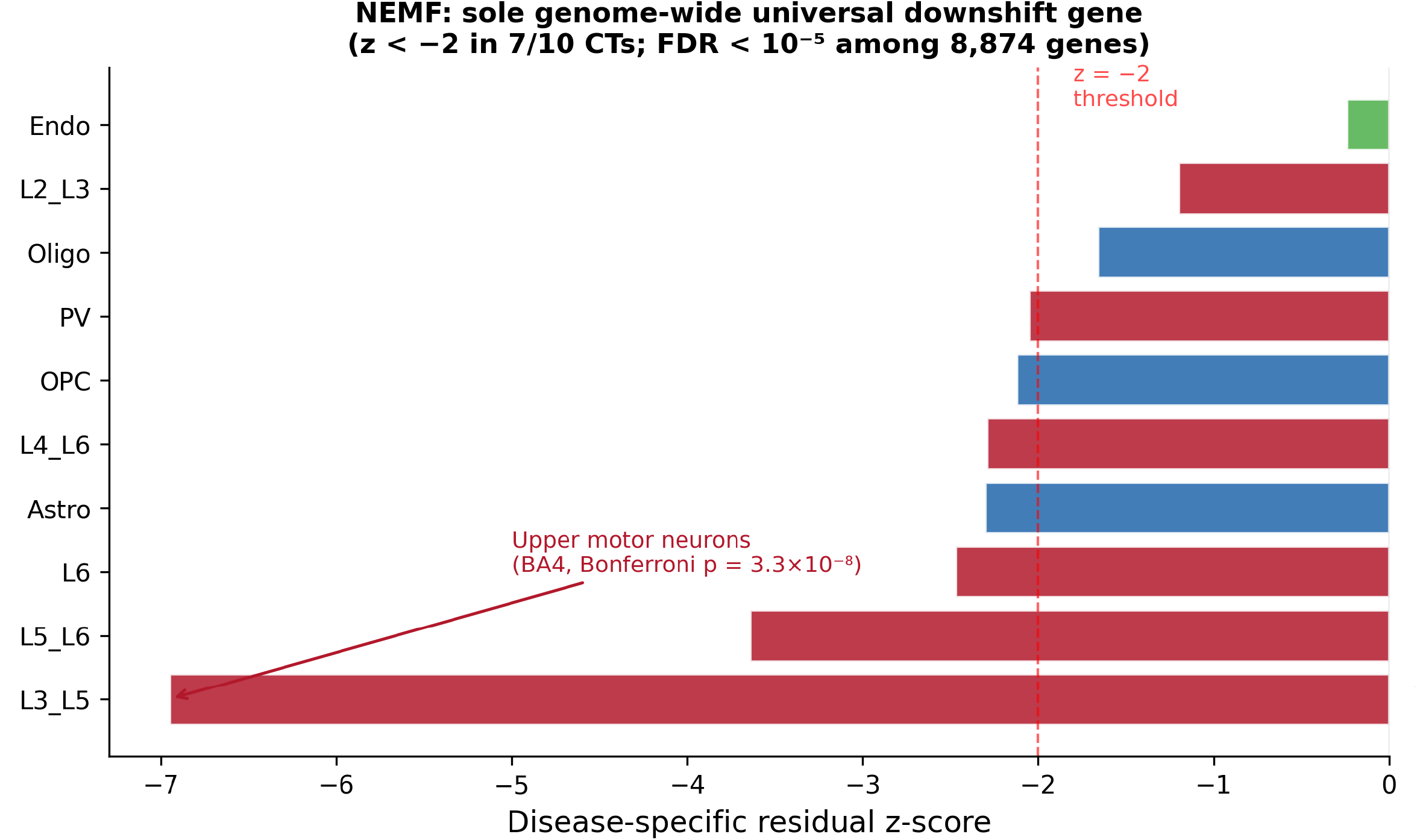
NEMF shows cross-cell-type disease-specific loss of co-expression coherence. Disease-specific residual z-scores for NEMF across ten cell types in the sALS cohort. NEMF shows direction-consistent negative residuals in all ten cell types, passing the z < −2 threshold in seven. The strongest effect is in L3/L5 (z = −6.95), the cortical layer containing upper motor neurons in BA4 (p_adj = 3.3 × 10[[after Bonferroni correction for 8,874 protein-coding genes). NEMF mRNA expression is not significantly changed at the individual-cell-type level (sALS-only DESeq2 mean log[FC = −0.11, range −0.06 to −0.19; all p_adj > 0.05); the signal is in the loss of co-expression coupling.

**Figure 5.**
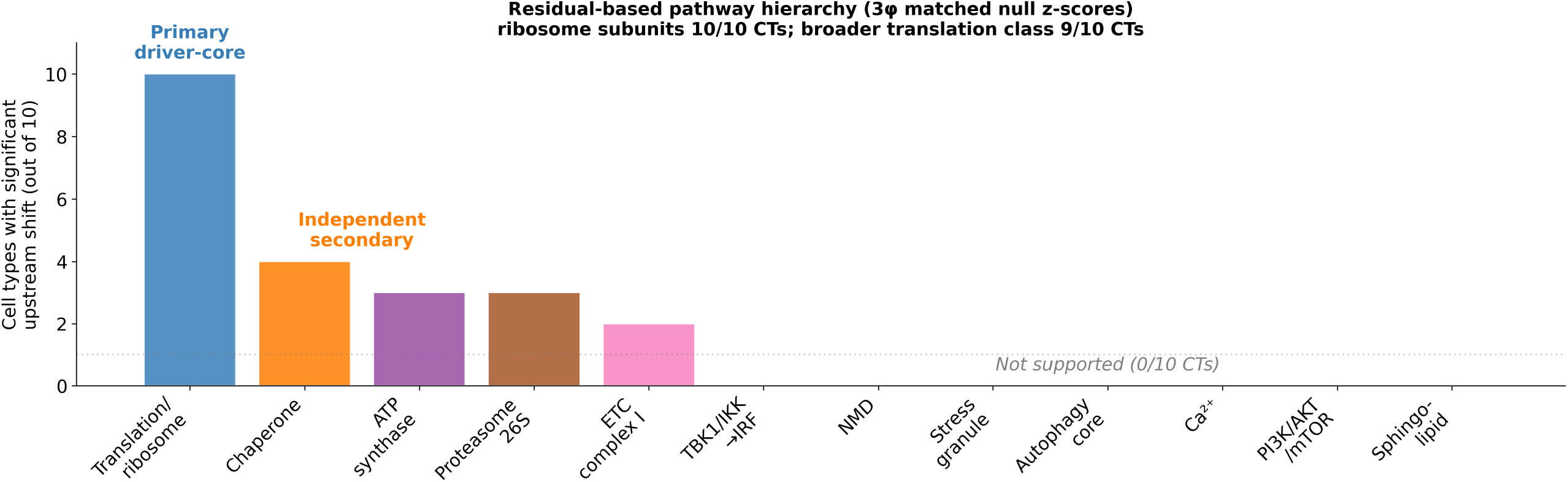
Residual-based pathway hierarchy identifies translation/ribosome as the sole cross-cell-type primary structural stress core. Number of cell types (out of 10) showing significant disease-specific upstream shift for twelve candidate pathways, computed via 3φ matched-null z-scores. Translation/ribosome (10/10 cell types; labelled “Primary driver-core” in the figure) is the only programme reaching the universality criterion; the broader translation class (RPL+RPS+EIF+EEF) reaches significance in 9/10 cell types under the same threshold. Secondary pathways with translation-independent significance (labelled “Independent secondary” in the figure): chaperones (4/10), ATP synthase (3/10), 26S proteasome (3/10), ETC complex I (2/10). Seven previously implicated pathways (TBK1/IKK→IRF, nonsense-mediated decay, stress granule dynamics, autophagy core, calcium signalling, PI3K/AKT/mTOR, sphingolipid synthesis) fail to reach significance under residual analysis (0/10 cell types each).

The cross-cell-type structure of NEMF residuals reveals a dual-axis architecture (Fig. 6). Genes co-collapsing with NEMF (positive ρ with NEMF residual z) are enriched for RNA-quality-control components and collision-response kinases — LSM14A (ρ = +0.94), DROSHA (ρ = +0.89), FXR1 (ρ = +0.89), PKR/EIF2AK2 (ρ = +0.73), NGLY1 (ρ = +0.83). Genes occupying compensatory positions (negative ρ) are enriched for translational suppressors and mitochondrial components — EIF4EBP2 (ρ = −0.99), MRPL39 (ρ = −0.89), COX6B1 (ρ = −0.86), MCUR1 (ρ = −0.84), ATP5F1D (ρ = −0.81). The cross-cell-type co-collapse between NEMF and PKR is consistent with impaired initiation of collision-responsive integrated-stress-response signalling, and the compensatory positioning of EIF4EBP2 and mitochondrial-translation / oxidative-phosphorylation genes is consistent with a translational-suppression / energetic-compensation programme triggered by CATylation failure.

**Figure 6.**
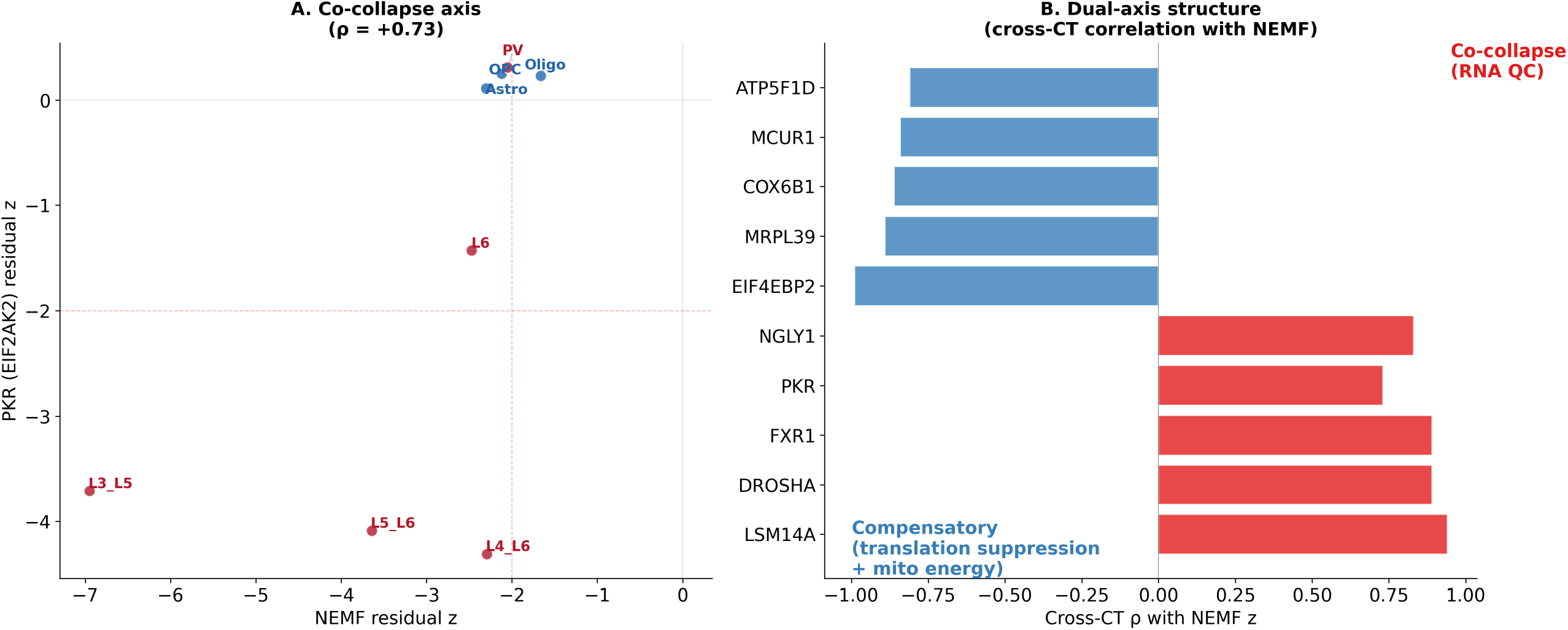
Dual-axis structure of the disease-specific response. (A) Cross-cell-type co-collapse between NEMF and PKR (EIF2AK2), a ribosome-collision-sensing kinase of the ISR. Red points denote the four neuronal cell types with NEMF residual z < −2 (L3/L5, L5/L6, L4/L6, L6); among these, three (L3/L5, L5/L6, L4/L6) additionally show PKR z < −2 and occupy the strict lower-left quadrant defined by the dashed thresholds. Blue points (PV, Oligo, OPC, Astro) cluster near the origin with preserved coupling. Positive cross-cell-type correlation ρ = +0.73 (n = 8 cell types for which both z-scores are defined). (B) Dual-axis structure from cross-cell-type correlation of NEMF residual z-scores with all other genes. Red bars (co-collapse axis; positive ρ with NEMF z) enrich for RNA quality control (LSM14A ρ = +0.94, DROSHA ρ = +0.89, FXR1 ρ = +0.89, PKR ρ = +0.73, NGLY1 ρ = +0.83). Blue bars (compensatory axis; negative ρ) enrich for translational suppressors and mitochondrial components (EIF4EBP2 ρ = −0.99, MRPL39 ρ = −0.89, COX6B1 ρ = −0.86, MCUR1 ρ = −0.84, ATP5F1D ρ = −0.81). Panel B reports cross-cell-type co-expression coupling correlations, not expression-level effects.

These observations converge with genetic evidence. ENU-derived mouse Nemf mutations that selectively impair CATylation cause progressive motor neuron degeneration (Martin et al. 2020), and NEMF-mutant mice develop TDP-43 proteinopathy through CATylation failure leading to truncated peptide accumulation, Importin-β pathological interaction, nucleocytoplasmic transport defects, and ultimately TDP-43 nuclear loss with cytoplasmic phosphorylation (Plessis-Belair et al. 2024). The selective disruption of NEMF co-expression coherence in sporadic ALS — where no NEMF coding mutations are present (Hop et al. 2026: NEMF exome p = 0.53) — suggests that the same CATylation bottleneck can arise through non-genetic mechanisms, providing a candidate explanation for the ∼90% of ALS cases that show TDP-43 proteinopathy without identifiable mutations in TDP-43 or known RQC components.

### 7. Cross-cohort validation in three independent BA4/MCX cohorts (N = 107)

We applied the same SPD/log-Euclidean Hodge operator family to three independent BA4/MCX sALS snRNA-seq cohorts: the NYGC discovery cohort (33 donors: 17 sALS + 16 PN after exclusions); Takeuchi et al. 2025 (Brain awaf426; 12 donors: 6 sALS + 6 controls, 10x Multiome); and Ruf et al. 2026 (Nat Commun; 62 donors: 30 sALS + 32 PN, 10x Multiome with WNN/scVI/peakVI integration). The total N = 107 donors after exclusion of C9ALS/FTLD cases. A pre-specified merged-pool diffusion-map pseudotime (PT-B) failed pre-specified direction and effect-size thresholds in this 3-cohort pool (Spearman ρ(PT-B, disease) = +0.116; Cramér’s V cohort × PT-B-quartile = 0.65), indicating that cohort manifold geometry dominates over the biological gradient at the merged level. Per the pre-specified fallback protocol, primary analysis pivoted to condition-displacement φ, computed as the Frobenius norm of the cohort-baseline-zeroed tangent representation difference between ALS and CTRL donor means, decomposed via Hodge on the 7-cell-type complete graph.

A brain-origin, circuit-distal architecture (*φ*_Oligo_ > *φ*_IT_ > *φ*_ET_, with *φ*_Oligo_ > 0 and *φ*_ET_ ≤ 0) emerged in all three cohorts (Table 4). After matched-donor B = 1000 bootstrap hardening, P(brain-origin) ≥ 0.950 in each cohort and ≥ 0.999 across the joint NYGC + Ruf2026 ET-focused bootstrap (Takeuchi-independent, n_ALS_ = 36 / n_CTRL_ = 36; *φ*_ET_ 95% CI [−1.057, −0.476]).

**Table 4.** Per-cohort condition φ across 7 common cell types (N = 107).

| Cohort | $\phi_{\text{Oligo}}$ | $\phi_{\text{OPC}}$ | $\phi_{\text{Astro}}$ | $\phi_{\text{Micro}}$ | $\phi_{\text{IT}}$ | $\phi_{\text{ET}}$ | $\phi_{\text{GABA}}$ | Brain-origin? |
| --- | --- | --- | --- | --- | --- | --- | --- | --- |
| NYGC | **+0.605** | +0.150 | -0.282 | +0.604 | +0.017 | **-1.137** | -0.001 | □ |
| Takeuchi 2025 | **+1.340** | -0.190 | -0.107 | +0.395 | +0.153 | **-1.662** | +0.071 | □ |
| Ruf 2026 | **+0.357** | -0.054 | -0.097 | +0.151 | -0.189 | **-0.322** | +0.142 | □ |

Three independent cell-type labelling schemes — cohort-author labels, T-core canonical broad-CT markers, and T-NYGC subtype-marker projection — gave concordant results for the cardinal Oligo+ET architecture (3/3 cohorts in all three schemes; ET-focused stratified bootstrap $P(\phi_{\text{Oligo}} > 0) = 1.000$ throughout). Microglia $\phi$ sign was taxonomy-sensitive (cohort-author 3/3 positive, T-core 0/3 positive, T-NYGC-FULL 1/3 positive) and is therefore not retained as a cross-cohort finding; the cardinal Oligo+ET pattern does not depend on Microglia. The NEMF gene-level finding was tested with a calibration-matched matched-null $z$ framework in each cohort. NEMF showed predominantly negative direction across all three cohorts and breadth replication ( $z < -2$ in $\geq 7/10$ cell types) in NYGC and Ruf2026 (two-cohort breadth, three-cohort direction). The strongest cross-cohort residual signal therefore generalises beyond the discovery cohort under independent CT-labelling schemes, supporting NEMF/CATylation disruption as a candidate molecular intermediate rather than a cohort-specific artefact.

## Discussion

The main methodological finding of this study is that perturbation-source information is not contained in static covariance itself, but becomes recoverable in the control-normalised covariance-response field. Norman Perturb-seq bridges interventionist and structural causality: genes that are causal sources by CRISPR intervention are recovered as structural sources by off-diagonal covariance-response aggregation. On complete graphs this readout is shared by Hodge φ and classical centrality measures applied to the same whitened Δ matrix; on sparse k-NN graphs Hodge gains a regime-dependent advantage through Laplacian smoothing. The Hodge framework therefore functions not as a uniquely superior complete-graph ranking heuristic, but as the field-theoretic representation that gives covariance-response centrality source/sink semantics and enables gradient/curl decomposition, multi-transition integration, and 3φ disease-residualisation against healthy co-expression topology. External benchmarks anchor validity at multiple layers — temporal recovery in TimeVault, intervention-source recovery in Norman Perturb-seq, functional-importance recovery in spatial CRISPR, and label-free clinical-staging recapitulation in cross-disease glioma (ρ = −0.755 with grade) — and the same readout, applied to sALS across three independent BA4/MCX cohorts (N = 107), yields four convergent observational findings discussed below.

The reasoning underlying this thesis followed a non-obvious arc. Hodge decomposition was initially adopted to infer causal direction from covariance dynamics, but application to sALS revealed that on the complete graph, Hodge φ is rank-equivalent to the row-engagement scalar d_i and to the broader structural-centrality family (L1, PageRank, eigenvector; Supplementary Note 6). Re-verification on Norman Perturb-seq confirmed that experimentally imposed CRISPR sources are recovered by any family member — locating causal-source information in the hub-centrality structure of the covariance-response field itself, with Hodge supplying source/sink semantics, sparse-graph Laplacian recovery, and 3φ disease residualisation.

### φ measures co-expression topology by default and disease information through its residuals

On the complete graph K_N_, *φ* is practically equivalent to a monotone transformation of the row-engagement scalar d_i_ (Spearman ρ ≈ −1.000), and 82–92% of *φ* variance in glia reflects the weighted strength of each gene in the healthy co-expression network. The biological signal therefore does not sit in raw *φ* position but in the residuals after topology regression — the 3*φ* framework. This separation is itself biologically informative: in glia, sALS operates largely within the existing co-expression manifold (R^2^ = 0.92–0.95); in neurons, the manifold itself is deformed (R^2^ = 0.58–0.85); endothelial cells exhibit rewiring-dominated architecture (ratio 1.67). These three modes — manifold-preserving with disease-specific rewiring of the translational core (oligodendrocytes), collapse-dominant manifold deformation (neurons), and rewiring-dominant network construction (endothelium) — are reproducible at the gene level across the cohort and define the architectural context for the molecular findings. The same comparison in C9ALS collapses the glia-neuron gradient (all R^2^ < 0.30), demonstrating that the topology-preservation gradient is sALS-specific rather than a generic feature of ALS.

### Translation/ribosome is the only structural stress core that universally signals across cell types

Among the twelve pathways tested, only ribosome subunits (10/10) and the broader translation class (9/10) reach cross-cell-type significance under residual analysis. Seven pathways previously implicated in ALS pathology — TBK1/IKK signalling, NMD, stress granule dynamics, autophagy, calcium signalling, PI3K/AKT/mTOR, sphingolipid synthesis — reach high raw-*φ* positions but fail residual analysis, indicating that their apparent upstream-ness reflects network-hub topology rather than disease-specific perturbation. This is a methodologically important distinction: pathway-level claims in single-snapshot data must contend with the fact that highly connected nodes are upstream by topology in nearly every co-expression network, and a residual or null-controlled analysis is required to disentangle topology from disease signal. The coordinated upward shift of the translation class in differential expression (pooled-neuron one-sided MWU p = 5.6 × 10^−21^) and the 2.14-fold top-decile enrichment further support that the translation signal is a coordinated set-level shift, not a topology artefact. Translation upstream-ness is therefore the only programme in our analysis with strong evidence for being both structurally upstream and disease-specific across cell types.

### NEMF/CATylation network disruption is the strongest disease-specific single-gene signal

NEMF was the sole gene in the protein-coding genome with disease-specific co-expression-coherence loss in ≥ 7 of 10 cell types in the discovery cohort (binomial p = 3.62 × 10^−10^), unchanged in mRNA expression, with the strongest residual in L3/L5 (z = −6.95), the cortical layer containing upper motor neurons. The cross-cell-type co-collapse with the ribosome-collision sensor PKR and the compensatory positioning of EIF4EBP2 and mitochondrial-translation / oxidative-phosphorylation components produce a dual-axis structure interpretable as collision-response initiation failure coupled with a translational-suppression / mitochondrial-energy compensatory programme. Mouse Nemf mutations selectively impairing CATylation cause progressive motor neuron degeneration (Martin et al. 2020) and TDP-43 proteinopathy through truncated-peptide-driven Importin-β sequestration and nucleocytoplasmic-transport defects (Plessis-Belair et al. 2024); the absence of NEMF coding mutations in sporadic ALS (Hop et al. 2026) raises the possibility that the same CATylation bottleneck arises through non-genetic disruption of NEMF’s regulatory neighbourhood. The NEMF gene-level finding cross-validates in 2/3 cohorts at the ≥ 7/10-CT breadth criterion and shows direction-consistency in 3/3 cohorts.

### Cross-cohort generalisability and the brain-origin pattern

The condition-displacement *φ* architecture — *φ*_Oligo_ > 0 with *φ*_ET_ < 0 — is bootstrap-supported (P ≥ 0.95) in 3/3 cohorts and ≥ 0.999 in the Takeuchi-independent NYGC + Ruf2026 stratified bootstrap. The pattern is robust to three independent cell-type labelling schemes. This central preservation in oligodendrocytes combined with extratelencephalic / upper-motor-neuron-containing layer sink behaviour supports a brain-origin, circuit-distal architecture in which structural pathology originates centrally and glially, producing a clinical phenotype that resembles classical dying-back (in which motor neuron loss appears to initiate at distal axons and propagate retrogradely) yet originates centrally. Microglia did not generalise across taxonomies and is therefore not retained as a cross-cohort finding.

### Limitations

Raw φ ranking on K_N is not a Hodge-specific causal estimator; it lies in the same structural-centrality family as L1 strength, PageRank, and eigenvector centrality on the whitened Δ matrix (Supplementary Note 6). This equivalence is part of the finding: covariance-response geometry contains source information, and Hodge licenses its source/sink interpretation by virtue of the K_N rank equivalence with these classical centrality measures (closely related to the HodgeRank framework; Jiang et al. 2011). Hodge-specific ranking advantage appears in the sparse-graph regime through Laplacian smoothing (+18.6 pp over raw signed divergence at k = 10, strict dominance), where divergence aggregated over few k-NN edges is locally noisy and benefits from global random-walk-like averaging. Disease-specific biological interpretation requires the 3φ topology-residual framework rather than raw φ position. The method performs intervention-consistent causal-source recovery under benchmark perturbations, not Pearl/Rubin causal-effect estimation.

The instrument does not estimate causal effects in the Pearl/Rubin sense. Raw φ on K_N primarily reflects co-expression topology; disease-specific information resides in residuals. The CRISPR validation confirms that φ-upstream genes are functionally important when removed, but cannot be separated from a generic hub-removal effect; disease-specific upstream-ness is better supported by the rewiring residual analysis. Roughly 34% of perturbation groups fail at Top-1 identification in Norman Perturb-seq; diffuse, indirect, non-transcriptional, and epistatically masked effectors fall outside the instrument’s scope. The Part I sALS discovery analyses derive from a single primary cohort (24 donors), with the cross-cohort meta-analysis partially addressing this at the cell-type-level condition-displacement architecture and at the gene-level NEMF finding; independent replication of the gradient-curl separation itself remains essential. The pseudotime-based meta-analysis was not feasible for the 3-cohort merged pool (cohort manifold dominates over biological gradient), restricting the cross-cohort claim to condition-displacement φ. All three meta-cohorts are BA4/MCX cortex from autopsy, so the cross-cohort claim does not extend to spinal cord motor neuron pathology. Sphingolipid biosynthesis in oligodendrocytes — masked by PCA-based gene selection — emerges as an additional upstream programme under detection-rate selection (Supplementary Note 2); the complete upstream landscape may therefore be broader than the present pipeline captures. Two-sample Mendelian randomisation on oligodendrocyte cis-eQTLs and ALS GWAS was non-significant (IVW OR = 0.988, p = 0.159), reflecting a power-limited null (minimum detectable OR = 1.012); Bayesian colocalisation identified KANSL1 at chr17q21.31 (PP.H4 = 0.794). The pipeline identifies translation as the structurally upstream programme but cannot, from observational data alone, distinguish causal disease involvement from stable housekeeping behaviour; targeted perturbation of translation-related genes in ALS-relevant cell types is required to definitively resolve this distinction.

### Hypothesis and outlook

The observations support a candidate cascade in which oligodendrocyte translational stress (preserved-topology rewiring of the ribosomal core), motor-neuron NEMF/CATylation network disruption (collision-response initiation failure with compensatory translational suppression), and downstream TDP-43 pathology occupy successive positions in a brain-origin architecture. Falsifiable predictions include: (i) immunohistochemistry for NEMF in sALS post-mortem spinal cord motor neurons should reveal altered expression or subcellular localisation, preceding or co-occurring with TDP-43 pathology; (ii) ribosome stalling reporter assays in sALS patient-derived iPSC motor neurons should show reduced CATylation efficiency without NEMF coding mutations; (iii) AAV-mediated NEMF overexpression in NEMF-mutant mice should rescue TDP-43 nuclear loss if CATylation is the rate-limiting step; (iv) ISR-kinase protein levels in sALS motor cortex neurons should show selective reduction of GCN2 and PKR with PERK preserved, mirroring the co-expression pattern. A parallel observation — early ATM-pathway decline in capillary endothelium identified by an independent module-level causal pipeline applied to the same NYGC dataset — and a tentative integrative cascade model connecting NVU trigger, R-loop accumulation, ribosome-stalling, RQC overload, and TDP-43 convergence are presented in Supplementary Discussion S1–S2; they constitute speculative working hypotheses, not established mechanisms, and are independent of the empirical findings reported in the main text.

## Methods

*Concise overview only; full methodological detail, including stage-by-stage computational specifications, the axiomatic justification of the edge-weight flow construction, the analytical derivation of the GF*□ *null baseline, and Lipschitz-type stability bounds, is provided in Supplementary Methods and Supplementary Notes 1–7*.

### Data

Discovery cohort: NYGC ALS Consortium snRNA-seq, primary motor cortex (BA4). The full MCX panel comprises 17 sALS + 16 PN + 16 C9ALS + FTLD donors. The main Hodge pipeline (cell-type ranking, two-axis model, multi-transition integration, pseudotime construction) uses a 14 sALS + 10 PN subset after joint cell-count QC (≥ 100 cells per donor in each of the top four cell types). The 3φ residual framework and sALS-only differential expression use the full 17 sALS + 16 PN pool with a per-CT ≥ 20 cells per donor filter. The sALS-vs-C9ALS condition-φ comparison uses 16 C9ALS donors paired with the same 10 PN donors. Two complementary 10-cell-type sets are used (nine shared core types plus 5HT3aR or Endo, depending on the per-pipeline sample-size constraints); full rationale and per-CT donor counts in Supplementary Methods. Validation cohorts: Takeuchi et al. 2025 (n = 12; doi:10.1093/brain/awaf426) and Ruf et al. 2026 (n = 62; doi:10.1038/s41467-026-69944-6), both BA4/MCX 10x Multiome. Benchmark datasets: Norman Perturb-seq (scPerturb, 111,122 K562 cells); spatial CRISPR (Shen et al. 2026, GSE274058); TCGA + GTEx glioma (TOIL pipeline, 723 samples); TimeVault (Chao et al. 2026, 12 bulk samples). Preprocessing: log □ □ (CPM+1); OLS regression of donor, nUMI, and percent_mito covariates before SPD covariance computation.

### Five-stage Hodge pipeline

(1) Per-donor PCA (k = 30), Ledoit–Wolf shrinkage covariance, log-Euclidean mapping log(C) = Q diag(log λ_i_) Q^T^. (2) Diffusion pseudotime (PT-B) on the inter-donor log-Euclidean distance matrix; an expression-based PT-A is used for sensitivity analysis (cross-pipeline gene-level *φ* Spearman ρ = 0.975). The 24 donors are partitioned into 6 windows of 4 donors each. (3) Delta matrix Δ_t_ = mean(log C_wt+1_) − mean(log C_wt_); edge-weight flow f(i,j) = |Δ_ij_| × sign(d_i_ − d_j_) with d_i_ = □ Δ_i,·_□_2_. (4) Hodge decomposition on K_N_ via φ = L^+^ div(f), where L^+^ = (1/N)(I − J/N); GF = □ grad □^2^/□f □^2^. On k-NN sparse graphs, φ is computed via LSQR; curl per triangle Γ_ijk_ = f(i,j) + f(j,k) − f(i,k) followed by Louvain community detection. (5) Stable-High genes are top quantile in ≥ 4/5 transitions across 100 bootstrap iterations. Axiomatic justification of the edge-weight flow construction (Supplementary Note 3); closed-form GF_0_ = 2/[3(1+CV^2^)] baseline derivation (Supplementary Note 4); Lipschitz-type stability bound for the LOCO rank-determining quantity (Supplementary Note 1). Natural-gradient interpretation: under the log-Euclidean metric on SPD(n), for a differentiable stress functional *** on the tangent coordinate X = log Σ, the consecutive-window displacement Δ_w = X_(w+1) − X_w = −η_w □[**** (X_w) + O(η_w^2^) is the first-order observable tangent displacement of a stress-response dynamics, so the edge-weight flow constructed from Δ inherits source/sink semantics under the Hodge decomposition (full derivation, axiomatic justification, and conditional source-identifiability statement in Supplementary Note 7).

### 3φ residual framework

φ_static (10 PN donors, zero disease information), φ_disease (14 sALS donors), and φ_condition (pooled 24-donor panel) are computed by the same five-stage pipeline. Per cell type, φ_disease is regressed on φ_static via cubic polynomial fit (numpy.polyfit, degree 3) in raw-φ space; residuals are standardised per cell type. Matched null z-scores are computed by 1,000 ALS/PN label permutations and used for the pathway-and gene-level upstream tests. Genes with z > +2 are classified as rewiring; z < −2 as collapse.

### Whitening for perturbation data

W = V D^{−1/2} V^T from Ledoit–Wolf Σ_ctrl (eigenvalues clamped at 10 □^1^□); bootstrap B = 100. Whitening is not applied in the sALS pipeline because the pseudotime delta construction serves an analogous role.

Variance decomposition and structural-family benchmarks. To distinguish marginal-variance, whitening, and off-diagonal covariance-response components of perturbation-source recovery, eight scorers were computed on the same Norman test split, candidate-gene universe, control whitening, and bootstrap protocol (n = 100 bootstrap iterations per perturbation group, seed = 42, control subsample = min(max(n_pert_, 100), n_ctrl_)). Diagonal scorers: raw per-gene variance ratio |log(var_pert_ / var_ctrl_)|, per-gene scale-corrected variance (diagonal z), ZCA-whitened per-coordinate variance, and control-conditioned residual variance via Schur complement on the control precision matrix (r_j_[cell] = (X[cell] · Ω_c_)[j] / Ω_c_[j,j]; preserves gene identity). Structural-family scorers on the whitened SPD log-correlation Δ matrix: L2 row norm (d_corr_), L1 row sum, PageRank (α = 0.85, power iteration), eigenvector centrality (top eigenvector of |Δ|), and Hodge φ. Sparse-graph variants (k = 10, 20, 50, 100), signed-divergence controls (Hodge φ vs raw divergence on the same k-NN graph), and per-pair McNemar / Spearman ρ analyses are described in Supplementary Note 6. The benchmark establishes that Top-1 source identification specifically requires off-diagonal covariance-response aggregation, with Hodge-specific Laplacian-smoothing advantage emerging in the sparse-graph regime.

### Differential expression and pathway enrichment

Per-cell-type pseudobulk PyDESeq2 0.5.0 with Wald test on ∼ condition + sex (sALS vs PN), apeglm shrinkage, BH-FDR; per-CT filter ≥ 20 cells per donor and ≥ 1 count in ≥ 3 donors. GSEA-style rank test: one-sided Mann–Whitney U on transcriptome-wide ranked log□FC for the Translation gene class (RPL+RPS+EIF+EEF) versus all other genes per cell type; pooled-neuron analysis concatenates L2/3, L4/6, L5/6, PV (n = 543 Translation genes). Pathway enrichment of rewiring genes (z > +1): Metascape (Zhou et al. 2019; GO BP, KEGG, Reactome, CORUM; min overlap 3, min enrichment 1.5; BH-adjusted). Preranked GSEA on φ-ranked lists via GSEApy/Enrichr (Fang et al. 2023). DisGeNET cross-reference via the Metascape module.

### Cross-cohort meta-analysis

Seven broad common cell types (Oligo, OPC, Astro, Micro, IT_neurons, ET_neurons, GABAergic_neurons). Per-cohort residualisation (log-CPM, OLS on donor + nUMI + percent_mito), joint PCA (k = 80) on cohort-concatenated residual matrix, per-donor Ledoit–Wolf SPD covariance, and Option C cohort-baseline subtraction in log-Euclidean tangent space (V_{d,c,t} = log Σ_{d,c,t} − log Σ□^{ctrl}_{c,t}). Condition-displacement φ: drive_t(c) = □mean(V_ALS) − mean(V_CTRL) □_F; flow_{i,j}(c) = drive_i(c) − drive_j(c) on K_7; φ_t(c) = mean(drive) − drive_t (φ > 0 = preserved; φ < 0 = sink-like). Three independent CT-labelling schemes (cohort-author, T-core canonical markers, T-NYGC-FULL subtype-marker projection) re-execute the full pipeline; matched-donor B = 1000 bootstrap hardening. The pre-specified fallback protocol when PT-B fails direction/effect-size thresholds and the locked decision rules are described in Supplementary Methods.

### Mendelian randomisation

Two-sample MR using oligodendrocyte cis-eQTLs (Bryois et al. 2022, n = 196) as instruments and ALS GWAS (van Rheenen et al. 2021, N_eff = 80,713) as outcome. Instruments selected at p < 5 × 10 □□, LD-clumped (r^2^ < 0.001, 10 Mb window, EUR reference), F ≥ 10. Four methods (IVW, MR-Egger, weighted median, weighted mode); sensitivity by Egger intercept, Cochran’s Q, leave-one-out. Bayesian colocalisation via coloc.abf (Giambartolomei et al. 2014), ±500 kb, top 100 genes.

### Robustness validations

Leave-one-donor-out (LOCO, 24 iterations): cross-run Spearman ρ = 0.981 on gene-level φ rankings (mean across cell types). PT-A vs PT-B: gene-level Spearman ρ = 0.975. PCA-based vs detection-rate-based gene selection: ρ = 0.992; Wilcoxon translation upstream in 9/9 cell types. Cell-type-level leave-one-donor-out bootstrap (LODO): 24/24 iterations retained the top-ranked oligodendrocyte position.

### Software and code availability

Python ≥ 3.10, PyTorch, anndata, scanpy, scikit-learn, scipy, PyDESeq2 0.5.0. Code: https://github.com/akacola2006/scrnaseq-hodge-pipeline (commit 10d60e7, 2026-04-26; a tagged release and Zenodo DOI will be deposited upon manuscript submission).

## Ethics statement

This study used publicly available, de-identified secondary human transcriptomic datasets. Ethical approval and informed consent were obtained by the original data-generating studies, as described in the corresponding publications. No new human participants were recruited and no new identifiable human data were generated in this study.

## Data availability

NYGC ALS Consortium data as processed by Pineda et al. (2024); BioProject PRJNA1073234; Synapse syn51105515. Only MCX (BA4) data were used; DLPFC (BA9) data from the same cohort were not included. Three donor subsets are used across the analysis pipelines (full per-pipeline criteria in Supplementary Methods). Cross-dataset validation: Wang H.V. et al. (2025) GSE212630 (C9orf72 ALS/FTD dlPFC). Norman Perturb-seq via scPerturb; CRISPR data Shen et al. 2026 GSE274058. Glioma data: TCGA glioma and GTEx via UCSC Xena TOIL pipeline. TimeVault: Chao et al. 2026. ALS exome statistics: Hop et al. 2026 Supplementary Tables MOESM6/7/10. Takeuchi 2025 (doi:10.1093/brain/awaf426) and Ruf 2026 (doi:10.1038/s41467-026-69944-6) accessed via the original deposition records.

## Code availability

The complete analysis pipeline is available at https://github.com/akacola2006/scrnaseq-hodge-pipeline (commit 10d60e7, 2026-04-26). The repository includes the core Hodge decomposition pipeline, Gene Hodge analysis scripts, WGCNA integration, all-gene insertion ranking, condition φ analysis, and reproduction workflows for all main figures and tables.

## Acknowledgements

Data used in the preparation of this article were obtained from the NYGC ALS Consortium (Synapse syn51105515; Pineda et al., Cell 2024). We thank the NYGC ALS Consortium, the donors, and their families for making these data publicly available. The results published here are in part based upon data generated by the TCGA Research Network; GTEx data were obtained from the GTEx Portal. The authors used AI language tools (Claude, Anthropic; Gemini, Google DeepMind; ChatGPT, OpenAI) to assist with manuscript drafting and editing, mathematical notation, and code development. All analyses were designed, executed, and verified by the authors, who take full responsibility for the integrity and accuracy of the work.

## Author contributions

S.K. conceived the framework, designed the pipeline, performed all analyses, and wrote the manuscript.

M.U. provided clinical expertise and supervision.

## Competing interests

The authors declare no competing interests.

## Funding

This work was supported by the Japan Agency for Medical Research and Development (AMED) under Grant Number JP[26wm0625522h0002] (Brain/MINDS 2.0: Multidisciplinary Frontier Brain and Neuroscience Discoveries), and JP[26bk0104169s0703] (Practical Research Project for Rare/Intractable Diseases), Health, Labour and Welfare Sciences Research Grants from the Ministry of Health, Labour and Welfare of Japan (MHLW) under Grant Number [26FC1008] (Research Project on Rare/Intractable Disease), and by KAKENHI (Kiban B, 23K24211) to M.U.

## Notes

https://github.com/akacola2006/scrnaseq-hodge-pipeline.

### Competing Interest Statement

The authors have declared no competing interest.

### Summary of Updates

The abstract shown on the landing page reflected an early version of the manuscript and had not been updated together with subsequent file revisions. This revision updates the displayed abstract to match the current version of the PDF. No further changes to the manuscript.

