## supplementary text for "Intervention-consistent causal-source recovery from covariance-response geometry reveals upstream organisation in sporadic ALS"

Kaneko S.^1^, Urushitani M.^1^

*1Department of Neurology, Shiga University of Medical Science, Otsu, Japan*

### Contents

- Supplementary Methods (S1–S19)
- Supplementary Note 1 — Theoretical Background for the Stress-Covariance Framework
- Supplementary Note 2 — PCA Selection Sensitivity Analysis
- Supplementary Note 3 — Axiomatic Justification of the Edge-Weight Flow Construction
- Supplementary Note 4 — Analytical Derivation of the Gradient Fraction Null Baseline GF₀
- Supplementary Note 5 — Stage-by-Stage Computational Details of the IDS Pipeline
- Supplementary Note 6 — Norman Perturb-seq Covariance-Response Source-Recovery Audits
- Supplementary Note 7 — Structural causal upstreamness from covariance-response geometry
- Supplementary Discussion S1 — ATM Brake Loss and the NVU Trigger Hypothesis
- Supplementary Discussion S2 — Hypothetical Integration: A Multi-Layer Cascade Model for sALS

### Supplementary Methods

The Methods overview in the main text summarises the computational pipeline at the level required for general readers; this Supplementary Methods section provides the implementation-level detail required for reproduction. Subsection numbering (S1–S19) corresponds one-to-one to the original Methods sections 6.1–6.19 used during manuscript preparation. Theoretical background for the stress-covariance framework (Supplementary Note 1), gene-selection sensitivity (Supplementary Note 2), the axiomatic justification of the edge-weight flow rule (Supplementary Note 3), the closed-form gradient-fraction null derivation (Supplementary Note 4), and the unified stage-by-stage computational reference (Supplementary Note 5) are provided in the Supplementary Notes section that follows.

#### S1. Data (original §6.1)

**sALS snRNA-seq.** NYGC ALS Consortium (Pineda et al. 2024), primary motor cortex (MCX, Brodmann area 4). The NYGC cohort includes both MCX (BA4) and DLPFC (BA9); only MCX data were used in this pipeline. The full NYGC MCX cohort comprises 17 sporadic ALS (sALS) and 16 pathologically normal (PN) donors, plus C9ALS and FTLD donors that are used separately (16 C9ALS donors are used for the condition φ comparison in Results §4).

Three different donor subsets are used across three stages of the analysis, each matched to the statistical requirements of its pipeline. (i) The main Hodge pipeline (Part I cell type φ ranking, two-axis model, multi-transition integration, and pseudotime construction) uses a reduced 14 sALS + 10 PN subset after applying stringent joint cell-count QC that requires ≥ 100 cells per donor in each of the top four cell types used for pseudotime construction; three sALS donors (IDs 102, 112, 124) and six PN donors (IDs 317-319, 322-324) were excluded at this step because they fell below the per-donor-per-cell-type cell count threshold in at least one of the top-four cell types. (ii) The 3φ residual framework (Sections 4.1, 4.2, 8.9, and Fig. 3) and the sALS-only differential expression analyses (Methods (Differential expression and pathway enrichment)) use the full 17 sALS + 16 PN pool subject to a less stringent per-CT filter of ≥ 20 cells per donor, which reduces the effective sample size to 15-16 sALS and 14-16 PN depending on the cell type. (iii) The condition φ sALS-C9ALS comparison (Results §4) uses the 16 C9ALS donors paired with the same 10 PN donors that are used in the main pipeline. All three stages draw from the same underlying Pineda 2024 MCX cohort; the donor-count differences reflect only the differing sample-size requirements of SPD covariance estimation, pseudobulk DE, and PN-only φ_static estimation.

*Cell type selection: two complementary 10-cell-type sets.* Two different 10-cell-type sets are used in this paper, each matched to the statistical requirements of a specific pipeline. Nine cell types are shared between the two sets; the one-cell-type difference reflects distinct sample-size constraints at the Ledoit-Wolf covariance step (for Endothelial cells) and at the PN-only φ_static estimation step (for 5HT3aR interneurons).

The main pipeline (Part I cell type ranking, two-axis model, multi-transition integration, and all analyses that pool donors rather than splitting on condition) analysed the ten cell types Oligo, OPC, Astro, L2/3, L3/5, L4/6, L5/6, L6, PV, and 5HT3aR. Endothelial cells (Endo) were excluded from this pipeline because per-donor Endo cell counts (median 14-40 cells) are insufficient for stable Ledoit-Wolf covariance estimation at the gene set sizes used in the main pipeline (1,000-3,500 genes; Ledoit-Wolf requires approximately n_cells ≥ p/10 for stable shrinkage target estimation).

The 3φ residual framework (used for Sections 4.1 (condition φ R² table), 4.2 (rewiring/collapse analysis), 8.9 (NEMF CATylation bottleneck screen), and Fig. 3) analysed a different ten-cell-type set: the same nine core cell types (Oligo, OPC, Astro, L2/3, L3/5, L4/6, L5/6, L6, PV) plus Endothelial (Endo), with 5HT3aR replaced by Endo. The framework enabled Endo by reducing the base gene set to 200 genes per cell type, bringing the Ledoit-Wolf sample-size requirement within reach of Endo's cell counts (n_cells ≥ 20 with p = 200). Conversely, 5HT3aR was excluded from the residual framework because only one PN donor in the 24-donor panel had ≥ 150 5HT3aR cells, and stable φ_static estimation (a PN-only mean log-correlation matrix) requires at least three PN donors with adequate cell coverage. Both pipelines therefore share nine core cell types; the single-cell-type difference is a methodological consequence of each pipeline's sample-size requirements rather than a biological choice, and it does not affect any central finding.

Microglia, SOM interneurons, and other rare populations were excluded from both pipelines because per-donor cell counts were insufficient even at reduced gene set sizes. Preprocessing: log₁₀(CPM+1); OLS regression of donor, nUMI, and percent_mito covariates was applied before SPD covariance computation.

**Norman Perturb-seq.** scPerturb, 111,122 K562 cells (test: 34,624), 235 classes, 5,044 genes. log₁₀(TP10K+1). Leiden-clustering train/test split.

**CRISPR.** Shen et al. 2026, GSE274058, 41,749 mouse hippocampal cells, 18 perturbation conditions (17 single-gene KOs + 1 mSafe negative control). 16 targets evaluated (Ndufaf2 excluded: n = 12 KO cells below threshold; mSafe excluded: no corresponding gene).

**Glioma.** TCGA + GTEx (TOIL), log₂(TPM+1), 723 samples, 2,000 genes, 5 clinical windows.

**TimeVault.** Chao et al. 2026, PC9 cells, osimertinib, 12 bulk samples (4 conditions × 3 replicates), 17,543 genes.

#### S2. SPD geometry (original §6.2)

Per-donor PCA (k = 30). Ledoit-Wolf covariance. log-Euclidean mapping: log(C) = Q diag(log λ_i) Q^T.

#### S3. Pseudotime (original §6.3)

Two alternative pseudotime constructions were used to assess robustness. PT-B (covariance-based) is the primary construction used throughout the paper: diffusion pseudotime (DPT) was computed from diffusion maps on the inter-donor log-Euclidean distance matrix. Pairwise distances d_LE(C_i, C_j) = ‖log(C_i) − log(C_j)‖_F between donor SPD matrices defined a Gaussian kernel (bandwidth set to the median pairwise distance), from which the first non-trivial diffusion coordinate provided the pseudotime axis. PT-A (expression-based) is an independent construction used for sensitivity analysis: expression-level PCA was applied to per-donor mean-expression vectors (top 30 principal components), followed by diffusion map to extract the dominant progression axis. PT-A and PT-B order donors differently (window-assignment correlation ρ = −0.29) yet yield concordant gene-level φ rankings (Spearman ρ = 0.975 across cell types; Results §1), demonstrating that the Hodge gradient captures intrinsic co-expression network direction rather than pseudotime-imposed ordering. In both constructions, the 24 retained donors (14 sALS + 10 PN) were partitioned into 6 windows of 4 donors each, ordered by the pseudotime value. The sign convention orients the axis so that PN donors tend toward low pseudotime, but donor condition labels are not used during axis construction itself (no supervision by clinical metadata).

#### S4. Flow construction (original §6.4)

Delta matrix: Δ_t = mean(log C_{w_{t+1}}) − mean(log C_{w_t}). Edge-weight flow: f(i,j) = |Δ_ij| × sign(d_i − d_j), where d_i = ‖Δ_{i,·}‖₂.

Among antisymmetric edge flows that preserve entry-wise magnitudes |Δ_ij|, the edge-weight construction is the unique rule whose direction field is aligned with the discrete gradient of the node-level scalar d_i (node coherence). Alternative rules based on the entry-wise sign of Δ_ij determine direction locally per edge and do not satisfy node coherence; rules based on mean expression or PCA loadings are not invariant under orthogonal transformation of gene space. Under the Hodge decomposition f = grad(φ), the convention f(i,j) &gt; 0 when d_i &gt; d_j implies φ_j &gt; φ_i, so genes with the smallest d_i receive the highest φ (see Results §1, Level 1).

#### S5. Hodge decomposition (original §6.5)

K_N: φ = L⁺ div(f), where L⁺ = (1/N)(I − J/N). GF = ‖grad‖²/‖f‖².

k-NN: k-NN graph from delta row distances. φ via least-squares solver (LSQR; scipy.sparse.linalg.lsqr). Curl per triangle: Γ_{ijk} = f(i,j) + f(j,k) − f(i,k). Louvain community detection on curl-weighted network.

#### S6. Gradient fraction baseline (original §6.6)

GF_0 = 2/[3(1+CV²)], validated across 316 conditions (r = 0.994).

#### S7. Multi-transition integration (original §6.7)

Consistency-weighted averaging across 5 transitions. Stable-High: top quantile in ≥ 4/5 (100 bootstrap iterations).

#### S8. Global whitening for perturbation data (original §6.8)

W = V D^{−1/2} V^T from Ledoit-Wolf Σ_ctrl (eigenvalues clamped at 10⁻¹⁰). Bootstrap B = 100.

#### S9. WGCNA and ablation (original §6.9)

Per-cell type pseudobulk WGCNA (24 donors × N genes). Scale-free R² ≥ 0.80. Source module: lowest Top-50 Jaccard under removal. Module labels (black, darkgrey, silver, lightgrey, etc.) are WGCNA-assigned color identifiers and carry no biological meaning.

#### S10. Two-axis model (original §6.10)

Stable-High genes are those ranking in the top 10th percentile of φ in at least 4 of 5 consecutive-window transitions, identified via 100 bootstrap iterations over donors (resampling with replacement). The stable-High set per cell type defines the maximally preserved co-expression core.

TRS (translation-resource score) is defined per donor as the mean upper-triangular log-correlation among stable-High genes in that donor's covariance matrix; higher TRS indicates tighter coordination of the stable-High core within that donor. MSS (morphogenesis-structure score) is defined per donor as the first principal component of the expression matrix restricted to the source module (the WGCNA module most disrupted by ablation, Supplementary Methods S9). Across the 24 donors, TRS and MSS show independent variation (Spearman ρ = 0.049, p = 0.71), establishing them as empirically orthogonal axes per donor. The 2.84-fold glia-neuron TRS separation and the OPC-specific 2.44-fold TRS increase along pseudotime are reported in Results §3.

#### S11. TimeVault temporal foundation (original §6.11)

TimeVault data (Chao et al. 2026): PC9 lung adenocarcinoma cells, osimertinib 100 nM for 4 days. 12 bulk RNA-seq samples: 4 conditions (D/Dc/ND/NDc) × 3 replicates. Recorded (Dc, NDc) = vault-protected past RNA (7 days prior); Present (D, ND) = cytosolic current RNA. Gene classification: DEG analysis (t-test, BH-corrected) on Dc vs NDc (Recorded) and D vs ND (Present); *upstream* genes = significant in Recorded only (n = 5,458); *downstream* genes = significant in Present only (n = 547). The Temporal Asymmetry Score (TAS), defined in the present work (not in Chao et al.), is TAS = (|dmean_recorded| + |dvar_recorded|) − (|dmean_present| + |dvar_present|), where dmean and dvar denote differences in condition means and variances between drug-treated and non-treated groups within each RNA fraction. TAS is a per-gene univariate score; it does not involve gene-gene covariance, Hodge decomposition, or IDS φ. ROC-AUC computed for TAS discrimination of upstream versus downstream genes as defined above.

#### S12. Mendelian randomisation (original §6.12)

Two-sample MR using oligodendrocyte cis-eQTLs from Bryois et al. (2022; Zenodo 7276971, n = 196) as instruments and ALS GWAS from van Rheenen et al. (2021; GCST90027164, N_eff = 80,713) as outcome. Instruments selected at p < 5 x 10^-6 (relaxed from genome-wide significance to retain sufficient instruments given the limited overlap between cell type eQTLs and ALS GWAS loci), LD-clumped (r^2 < 0.001, 10 Mb window, EUR reference). Weak instruments (F < 10) excluded. Four methods: IVW, MR-Egger, weighted median, weighted mode. Sensitivity: Egger intercept, Cochran's Q, leave-one-out. Colocalization: coloc.abf (Giambartolomei et al. 2014), +/- 500 kb, top 100 genes. Software: R 4.4.3, TwoSampleMR 0.7.0, coloc.

#### S13. Statistical tests (original §6.13)

Wilcoxon signed-rank (CRISPR). McNemar (method comparison). Fisher's exact (enrichment). BH-FDR. Bootstrap (B = 100-1,000). LOCO; LODO.

#### S14. Software (original §6.14)

Python ≥ 3.10, PyTorch, anndata, scanpy, scikit-learn, scipy, PyDESeq2 0.5.0. Code: https://github.com/akacola2006/scrnaseq-hodge-pipeline (version pinned at commit 10d60e7, 2026-04-26; a release tag and Zenodo DOI will be deposited upon manuscript submission).

#### S15. Three-φ (3φ) residual framework (original §6.15)

The 3φ residual framework operates on the Endo-inclusive ten-cell-type set described in Methods (Data) (nine core cell types shared with the main pipeline plus Endo, with 5HT3aR excluded due to insufficient PN donor coverage). To separate disease-specific signal from healthy-state co-expression topology, the Gene Hodge pipeline was executed three times per cell type, differing only in the donor subset used. φ_static was computed from the 10 PN donors only, carries zero disease information, and captures the weighted strength of each gene in the healthy co-expression network. φ_disease was computed from the 14 sALS donors only and captures the disease-state co-expression structure. φ_condition was computed from the pooled 24-donor panel (14 sALS + 10 PN) and captures a mixture of topology and disease.

Per cell type, disease-specific residuals were computed by regressing φ_static from φ_disease in the raw φ coordinate space (not percentile space):

φ_disease_i = f(φ_static_i) + ε_i

where f is a cubic polynomial (numpy.polyfit, degree 3) fit across all genes tested. The choice of raw-φ over percentile space is deliberate: in percentile space, ceiling clustering of high-φ genes combined with local slope reversals (notably in L3/5) causes the cubic fit to over-absorb disease-specific signal into the topology term; in raw-φ space, linear and cubic fits yield essentially identical rankings with the cubic preserving neuron-specific slope changes. Linear, cubic, and bin-mean fits were compared on the L3/5 ribosome class in raw-φ space and all three gave concordant significance (p = 1.2×10⁻⁸, 8.2×10⁻⁷, and 1.2×10⁻⁶ respectively). Cubic (poly3) in raw-φ space is reported as the default.

Residuals were standardised per cell type (divided by σ of ε across all genes tested in that cell type) to yield residual z-scores. Genes with z &gt; +2 were classified as rewiring (formation of pathological co-expression not predicted by healthy topology); genes with z &lt; −2 as collapse (disruption exceeding topology prediction). The per-cell-type R² of the poly3 fit quantifies how much of φ_disease is explained by healthy-state topology (glia R² = 0.93-0.95, neurons R² = 0.58-0.85; Table 2, Results §4).

Matched null z-scores were computed to correct for gene-specific variance structure. For each gene, a null distribution was generated by permuting donor ALS/PN labels (1,000 permutations) while preserving cell type assignment, re-running φ_disease and the poly3 regression, and recording the residual z under each permutation. The matched null z-score is (z_observed − z_null_mean) / z_null_std. Matched null z-scores are used for the pathway-level upstream-ness tests in Results §5 (translation 9-10/10 CTs; chaperone 4/10; ATP synthase 3/10) and for the single-gene NEMF screen in Results §6.

For the sALS-vs-C9ALS manifold comparison (Results §4), φ_static was held fixed (computed from the 10 PN donors), and φ_disease was computed twice: once from the 14 sALS donors and once from the 16 C9ALS donors from the same NYGC cohort. The per-cell-type R² of the poly3 fit was compared between the two disease groups.

#### S16. Differential expression analysis (sALS-only DESeq2) and GSEA-style rank test (original §6.16)

For the DE values reported in Results §5 and Results §6, per-cell-type pseudobulk counts were computed by summing raw UMI counts across cells within each (donor, cell type) pair. PyDESeq2 0.5.0 was applied per cell type, following the DE pipeline of Pineda et al. (2024). A per-CT low-expression filter retained donors with ≥ 20 cells in that cell type and genes with ≥ 1 count in ≥ 3 donors. Size factors were estimated by the median-of-ratios method, and the Wald test was applied to the design formula ~ condition + sex, with sALS vs PN as the effect of interest and sex as a covariate. Shrunken log₂ fold changes (apeglm) are reported; p-values were adjusted by Benjamini-Hochberg FDR.

All DE values reported in the main text (Sections 4.2 and 8.9) refer to this sALS-only analysis, using the full 17 sALS + 16 PN pool before per-CT cell count filtering, with effective per-CT sample sizes of 15-16 sALS and 14-16 PN depending on the cell type. A parallel ALS-combined analysis (sALS + 16 C9ALS versus PN, same design) was performed as a sensitivity check; its per-gene neuronal magnitudes are reported inline in Results §5.

Gene class means (RPL+RPS for the cytoplasmic ribosome class; Translation_full for RPL+RPS+EIF+EEF) reported in Results §5 are unweighted means of per-gene log₂ fold change across genes within each class that passed the DESeq2 low-expression filter. Gene class membership was defined by HGNC gene symbol prefix. Per-class gene counts vary across cell types because of the DESeq2 filter (RPL+RPS: 24-88 genes per CT; Translation_full: 38-143 genes per CT).

*GSEA-style rank test.* Because translation-apparatus genes are expected to shift as a coordinated set with modest per-gene effects, we supplemented per-gene DESeq2 with a class-level rank test following Pineda et al. 2024's GSEA approach. For each cell type, per-gene log₂ fold changes were ranked transcriptome-wide, and a one-sided Mann-Whitney U test was applied to compare the rank distribution of the Translation class (RPL+RPS+EIF+EEF) to all other genes. The null hypothesis is that translation genes are not systematically shifted upward in sALS; the one-sided alternative is that they are. For the pooled-neuron analysis reported in Results §5, the four neuronal cell types (L2/3, L4/6, L5/6, PV) were concatenated before ranking and testing. Top-decile enrichment (the fraction of translation genes appearing in the top 10 % of the genome-wide log₂FC distribution versus the expected 10 % under the null) was computed as a descriptive measure of coordinated-shift magnitude.

For individual-gene DE values cited in Results §6 (NEMF, LTN1, GCN2 / EIF2AK4, PKR / EIF2AK2, PERK / EIF2AK3, LSM14A, EIF4EBP2, DROSHA, FXR1), the same sALS-only pipeline was used and padj &lt; 0.05 was taken as the significance threshold where reported.

#### S17. Pathway enrichment via Metascape and GSEApy/Enrichr (original §6.17)

Gene-set enrichment of rewiring genes (Results §5) was performed using Metascape (Zhou et al. 2019). Genes with residual z &gt; +1 (relaxed from the z &gt; +2 classification threshold to preserve statistical power for enrichment testing) were submitted per cell type to Metascape with default parameters: GO Biological Process, KEGG, Reactome, and CORUM term sets; enrichment p-value cut-off 0.01; minimum overlap 3; minimum enrichment 1.5. Enrichment p-values reported are Metascape-adjusted by Benjamini-Hochberg. Cross-cell-type preranked GSEA on φ-ranked gene lists (Results §5), top-φ gene GO enrichment (Results §5), WGCNA module enrichment (Sections 3.3 and 6.9), and the cross-cohort PT-C residual GSEA in Ruf2026 (Cross-Cohort Supplement) were performed via GSEApy (Fang et al. 2023), which queries the Enrichr gene-set collection (Chen et al. 2013; Kuleshov et al. 2016; Xie et al. 2021) for the GO_Biological_Process_2023, KEGG_2021_Human, and Reactome_2022 libraries. DisGeNET cross-reference (sporadic ALS, Parkinson disease, Alzheimer disease term associations) was performed via the Metascape DisGeNET module and interpreted with reference to the DisGeNET knowledge platform (Piñero et al. 2020).

#### S18. Robustness validations (original §6.18)

Gene-level φ rankings were tested against three orthogonal perturbations. Leave-one-donor-out (LOCO): the full pipeline was re-executed 24 times, each time excluding one donor; per-gene φ rankings across the 24 runs had cross-run Spearman ρ = 0.981 (mean across cell types). PT-A vs PT-B: replacing the primary covariance-based pseudotime (PT-B) with the expression-based alternative (PT-A) yielded Spearman ρ = 0.975 between gene-level φ rankings despite the two axes ordering donors differently (window-assignment ρ = −0.29). Gene selection method: replacing the PCA-based gene set (1,578 genes) with the detection-rate-based gene set (3,683 genes; Supplementary Note 2) yielded gene-level φ Spearman ρ = 0.992 and 9/9 cell types with the translation class significantly upstream (Wilcoxon one-sided p < 0.05). At the cell type level, leave-one-donor-out bootstrap (LODO) of the cell type ranking was stable in 24/24 iterations for the top-ranked oligodendrocyte position. These three robustness tests confirm that the translation upstream finding is not an artefact of donor composition, pseudotime construction, or gene selection. A Lipschitz-type theoretical bound on the LOCO perturbation of the rank-determining quantity d_i, in terms of the within-window log-Euclidean dispersion σ, is provided in Supplementary Note 1 (section "Formal stability of the φ ranking under leave-one-donor-out"); for the main pipeline parameters (n_w = 4 donors per window, W − 1 = 5 transitions), the theorem yields a rank-stability threshold of 4σ/15 ≈ 0.267 σ on pairwise d-gaps, which provides a computable sufficient condition for the empirical ρ = 0.981 reported above.

#### S19. Pipeline-specific methods for the speculative discussion (original §6.19)

The gene-level Hodge pipeline used in Part II (detection-rate-based gene selection, per-cell-type gene-level φ) is described in Methods (Differential expression and pathway enrichment). The module-level causal analysis pipeline used in Part III (23 functional modules, diffusion pseudotime PT_dpt on the module-score feature space, LiNGAM, and time-lag analysis) is described in Supplementary Discussion S1, §S1.1. These Part-specific descriptions are retained alongside their respective Parts for readability; all shared upstream methods (data preprocessing, SPD geometry, pseudotime, Hodge decomposition, 3φ residual framework, DE analysis, enrichment, and robustness validations) are described above in Sections 6.1-6.18.

### Supplementary Notes

#### Supplementary Note 1: Theoretical Background for the Stress-Covariance Framework

This note provides a short, snapshot-level account of why covariance structure on the SPD manifold carries temporal stress-trajectory information, and why the same methodology applies coherently across multiple biological scales. Full derivations are given in a companion theoretical manuscript in preparation (Kaneko, in preparation); here we sketch the minimum necessary to read the empirical pipeline as an implementation of a principle theory rather than a purely computational construction. None of the biological findings in Sections 3-4 or 8 logically require this note; readers who prefer to read the paper as methodology only may treat it as optional background.

##### A stress functional on the covariance manifold

Let C∈SPD(p) denote a state of the gene-gene covariance structure for p genes, and let X=logC denote its log-Euclidean tangent representation. Under the Log-Euclidean metric, distance between covariance states and their volume are both expressible in the same linearised coordinate X:

dLE(C,C0)=∥X-X0∥F,  logdetC=tr(X).

This allows a free-energy-like functional on SPD(p):

F(C∣C0)=∥X-X0∥F2-λ⋅tr(X).

The first term penalises structural distortion from a reference state C0 (an energy-like term); the second term rewards accessible covariance degrees of freedom (an entropy-like term under Gaussian approximation). We treat F as the stress functional associated with the covariance state. Between discrete reorganisation events, the system evolves by natural gradient flow

dCdt=-gradgF(C∣C0)

with metric g determined by a mirror potential on SPD(p). This is the SPD-manifold expression of the general principle that a system operating under finite resources evolves by coordinate-invariant stress minimisation.

##### Monotone dissipation and the direction of pseudotime

Under the natural gradient flow, is monotonically non-increasing (the norm below is the metric norm ‖·‖_g induced by the Riemannian metric g):F

dFdt=-∥gradgF∥2≤0.

Equality holds only at critical points of F. This Lyapunov-type dissipation is the theoretical basis for the directional interpretation of our pseudotime axis: the diffusion-map ordering of donors along the inter-donor log-Euclidean distance matrix (Methods 6.3) is used as a data-driven proxy for the dominant descent direction of F, not a time axis imposed from clinical metadata. The empirical confirmations in Results §2 - label-free PC1 recovery of the Recorded-vs-Present axis at AUC = 1.0, and a per-gene variance-asymmetry score discriminating upstream from downstream genes at AUC = 0.9475 - can be read as empirical verifications that variance dynamics do track the theoretically predicted stress-descent trajectory, with a strength that cannot be dismissed as a consequence of definitions.

##### Hodge decomposition and the disease signature

The change in covariance structure between consecutive pseudotime windows defines an antisymmetric edge flow ω on the gene-gene graph. Discrete Hodge decomposition orthogonally splits this flow into three components,

ω=dΦ+δΨ+h,

where dΦ is the irreversible gradient component (dissipative drift), δΨ is the curl component (local circulatory structure preserving flow without net dissipation), and h is the harmonic component (a topology-constrained skeleton that is neither gradient nor curl). In healthy configurations, the curl and harmonic components provide structural braking against runaway drift: circulation recycles resources locally and the harmonic skeleton anchors global topology. The theory predicts disease-associated brake loss as an abnormal *allocation* between irreversible drift and feedback/anchoring components, rather than as a simple global increase of the gradient fraction. In sALS, this appears as a reproducible translation-gradient/synaptic-curl dichotomy across cell types (Results §3) and as collapse-dominant residual deformation in neurons (Results §5).

##### Time-character reading: past, present, and persistent residue

The three Hodge components admit a reading in terms of the *time-character* of the stress dynamics they capture. The covariance state C(t) on the SPD manifold at any moment is the integral of all past stress events processed along the natural gradient flow of the free-energy functional F: it carries the history of past perturbations compressed into its present structure. Hodge decomposition of the covariance change flow along pseudotime separates this integrated history into three time-characters. The gradient component dΦ represents *currently-active irreversible drift* - stress that is being processed at this moment in the trajectory and has not yet been absorbed into stable structure. The curl component δΨ represents *transient circulatory dynamics* - oscillatory patterns that are neither accumulating into persistent form nor being dissipated away, a time-reversible middle ground. The harmonic component h represents the *persistent structural residue*: the part of the covariance state that cannot be reduced further by gradient descent or removed by local circulation, which therefore carries forward the accumulated imprint of past perturbations as stable topological constraints. In this reading, h encodes residue-from-past-dynamics in the structural sense (not in the neuroscientific sense of stored-for-later-retrieval): it is the component of the covariance state that has already absorbed the cumulative history of stress events and now holds that history as network topology.

The TimeVault empirical finding of Results §2 is consistent with this reading at the per-gene marginal level. Genes showing large variance changes in the Recorded (past) fraction but small variance changes in the Present fraction are genes whose stress response has already been processed and dissipated - the per-gene analogue of a contribution to the harmonic residue. Genes showing the opposite asymmetry are genes whose stress response is currently active - the per-gene analogue of a contribution to the gradient drift. The per-gene marginal asymmetry captured by the Temporal Asymmetry Score (AUC = 0.9475 against TimeVault-defined upstream/downstream labels) is therefore consistent with, though not mathematically equivalent to, the network-level time-character separation performed by Hodge decomposition in the main pipeline. The two operate on different objects (marginal distribution versus gene-gene covariance network) but realise the same underlying principle that variance dynamics carry recoverable temporal stress-trajectory information.

##### The harmonic component as a structural anchor

In the complete-graph KN implementation used for the main φ ranking, the literal harmonic component vanishes (sparse k-NN graphs in Results §3 admit non-zero harmonic components used in curl analysis). The &quot;harmonic anchor&quot; interpretation in this note refers to the topology-preserving core in the general stress-covariance theory, operationalized in KN by minimal correlation-change magnitude (low di) and high φ.

When the allocation between gradient and curl is stabilised, the harmonic component h represents structural degrees of freedom that can be reduced neither by gradient descent nor by local circulation. It persists as a topological constraint that anchors the network against collapse. Results §1 (Level 1) notes that on the complete graph, φ is monotone in the negative of the per-gene correlation change magnitude di (Spearman ρ≈-1.000): the genes we call &quot;upstream&quot; in the pipeline are the genes whose network relationships are least perturbed along the progression axis. In the theoretical language of this note, these are the genes that most fully participate in the harmonic anchor - the structural core that neither flows nor circulates but holds the network in place. The observation that translation genes occupy this anchor position in every cell type tested (Results §5) is therefore not an accidental empirical pattern but the expected biological realisation of a harmonic-subspace core.

##### Nested architecture: the same framework at multiple scales

A central feature of the theory is that the dynamics described above are scale-free: the same free-energy functional ℱ, the same natural gradient flow, and the same Hodge decomposition apply at every nesting level of the covariance architecture. In the language of this paper, each observable covariance state C at one scale can be unpacked into components (gene modules, pathways, sub-networks) that are themselves SPD states at a finer scale, each with its own ℱ, its own gradient flow, and its own Hodge split. Concretely, the main pipeline operates at the cell-type scale (each cell type carries a gene-gene SPD trajectory; Results §5); the residual framework descends to the pathway scale (modules such as cytoplasmic translation or oxidative phosphorylation carry their own internal structure; Results §5); and the single-gene analysis descends further to the individual-gene scale (for example the NEMF CATylation bottleneck screen; Results §6). The same 3φ residual framework applies at each of these levels because each level is a covariance state on its own SPD manifold. None of these scales is fundamental in the reductionist sense - the pipeline's ability to move seamlessly between them reflects the nested structure of the underlying theory rather than a computational coincidence.

##### Formal stability of the φ ranking under leave-one-donor-out

The leave-one-donor-out robustness observations reported in Methods 6.18 admit a direct Lipschitz-type stability bound on the log-Euclidean tangent space. We state the bound here in full, because the argument is elementary and because its constants are computable from observable quantities, so the theorem can be checked against data without appealing to the companion theory.

**Setup and notation.** Let Ck∈SPD(p) denote donor k's gene-gene covariance matrix and Lk=logCk its log-Euclidean representation (Methods 6.2). The 24 retained donors are partitioned into W=6 pseudotime windows of nw=4 donors each (Methods 6.3). For each window w∈{1,…,W}, write the within-window mean as Lw=(1/nw)k∈wLk, and for each transition w∈{1,…,W-1} write the delta matrix as Δw=Lw+1-Lw. For each gene i, define the transition-averaged row norm

di:=1W-1w=1W-1∥Δw,i,⋅∥2,

where is the -th row of . This is a conservative summary of the per-transition row norms that drive the edge-flow construction in Methods 6.4. The multi-transition integration of Methods 6.7 identifies genes whose top-quantile position is preserved in at least 4 of 5 consecutive transitions (stable-High), which is a strictly more demanding stability criterion than stability of the averaged d_i; the Lipschitz bound below therefore provides a sufficient condition for stability of the transition-averaged d_i / phi ranking. Stability of stable-High membership, which depends on per-transition top-quantile thresholds, additionally requires per-transition margin control beyond what the averaged-d_i bound alone guarantees. Since φ on the complete graph K_N is a monotone decreasing transformation of d_i up to Spearman ρ ≈ −1.000 (Results §1, Level 1), rank stability of d_i is rank stability of φ.Δw,i,⋅iΔwdidi(w)=∥Δw,i,⋅∥2

Let the within-window dispersion at window w be

σw:=maxk∈w∥Lk-Lw∥F,

and the global within-window dispersion be σ:=maxwσw. Both σw and σ are computable from the data (they are Frobenius distances in the log-Euclidean tangent space) and are therefore empirically observable constants.

We define fixed-window leave-one-donor-out as the operation that removes donor k from its assigned window w(k) while leaving all other donor-to-window assignments and the window partition unchanged. We write Lw(-k), Δw(-k), and di(-k) for the resulting window means, delta matrices, and averaged row norms. The empirical LODO evaluation reported in Methods 6.18 goes further and recomputes the diffusion-map pseudotime after donor removal; we return to this distinction after the main theorem.

**Lemma 1 (window-mean perturbation).** Under fixed-window LODO, window means are unchanged for w≠w(k), and for w=w(k),

∥Lw(k)(-k)-Lw(k)∥F=∥Lk-Lw(k)∥Fnw-1.

*Proof.* Windows not containing donor k have Lw(-k)=Lw by definition. For w=w(k),

Lw(k)(-k)=1nw-1j∈w(k), j≠kLj=nwLw(k)-Lknw-1.

Subtracting Lw(k) from both sides,

Lw(k)(-k)-Lw(k)&=nwLw(k)-Lknw-1-Lw(k)&=nwLw(k)-Lk-(nw-1)Lw(k)nw-1&=Lw(k)-Lknw-1.

Taking Frobenius norm of both sides gives the stated identity. For nw=4 this is σw(k)/3, bounded above by σ/3. ▫

**Lemma 2 (delta-matrix perturbation).** Under fixed-window LODO, the delta matrices are unchanged for all transitions w∉{w(k)-1,w(k)}∩{1,…,W-1}. The number of affected transitions is at most 2: exactly 2 when w(k) is an interior window (2≤w(k)≤W-1), and exactly 1 when w(k) is a boundary window (w(k)=1 or w(k)=W). For each affected transition w,

∥Δw(-k)-Δw∥F≤σnw-1.

*Proof.* By definition, Δw=Lw+1-Lw. The only window mean changed by the LODO operation is Lw(k) (Lemma 1). Hence Δw is perturbed if and only if w or w+1 equals w(k), i.e., w∈{w(k)-1,w(k)}. Transitions outside this set satisfy Δw(-k)=Δw exactly. Boundary cases: if w(k)=1, the candidate w=w(k)-1=0 is not a valid transition index, so only w=1 is affected; if w(k)=W, the candidate w=w(k)=W is not a valid transition index, so only w=W-1 is affected. For each affected transition w:

If w=w(k)-1: Δw(-k)-Δw=(Lw(k)(-k)-Lw(k)-1)-(Lw(k)-Lw(k)-1)=Lw(k)(-k)-Lw(k), using that Lw(k)-1 is unchanged and w+1=w(k).

If w=w(k): Δw(-k)-Δw=(Lw(k)+1-Lw(k)(-k))-(Lw(k)+1-Lw(k))=-(Lw(k)(-k)-Lw(k)), using that Lw(k)+1 is unchanged.

In both cases, taking Frobenius norm and applying Lemma 1,

∥Δw(-k)-Δw∥F=∥Lw(k)(-k)-Lw(k)∥F=∥Lk-Lw(k)∥Fnw-1≤σw(k)nw-1≤σnw-1. ▫

**Theorem (fixed-window LODO Lipschitz bound).** Under fixed-window LODO, for every gene i and every donor k,

| di(-k)-di | ≤2σ(nw-1)(W-1). |
| --- | --- |

*Proof.* By definition,

di(-k)-di=1W-1w=1W-1∥Δw,i,⋅(-k)∥2-∥Δw,i,⋅∥2.

Taking absolute value and using the triangle inequality for the outer sum,

| di(-k)-di | ≤1W-1w=1W-1 | ∥Δw,i,⋅(-k)∥2-∥Δw,i,⋅∥2 | . |
| --- | --- | --- | --- |

For each transition w, the reverse triangle inequality for the Euclidean norm gives

| ∥Δw,i,⋅(-k)∥2-∥Δw,i,⋅∥2 | ≤∥Δw,i,⋅(-k)-Δw,i,⋅∥2. |
| --- | --- |

The per-row Euclidean norm of any matrix M is bounded above by the Frobenius norm of M:

∥Mi,⋅∥2=jMi,j2≤i',jMi',j2=∥M∥F,

so

∥Δw,i,⋅(-k)-Δw,i,⋅∥2≤∥Δw(-k)-Δw∥F.

By Lemma 2, the right-hand side vanishes for all transitions outside {w(k)-1,w(k)}∩{1,…,W-1}, and is at most σ/(nw-1) for each transition inside this set. Since the set contains at most 2 elements,

w=1W-1∥Δw(-k)-Δw∥F≤2σnw-1.

Substituting into the outer sum,

| di(-k)-di | ≤1W-1⋅2σnw-1=2σ(nw-1)(W-1).  ▫ |
| --- | --- |

**Corollary (pairwise rank stability).** For any two genes i,j, if |di-dj|>4σ/[(nw-1)(W-1)], then for every LODO removal k, sign(di(-k)-dj(-k))=sign(di-dj).

*Proof.* By the theorem applied to both genes,

| (di(-k)-dj(-k))-(di-dj) | &= | (di(-k)-di)-(dj(-k)-dj) | &≤ | di(-k)-di | + | dj(-k)-dj | &≤4σ(nw-1)(W-1). |
| --- | --- | --- | --- | --- | --- | --- | --- |

If |di-dj| strictly exceeds this bound, then (di(-k)-dj(-k)) cannot change sign relative to (di-dj). ▫

**Numerical instantiation for the main pipeline.** With nw=4 and W-1=5 (Methods 6.1, 6.3), the theorem gives |di(-k)-di|≤2σ/15, and the corollary gives the rank-stability threshold

gap*(i,j):=4σ15≈0.267⋅σ.

Gene pairs whose d-gap exceeds roughly a quarter of σ are provably rank-stable under fixed-window LODO. This is a sufficient condition for the empirical rank stability observed in Methods 6.18 (LOCO mean Spearman ρ=0.981).

**Scope and relation to the empirical LODO evaluation.** The bound above applies strictly to the fixed-window LODO operation defined in the setup, in which the donor-to-window assignment and the window partition are held constant. The empirical LODO evaluation reported in Methods 6.18 additionally recomputes the diffusion-map pseudotime on the remaining 23 donors and reconstructs the window partition, which introduces a secondary perturbation through shifted window boundaries. The fixed-window bound therefore captures the dominant perturbation source but does not, on its own, bound the full empirical LODO operation. Two facts make the additional pseudotime-recomputation contribution small in practice. First, pseudotime is itself highly stable under alternative constructions: replacing the primary covariance-based PT-B with an expression-based PT-A (Methods 6.3) yields gene-level Spearman ρ=0.975 for the φ ranking across cell types, despite the two pseudotime axes ordering donors differently at the window-assignment level (ρ=-0.29). Window membership is therefore robust to perturbations far larger than single-donor removal. Second, removal of one donor from a panel of 24 can reshuffle at most one window boundary by at most one donor position, contributing a residual window-mean perturbation of order at most σ/nw, which is of the same order as (and typically smaller than) the dominant term already captured by the theorem. A full proof including pseudotime recomputation would make the second fact quantitative; we leave that technical extension to the companion theoretical manuscript (Kaneko, in preparation). For the purposes of reading Methods 6.18 as empirical confirmation, the theorem above is the operative finite-sample guarantee: it bounds the dominant perturbation source in terms of an observable quantity σ, and its numerical instantiation (≈0.267 σ gap threshold) can be checked directly against the data.

**Relation to the Birkhoff-Hopf contraction framework.** The companion theoretical manuscript (Kaneko, in preparation) contains a separate proposition establishing projective stability of hierarchical composition under positive primitive updates, via the Birkhoff-Hopf contraction theorem. That proposition is asymptotic in nature: it guarantees convergence to a unique projective attractor as the number of hierarchical levels grows. It supports the background assumption that σ itself remains bounded as the analysis is extended to larger donor panels or finer window resolutions. The Lipschitz bound stated above is complementary: it provides a direct, finite-sample, computable stability guarantee at fixed W, using only σ as the observable constant. For the present paper, the Lipschitz bound is the operative tool because Methods 6.18 reports finite-sample LODO robustness rather than an asymptotic attractor statement.

##### Scope of this note

This note is deliberately a snapshot. It describes only the ingredients needed to read the pipeline as an implementation of a principle theory of stress on covariance structures: the Log-Euclidean free-energy functional, its natural gradient flow and Lyapunov dissipation, the Hodge decomposition with its brake-loss disease signature, the harmonic anchor interpretation of upstream genes, the scale-free nested architecture that justifies applying the same framework at cell-type, pathway, and single-gene resolutions, and the Lipschitz stability bound that quantifies the empirical LODO robustness reported in Methods 6.18. Additional results in the companion theoretical manuscript (Kaneko, in preparation) - including the Birkhoff-Hopf projective stability proposition referenced above, and corollaries involving the golden ratio as a minimax-optimal binary allocation ratio under self-similar relay cost - are not needed for any biological claim in this paper. The golden ratio in particular appears in the theory as a downstream corollary of the minimal binary case; it plays no role in the ALS analysis and should not be taken as a central claim of this work.

#### Supplementary Note 2: PCA Selection Sensitivity Analysis

##### Motivation

PCA-based dimensionality reduction preferentially preserves genes that participate in large coordinated covariance blocks. If Hodge-φ rankings are driven by this selection step rather than by genuine structural features of the disease transcriptome, the biological conclusions reported in the main text would be artifactual. We therefore constructed an alternative gene set that involves no PCA at any stage and re-ran the full φ pipeline.

##### Alternative gene selection (detection-rate method)

For each donor, a gene was retained if it was expressed (count > 0) in at least 10 % of cells. A gene entered the final set only if it met this criterion in 80 % or more of donors. This procedure yielded 3 683 genes (compared with 1 578 in the PCA-based set). Because the detection-rate set is strictly defined by per-donor prevalence, it is free of any PCA-induced covariance bias. A reduced 1 578-gene detection-rate set was also analysed to control for gene-set size.

##### Locked screening metrics and decision rules

Two concordance measures were specified before running the sensitivity analysis:

**Table 27. PCA selection sensitivity concordance metrics.**

| Metric | Definition | Threshold |
| --- | --- | --- |
| Gene-level φ Spearman ρ | Rank correlation of per-gene φ between PCA and detection-rate sets | > 0.8 → LOW_SENSITIVITY; 0.5—0.8 → MODERATE; < 0.5 → HIGH |
| Pathway median-φ rank correlation | Rank correlation of pathway-level median φ between the two sets | Same thresholds |

##### Results

*Gene-level concordance.* The Spearman rank correlation of gene-level φ values between PCA-selected and detection-rate-selected gene sets was ρ = 0.992 (LOW_SENSITIVITY). Individual gene upstream positions are therefore robust to the choice of selection method.

*Pathway-level concordance.* Pathway median-φ rank correlation was ρ = 0.630 (MODERATE_SENSITIVITY). Most pathways retained similar relative positions, but a notable exception was sphingolipid metabolism, whose rank shifted from 21 (PCA set) to 2 (detection-rate set) in oligodendrocytes. This indicates that PCA selection had excluded sphingolipid-associated genes from the analysed gene space, thereby masking an additional upstream programme.

*Cross-cell type Wilcoxon validation.* To test whether the translational/ribosomal upstream finding depends on gene selection method, we performed one-sided Wilcoxon rank-sum tests comparing φ values of translation/ribosome-annotated genes against all other genes, separately in each cell type. Ribosomal genes were significantly upstream in 9/9 cell types (Wilcoxon one-sided p < 0.05; mean rank-biserial correlation r = −0.50), confirming that translational structural precedence is independent of PCA selection.

Other pathways could not be statistically confirmed across cell types owing to insufficient gene coverage in the detection-rate set.

##### Interpretation

The sensitivity analysis reveals that PCA bias operates not by distorting φ estimates for the genes it retains (ρ = 0.992) but by omitting entire gene neighbourhoods from the analysed space. The primary biological conclusion, translational/ribosomal structural precedence, is fully robust. However, the complete upstream landscape is likely broader than PCA-based analysis alone suggests, with sphingolipid metabolism in oligodendrocytes constituting one such additional upstream programme.

#### Supplementary Note 3: Axiomatic Justification of the Edge-Weight Flow Construction

##### Purpose

The main pipeline constructs an edge flow from the pseudotime delta matrix Δ via

f(i,j)=|Δij|⋅sign(di-dj),  di:=∥Δi,⋅∥2.

This note establishes that, given a minimal set of structural axioms, the edge-weight construction satisfies a conditional uniqueness property (Proposition 3.1) and admits a variational characterization that is unique without separately postulating the node-scalar map in A3, once the engagement scalar d_i = ‖Δ_i,·‖₂ is fixed in the alignment objective (Theorem 3.3).

3.1 Setting

Let p denote the number of genes and let Δ∈Rp×p be symmetric, encoding the window-to-window change of the log-Euclidean covariance representation (Methods 6.4). Define the row engagement scalar di:=∥Δi,⋅∥2. We seek an antisymmetric edge flow f on Kp, i.e., f(i,j)=-f(j,i).

3.2 Axiom System

**Axiom A1 (Antisymmetry).** f(i,j)=-f(j,i) for all i≠j.

**Axiom A2 (Magnitude fidelity).** |f(i,j)|=|Δij| for all i≠j.

**Axiom A3 (Node coherence).** The sign of f(i,j) depends only on node-level scalars, not on edge-local information beyond |Δij|. Formally, there exist σ:R2→{-1,0,+1} and a node scalar map η:Rp×p→Rp such that sign(f(i,j))=σ(η(Δ)i,η(Δ)j).

**Axiom A4 (Permutation invariance).** The flow is equivariant under relabeling of genes. This rules out constructions referencing information external to Δ (e.g., expression levels, PCA loadings).

**Axiom A5 (Energy fidelity).** i<j|f(i,j)|2=i<j|Δij|2.

3.3 Conditional Uniqueness

**Proposition 3.1 (Conditional uniqueness). Suppose , and assume strict separation d_i ≠ d_j on every edge with |Δ_ij| > 0 (ties may be assigned by any fixed deterministic antisymmetric rule). Under axioms A1—A5 together with order-preservation of the sign rule (sigma(a,b) = +1 whenever a > b), the unique flow (up to global sign) is**η(Δ)i=di:=∥Δi,⋅∥2

f⋆(i,j)=|Δij|⋅sign(di-dj).

**Proof.** By A1—A2, f(i,j)=s(i,j)|Δij| with s(j,i)=-s(i,j). By A3 with η=d, s(i,j)=σ(di,dj). Antisymmetry forces σ(a,b)=-σ(b,a). Setting σ(a,b)=sign(a-b) yields f⋆, which satisfies A1—A4 by construction and A5 since ∑|f⋆|2=∑Δij2. Any other monotone σ yields the same sign field on non-tied edges and hence the same Hodge decomposition. ▫

**Note.** Axioms A1—A5 do not uniquely determine the node scalar η: alternatives such as the L1 row norm also satisfy A1—A5. The choice η=di (row L2 norm) is canonical because (i) it uses all row entries with equal weight, (ii) it equals the Frobenius norm of the i-th row of the log-Euclidean tangent vector, and (iii) it aligns with the natural gradient descent interpretation (§3.5). The variational characterization (Theorem 3.3) provides uniqueness without this auxiliary choice.

3.4 Alternative Constructions and Axiom Violations

Note: in Table 28, formulas are interpreted as ordered-pair definitions on i < j, antisymmetrically extended to all (i,j). Under this convention, the entry sign(Delta_ij) is row-dependent only if extended antisymmetrically; treated as a symmetric formula it would violate A1.

**Table 28. Alternative flow constructions and axiom violations.**

| Construction | A1 | A2 | A3 | A4 | A5 |
| --- | --- | --- | --- | --- | --- |
| f(i,j) = sign(Δᵢⱼ) | ✓ | Fails | Fails (edge-local) | ✓ | Fails |
| f(i,j) = sign(μᵢ − μⱼ) (expression) | ✓ | Fails | ✓ | Fails (μ external) | Fails |
| f(i,j) = sign(ℓᵢ² − ℓⱼ²) (PCA loading) | ✓ | Fails | ✓ | Fails (ℓ external) | Fails |
| f(i,j) = Δᵢⱼ (raw entry) | Fails (Δ symmetric) | — | — | — | — |
| f(i,j) = sign(dᵢ − dⱼ) (sign flow) | ✓ | Fails | ✓ | ✓ | Fails |
| f(i,j) = │Δᵢⱼ│·sign(dᵢ − dⱼ) (edge-weight) | ✓ | ✓ | ✓ | ✓ | ✓ |

3.5 Variational Characterization

Let FΔ:={f∣f(i,j)=-f(j,i), |f(i,j)|=|Δij|}. Define the alignment

A(f):=i<jf(i,j)⋅(di-dj).

**Theorem 3.3 (Variational characterization).** *Assume* di≠dj *for all* i≠j*. The unique solution to* maxf∈FΔA(f) *is* f⋆(i,j)=|Δij|⋅sign(di-dj)*.*

**Proof.** The objective is additively separable: A(f)=i<j|Δij|⋅s(i,j)⋅(di-dj). Since (di-dj)≠0 and s(i,j)∈{-1,+1}, the maximum on each edge is achieved uniquely by s(i,j)=sign(di-dj). ▫

Under the Hodge decomposition with , the convention when implies : genes with the smallest receive the highest (Results §1, Level 1). Theorem 3.3 states that maximizes the inner product with the gradient field of among all flows in --- i.e., the flow component aligned with the node-engagement descent direction.f=grad(ϕ)+curl+harmonic(B0⊤ϕ)(i,j)=ϕj-ϕif(i,j)&gt;0di&gt;djϕj&gt;ϕidiϕf⋆-dFΔ

The row norm di=∥Δi,⋅∥2 measures the aggregate log-Euclidean displacement energy through node i per pseudotime step under the natural gradient flow of the stress functional (Supplementary Note 1). Theorem 3.3 then states that f⋆ is the unique energy-preserving discrete flow maximally aligned with this engagement ranking. A complete derivation connecting the continuous natural gradient flow to the discrete construction is deferred to the companion theoretical manuscript (Kaneko, in preparation).

#### Supplementary Note 4: Analytical Derivation of the Gradient Fraction Null Baseline GF₀

##### Purpose

The main text (Results §1) states the closed-form null baseline GF0=2/[3(1+CV2)] (r = 0.994 across 316 simulations). This note provides the analytical proof under an explicit random flow model.

4.1 Sign Flow Exact Identity

**Lemma 4.1 (for N ≥ 2). Let and . On ,**d1>d2>⋯>dNfsign(i,j)=sign(di-dj)KN

GFsign=2(N+1)3N.

**Proof.** WLOG d1>⋯>dN, so fsign(i,j)=+1 for all i<j and ∥fsign∥2=N2. Using the incidence-matrix convention (edges oriented from lower to higher index), the divergence at node v is divv=(v-1)-(N-v)=2v-N-1; Hodge projection then gives ϕv=(2v-N-1)/N. The gradient energy is

∥fgradsign∥2=4N2k=1N-1k2(N-k)=(N-1)(N+1)3,

using k=1N-1k2(N-k)=N2(N-1)(N+1)/12. Therefore GFsign=2(N+1)3N. ▫

*Verification*: N=901: predicted 0.667407, pipeline output 0.667407, residual <5×10-16.

4.2 Random Flow Model

Let {wij}i<j be non-negative i.i.d. random variables with mean μ, second moment μ2, and CV2=μ2/μ2-1. The random edge-weight flow is f(i,j)=wij⋅sign(di-dj).

Independence hypothesis (H_indep). Edge weights are mutually independent and identically distributed. This is satisfied exactly by random-matrix baselines with i.i.d. entries. In the actual row-norm-induced flow used in the pipeline, the edge weights w_ij = |Δ_ij| and the sign field sign(d_i − d_j) are both functions of Δ and not strictly independent; the closed form below should therefore be interpreted as an analytically tractable null baseline whose applicability to random-matrix preprocessing is verified empirically (r = 0.994 across 316 simulations).{wij}

4.3 GF₀ Theorem

**Theorem 4.2 (large-N ratio-of-expectations baseline). Define GF_0(N, CV) := E[‖P_grad f‖²]/E[‖f‖²] as the deterministic ratio-of-expectations baseline. Under H_indep,**

GF_0=2(N+1)3N(1+CV2)+O(1/N) →N→∞ 23(1+CV2).

**Proof.**

*Step 1.* E[∥f∥2]=N2μ2.

*Step 2.* Decompose f=μ⋅s⋆+δ where δij=(wij-μ)s⋆(i,j), E[δ]=0, independent entries with variance μ2CV2.

*Step 3.* By linearity of Pgrad and orthogonality:

E[∥Pgradf∥2]=μ2∥Pgrads⋆∥2+E[∥Pgradδ∥2].

By Lemma 4.1, μ2∥Pgrads⋆∥2=μ2(N-1)(N+1)/3. Since Pgrad has rank N-1 and δ entries are i.i.d.: E[∥Pgradδ∥2]=(N-1)μ2CV2.

*Step 4.* Dividing Steps 3 by Step 1 and substituting μ2=μ2(1+CV2):

GF_0≈2(N+1)3N(1+CV2)+2CV2N(1+CV2)=23(1+CV2)+O(1/N).  ▫

Concentration. Under H_indep with finite fourth moments of edge weights, |f|^2 concentrates around its mean, so GF = |P_grad f|^2 / |f|^2 concentrates around the ratio of expectations as N grows. Empirically, this concentration is verified at N = 922 by r = 0.994 across 316 random matrix simulations (Appendix Q of sALS Validation Report).N=922r=0.994∥f∥2N

4.4 Regime of Validity and Falsifiability

Theorem 4.2 holds under H_indep, satisfied by random matrix baselines and biological data with spatially uncorrelated edge-weight variation. Block-structured (module-type) signals under-predict GF by |δGF|≈0.02, consistent with the sALS residual |δGF|=0.027.

Sharp prediction: |GFnull-2/[3(1+CV2)]|<0.01 for any i.i.d. preprocessing. Verified: sALS (N=922): 5×10-4; glioma (N=2000): 6×10-4; 316 random simulations: r=0.994.

#### Supplementary Note 5: Stage-by-Stage Computational Details of the IDS Pipeline

This note provides a unified, implementation-level description of the five computational stages of the Intrinsic Direction System (IDS) pipeline. It is intended to bridge the conceptual overview in Results §1 and the condensed formulas in Methods (Five-stage Hodge pipeline). Theoretical justifications for key design choices are cross-referenced to Supplementary Notes 1, 3, and 4 rather than reproduced here.

##### Stage 1: Covariance Estimation on the SPD Manifold

**Input.** For each donor k and cell type, a cell × gene expression matrix after preprocessing: log₁₀(CPM+1), followed by OLS regression to remove donor, nUMI, and percent_mito covariates (Methods (Data)).

**Step 1.1 — PCA gene selection.** PCA is applied to the pooled (all-donor) expression matrix. The p genes contributing most to the top 30 principal components are retained (1,000–3,500 genes depending on cell type and pipeline). Sensitivity to this selection is assessed in Supplementary Note 2.

**Step 1.2 — Per-donor covariance estimation.** For each donor k, compute the gene-gene covariance matrix C(k) using Ledoit-Wolf shrinkage:

C(k)=(1-α)Σ(k)+α μ I,

where Σ(k) is the sample covariance, and α, μ are the analytically optimal shrinkage coefficient and target intensity (Ledoit and Wolf 2004). Shrinkage is applied because the number of cells per donor per cell type may be of the same order as p, making the sample covariance ill-conditioned.

**Step 1.3 — Log-Euclidean mapping.** Map each C(k)∈SPD(p) to its log-Euclidean tangent representation:

L(k)=logC(k)=Q(k) diag(logλi(k)) (Q(k))T,

where Q(k) and {λi(k)} are the eigenvectors and eigenvalues of C(k). This mapping linearises the curved SPD manifold while preserving Riemannian distances: dLE(Ci,Cj)=∥L(i)-L(j)∥F (Arsigny et al. 2006). The theoretical motivation — treating L(k) as a coordinate on the stress manifold governed by Lyapunov dissipation — is developed in Supplementary Note 1.

**Output.** Per-donor symmetric matrices {L(k)}k=1N.

##### Stage 2: Pseudotime Construction

**Input.** {L(k)}k=1N; donor condition labels are not used.

**Step 2.1 — Pairwise distance matrix.** Compute the N×N matrix of log-Euclidean Frobenius distances:

Dij=∥L(i)-L(j)∥F.

**Step 2.2 — Gaussian kernel.** Convert distances to affinities:

Kij=exp-Dij2/ε,

where the bandwidth ε is set to the median pairwise squared distance (standard choice for diffusion maps; Coifman and Lafon 2006).

**Step 2.3 — Diffusion map and DPT.** Row-normalise K to obtain the Markov (diffusion) operator P. Compute its eigendecomposition via scipy.sparse.linalg.eigsh. The first non-trivial eigenvector (skipping the constant eigenvector ψ0=1) defines the primary diffusion coordinate. Diffusion pseudotime (DPT; Haghverdi et al. 2016) is computed as geodesic distance in diffusion space from a fixed root donor (the PN donor with the lowest first diffusion coordinate value).

**Step 2.4 — Window partition.** Sort the N=24 donors by DPT value. Orient the axis so that PN donors tend toward lower DPT (without using condition labels to enforce separation). Partition into W=6 windows of nw=4 donors each. Robustness to pseudotime construction is evaluated using two independent constructions (PT-A: expression-based; PT-B: covariance-based) that order donors differently (window-assignment ρ=-0.29) yet yield concordant gene-level φ rankings (Spearman ρ=0.975; Results §1 and Methods (Five-stage Hodge pipeline, Stage 2)).

**Output.** Window membership w(k)∈{1,…,W} for each donor k.

##### Stage 3: Delta Matrix and Flow Construction

**Input.** Window membership {w(k)}; log-Euclidean matrices {L(k)}.

**Step 3.1 — Window-averaged tangent matrices.**

Lw=1nwk: w(k)=wL(k).

**Step 3.2 — Delta matrices.** For each consecutive window pair t∈{1,…,W-1}:

Δt=Lt+1-Lt.

The (i,j) entry of Δt encodes the change in log-covariance between genes i and j across transition t.

**Step 3.3 — Node engagement scalar.** For each gene i and transition t:

di(t)=∥Δt, i,⋅∥2,

the l2 norm of the i-th row of Δt, measuring how much gene i's co-expression pattern shifted across transition t.

**Step 3.4 — Edge-weight flow.** For each gene pair (i,j) and transition t:

f(t)(i,j)=|Δt,ij|×sign(di(t)-dj(t)).

This is antisymmetric: f(t)(i,j)=-f(t)(j,i). The magnitude |Δt,ij| records how much the pairwise co-expression changed; the sign encodes direction from the gene with greater overall correlation shift toward the gene with lesser shift. Among all antisymmetric flows preserving entry-wise magnitudes |Δt,ij|, this is the unique construction whose direction field is aligned with the discrete gradient of di (Supplementary Note 3, Theorem 3.3).

**Output.** Antisymmetric edge flow f(t) on KN (or k-NN graph; see Stage 5) for each transition t∈{1,…,5}.

##### Stage 4: Hodge Decomposition and φ Computation

**Input.** Edge flow f(t) on a graph G with p nodes (genes).

The discrete Hodge decomposition (Eckmann 1944; Lim 2020) orthogonally splits any antisymmetric edge flow f into:

f=∇φ⏟gradient (irreversible)+δΨ⏟curl (circular)+h⏟harmonic,

where ∇ is the |E|×p oriented incidence matrix. Genes with high φ are structurally upstream; genes with low φ are downstream. For the theoretical interpretation of the three components in terms of disease dynamics (brake-loss signature, time-character reading), see Supplementary Note 1.

**K_N mode (primary — stable ranking and multi-transition integration).**

On the complete graph, the harmonic component vanishes identically. The graph Laplacian pseudoinverse has the closed form:

L+=1NI-JN,

where J is the all-ones matrix. The gradient potential is:

φ=L+ div(f),  div(f)i=j≠if(i,j).

On KN under strict ordering of di, φi is exactly linear in the rank of di, implying Spearman ρ=-1 algebraically (Results §1, Level 1); empirical values ρ≈-1.000 accommodate tie-handling. Genes with the smallest di (least correlation-shifted) receive the highest φ (most upstream). The gradient fraction:

GF=∥∇φ∥2/∥f∥2

quantifies the proportion of flow energy in the irreversible cascade component. Its closed-form null baseline GF0=2/[3(1+CV2)] is derived in Supplementary Note 4. Rank stability of φ under leave-one-donor-out perturbation is bounded analytically in Supplementary Note 1 (Lipschitz stability section).

**k-NN mode (signal amplification and curl analysis).**

Construct a k-nearest-neighbour graph on p genes using Euclidean distances between Δ row vectors (k=50 or k=100; Methods (Five-stage Hodge pipeline, Stage 4)). Solve the least-squares problem:

φ=argminφ∥f-∇φ∥2

via scipy.sparse.linalg.lsqr. The curl component per triangle (i,j,k):

Γijk=f(i,j)+f(j,k)-f(k,i).

Community structure in the curl-weighted network is detected by Louvain clustering (python-louvain). On sparse graphs, δGF≈0.14 (approximately five-fold amplification of the deviation from null, relative to the KN deviation δGFKN≈0.025; §2.2), making the k-NN mode informative for per-transition signal amplification. The two graph modes are not alternatives but complementary: KN provides stable, topology-anchored rankings; k-NN provides per-transition signal discrimination and curl community structure.

**Output.** Per-gene φ(t)∈Rp and GF(t)∈[0,1] for each transition t.

##### Stage 5: Multi-Transition Integration

**Input.** Per-transition φ(t) for t=1,…,W-1=5.

**Step 5.1 — Per-transition quantile ranking. For each transition t, convert φ(t) to quantile ranks (source-rank quantile under the source-oriented convention of Supplementary Note 7; lower rank quantile denotes higher source-oriented φ).**

**Step 5.2 — Consistency-weighted averaging.** Weight each transition by its cross-transition consistency (Spearman ρ with adjacent transitions) before averaging. This down-weights transitions associated with anomalous window compositions.

**Step 5.3 — Stable-High identification.** A gene is Stable-High if it falls in the top quantile of the φ distribution in at least 4 of 5 transitions, evaluated over 100 bootstrap iterations (bootstrap: sampling donors within each window with replacement, followed by full Stage 3–4 recomputation). The Lipschitz bound in Supplementary Note 1 provides a computable sufficient condition for rank stability of the underlying di ranking under leave-one-donor-out perturbation; the Stable-High criterion is strictly more demanding.

**Output.** Transition-averaged, KN-mode φ ranking over all p genes; Stable-High gene set (used for pathway enrichment in Results §5).

##### Implementation summary

| Stage | Key library call |
| --- | --- |

| — | — |
| --- | --- |

| Ledoit-Wolf covariance | sklearn.covariance.LedoitWolf |
| --- | --- |

| Matrix logarithm | scipy.linalg.logm |
| --- | --- |

| Diffusion map eigendecomposition | scipy.sparse.linalg.eigsh |
| --- | --- |

| Hodge decomposition (k-NN) | scipy.sparse.linalg.lsqr |
| --- | --- |

| Curl community detection | python-louvain (community package) |
| --- | --- |

The dual-mode architecture (KN for stable ranking; k-NN for signal amplification) is the minimal configuration satisfying three simultaneous requirements: temporal stability of rankings across transitions, statistical independence of curl from gradient, and data-dependent (non-trivial) gradient fraction. No single graph topology satisfies all three (Results §1, Stage 5; Methods (Five-stage Hodge pipeline, Stage 4)).

#### Supplementary Note 6: Norman Perturb-seq Covariance-Response Source-Recovery Audits

This note documents the four-tier benchmark series that supports the central methodological claim of this work: intervention-consistent structural cascade direction is recoverable from the control-whitened off-diagonal covariance-response field. The benchmarks isolate (i) the empirical equivalence of multiple structural-aggregation operators on the complete graph, (ii) the variance-decomposition contribution of the off-diagonal component, (iii) the dual-mode behaviour across complete versus sparse graphs, and (iv) the regime-specific contribution of Laplacian smoothing within the Hodge framework. All benchmarks use the same Norman et al. 2019 Perturb-seq test split (scPerturb deposition; 111,122 K562 cells, 232 perturbation groups after QC) and the same R2b pipeline (TP10K + log1p normalisation, global ZCA whitening from control cells, SPD Ledoit–Wolf log-correlation Δ matrix, bootstrap n = 100, seed = 42, control subsample = min(max(n_pert_, 100), n_ctrl_), per-perturbation Top-1 / Top-5 / Top-10 / median rank evaluation). The candidate gene universe (101 genes) is shared across all scorers within a single benchmark.

##### S6.1 Benchmark design and candidate gene universe.

The Norman Perturb-seq test split (scPerturb-harmonised; Peidli et al. 2024) defines 232 perturbation groups after the joint cell-count quality filter (n_cells ≥ 3 per group). The candidate gene set (n = 101) is the original Norman et al. 2019 perturbation-target list intersected with the gene-expression matrix; this candidate universe is reused across all scorers so that source-recovery accuracy is comparable. Per perturbation group, 100 bootstrap iterations are drawn with control subsample = min(max(n_pert, 100), n_ctrl). Within each bootstrap iteration, the whitened SPD log-correlation Δ matrix is computed once and shared across all scorers; the only difference between methods is the final aggregation operator. This shared-pipeline design ensures that differences in Top-1 reflect aggregation choice rather than preprocessing variability.

##### S6.2 Structural-family equivalence on the complete graph.

On the complete graph K_N_, four aggregation operators were computed on the same whitened Δ matrix: Hodge potential φ (computed via the K_N_ Laplacian pseudoinverse φ = L⁺ div(f), where f(i,j) = |Δ_ij_| × sign(d_i_ − d_j_) and d_i_ = ‖Δ_i,·_‖₂; reported under the source-oriented convention used throughout the main text and defined in Supplementary Note 7 (larger φ = upstream/source-like)); L1 node strength (row sum of |Δ|; ranked descending); PageRank on |Δ| (α = 0.85, power iteration with uniform teleport; ranked descending); and eigenvector centrality on |Δ| (top eigenvector of the symmetric matrix; ranked descending). L2 row norm of Δ (= d_corr_ in main text) is included as a fifth scorer for direct comparison with Table 1.

Top-1 / Top-5 / Top-10 results on 232 perturbation groups: Random ranking 0.98% / 4.9% / — (median rank 51.5); L2 / d_corr 57.3% / 79.7% / 85.8% (median rank 1.0); Hodge φ 65.9–66.4% / 81.0% / — (median rank 1.0); L1 node strength 67.7% / 84.1% / 89.7% (median rank 1.0); eigenvector centrality 67.7% / 83.2% / 90.5% (median rank 1.0); PageRank 68.1% / 84.9% / 90.1% (median rank 1.0); supervised differential-expression upper bound 81.0% / 90.9% / — (median rank 1.0).

Pairwise McNemar tests on Top-1 outcomes: L1 vs PageRank, discordant n = 3, p = 1.0 (n.s.); L1 vs Eigenvector, discordant n = 8, p = 1.0; PageRank vs Eigenvector, discordant n = 9, p = 1.0. L2 vs L1, discordant n = 26, p = 8.0 × 10⁻⁷; L2 vs PageRank, discordant n = 27, p = 4.2 × 10⁻⁷; L2 vs Eigenvector, discordant n = 26, p = 8.0 × 10⁻⁷. The three top-band scorers (L1, PageRank, Eigenvector) are therefore statistically indistinguishable at the Top-1 level, while L2 is significantly weaker than each of them by ≈ 10 percentage points.

Per-group Spearman correlations among full per-gene rankings: L1 vs PageRank, median ρ = 0.9994, IQR [0.9991, 0.9995], with 231 of 232 groups exhibiting ρ > 0.99; L1 vs Eigenvector, median ρ = 0.9973, 231 of 232 groups with ρ > 0.99; PageRank vs Eigenvector, median ρ = 0.9962, 224 of 232 groups with ρ > 0.99. Hodge φ versus each centrality measure yields median ρ ≈ −0.997 (sign-flipped relative to the raw Hodge solver output; after the source orientation defined in Supplementary Note 7 X.4 the source-score rankings are equivalent in absolute rank correlation). The structural-aggregation family is therefore empirically equivalent on K_N: the choice of aggregation operator is secondary, and the source-identification signal is encoded in the underlying whitened Δ structure shared by all four operators.

The same K_N equivalence was independently confirmed on the sALS cell-type panel. Across the ten NYGC cell types, per-CT Spearman ρ between IDS φ and each of L1 strength, L2 row norm, PageRank, and eigenvector centrality on the cell-type-specific Δ matrix ranged from −0.95 to −0.999 (median −0.988). Pairwise among the four centrality measures, per-CT ρ ranged from 0.955 to 1.000 (median 0.999; L1 ≡ PageRank to numerical precision in 10/10 cell types). The Hodge / centrality equivalence demonstrated on Norman therefore extends to the sALS application setting.

##### S6.3 Variance decomposition: separating diagonal from off-diagonal contributions.

To distinguish whitening artifacts, per-gene magnitude effects, and the genuinely off-diagonal covariance-response signal, eight scorers were compared on the same Norman test split. Four diagonal scorers preserve gene identity: raw per-gene variance ratio |log(var_pert_ / var_ctrl_)|; per-gene scale-corrected variance (control-normalised diagonal z-score); ZCA-whitened per-coordinate variance (variance ratio in the rotated coordinate system after X @ W, where coordinates are gene-aligned but linearly mixed by the whitening rotation); and control-conditioned residual variance via the Schur complement on the control precision matrix, computed as r_j_[cell] = (X[cell] · Ω_c_)[j] / Ω_c_[j,j] with Ω_c_ = Σ_c_⁻¹, which removes the linear conditional prediction of gene j from all other genes while preserving gene identity. Four off-diagonal scorers operate on the whitened Δ matrix: L2 row norm (d_corr_), L1 row sum, PageRank, eigenvector centrality, and Hodge φ.

Top-1 hit rates on 232 perturbation groups: random ranking 0.98%; raw per-gene variance 6.0%; per-gene scale correction (diagonal z) 28.4%; ZCA-whitened per-coordinate variance 18.5%; control-conditioned residual variance via Schur complement 19.8% Top-1 / 35.3% Top-5 / 63.8% Top-10 (median rank 8.0); L2 row norm 57.3%; Hodge φ / L1 / Eigenvector / PageRank centrality 65.9–68.1%; supervised DE upper bound 81.0%.

Layer-wise interpretation: raw variance to scale-corrected variance gains +22.4 percentage points (per-gene scale correction is informative on its own); scale-corrected variance to ZCA-whitened variance loses 9.9 percentage points (coordinate rotation mixes gene identities and degrades per-coordinate variance as a per-gene scorer); ZCA to Schur-complement conditional variance gains 1.3 percentage points on Top-1 and 21 percentage points on Top-10 (the diagonal-multivariate limit, where every per-gene effect that can be expressed as a change in residual variance given multivariate background is captured); conditional variance to off-diagonal centrality gains 38–48 percentage points on Top-1 (the dominant single contribution). The off-diagonal covariance-response aggregation contribution beyond diagonal and conditional-variance baselines is therefore +48 percentage points, and is not reducible to per-gene scale, per-coordinate variance, or full-multivariate-diagonal variance signals. The Schur-complement conditional variance reaches 63.8% Top-10 (median rank 8.0), indicating that gene-identity-preserving multivariate-diagonal information already identifies a top-10 candidate set of plausible perturbation sources; the off-diagonal aggregation is what disambiguates this candidate set down to Top-1.

##### S6.4 k-NN sparse-graph dual-mode validation.

The dual-mode architecture of the IDS pipeline (complete graph K_N for stable ranking and multi-transition integration; k-NN sparse graphs for per-transition signal amplification and gradient/curl decomposition) was validated empirically on Norman. k-NN graphs were constructed by row-distance of |Δ| (Euclidean distance between rows of the absolute-valued Δ matrix), retaining the top-k nearest neighbours per node and OR-symmetrising. Five scorers were computed at each k ∈ {10, 20, 50, 100}: Hodge φ on the k-NN graph (solved via LSQR on the sparse Laplacian L_knn φ = div(f); the antisymmetric flow uses the same f_ij = |Δ_ij| × sign(d_i − d_j) with d_i computed on the full Δ row, not restricted to k-NN neighbours); L1 row sum on the k-NN |Δ| adjacency; PageRank and eigenvector centrality on the k-NN |Δ| adjacency; and per-gene curl participation (aggregated Hodge-orthogonal residual curl = f − gradient).

Top-1 hit rates per k × scorer: at k = 10, Hodge φ 44.0%, L1 0.4%, PageRank 0.4%, eigenvector 1.3%, curl-per-gene 0.9%; at k = 20, Hodge φ 57.8%, L1 2.6%, PageRank 0.4%, eigenvector 5.2%, curl 0.9%; at k = 50, Hodge φ 64.2%, L1 22.0%, PageRank 18.5%, eigenvector 26.7%, curl 15.5%; at k = 100 (≈ K_N since n_candidates = 102), Hodge φ 66.4%, L1 67.7%, PageRank 68.5%, eigenvector 68.1%, curl 59.9%. At k = 100, all scorers reconverge to the K_N values within ±2 percentage points, confirming the sanity-check expectation.

Pairwise McNemar tests for Hodge φ versus each centrality measure: at k = 10, p = 2.1 × 10⁻²⁹ (vs L1), 2.1 × 10⁻²⁹ (vs PageRank), 3.2 × 10⁻³⁰ (vs Eigenvector); at k = 20, p = 1.9 × 10⁻³⁷, 6.2 × 10⁻³⁹, 1.2 × 10⁻³⁵; at k = 50, p = 1.3 × 10⁻²⁵, 9.0 × 10⁻²⁹, 1.7 × 10⁻²². At k = 100, p = 0.51, 0.23, 0.39 (all n.s.), reproducing the K_N family equivalence. The Hodge framework therefore markedly outperforms unsigned graph centrality on sparse graphs at k ≤ 50 (McNemar p ≤ 10⁻²² across all comparisons), whereas the family re-converges on dense graphs as k approaches N.

Per-gene curl participation (top-1 0.9% at k = 10, 15.5% at k = 50, 59.9% at k = 100) carries an independent edge-level signal (curl is mathematically orthogonal to gradient by Hodge decomposition), but at the per-gene Top-1 metric the curl summary collapses largely into the same ranking as φ (curl-only Top-1 wins across all k: 2 / 0 / 2 / 0 of 232 groups respectively). The hybrid φ ∪ curl Top-1 upper bound exceeds φ alone by only 0.4–0.9 percentage points. Curl carries genuine per-edge information about circular feedback structure, but per-gene aggregation does not add Top-1 source-identification capacity beyond what φ already provides.

##### S6.5 Signed-divergence baseline and Laplacian-smoothing specificity within the Hodge framework.

A potential interpretation of S6.4 is that the Hodge advantage on sparse graphs arises specifically from access to the global flow magnitude d_i_ = ‖Δ_i,·_‖₂ (computed over all genes, not only k-NN neighbours), and that the additional Laplacian inversion step (φ = L⁺ div(f)) is mathematically decorative. To distinguish these mechanisms, a signed-divergence baseline was computed on the same k-NN graph: the antisymmetric flow f_ij_ = |Δ_ij_| × sign(d_i_ − d_j_) was constructed identically to the Hodge construction, and the divergence div(f)_i_ = −Σ_j ∈ k-NN of i_ f_ij_ was used directly as a per-gene score (oriented to the same source-rank convention as Hodge φ, defined in Supplementary Note 7 X.4). The signed-divergence scorer therefore retains the global d_i_ magnitude and the signed-flow direction, but omits the Laplacian system solve.

Top-1 hit rates: at k = 10, Hodge φ 44.0% vs signed-divergence 25.4% (Δ = +18.6 pp); at k = 20, Hodge φ 57.8% vs signed-divergence 44.4% (Δ = +13.4 pp); at k = 50, Hodge φ 64.2% vs signed-divergence 64.2% (Δ = 0); at k = 100, Hodge φ 66.4% vs signed-divergence 66.4% (Δ = 0).

Pairwise McNemar tests on the 232 Top-1 outcomes: at k = 10, phi-only Top-1 wins = 43 of 232, signed-divergence-only Top-1 wins = 0, n_discordant = 43, p = 2.3 × 10⁻¹³; at k = 20, phi-only = 31, sdiv-only = 0, n_discordant = 31, p = 9.3 × 10⁻¹⁰; at k = 50, phi-only = 1, sdiv-only = 1, n_discordant = 2, p = 1.0 (n.s.); at k = 100, phi-only = 0, sdiv-only = 0, n_discordant = 0. At k = 10 and k = 20, Hodge φ achieves strict dominance over raw signed divergence: every group in which φ achieves Top-1 but the signed divergence does not is won by φ, and the converse never occurs.

Per-group Spearman correlations between Hodge φ and signed divergence rankings: at k = 10, median ρ = 0.731 (IQR [0.587, 0.797]); at k = 20, median ρ = 0.744 (IQR [0.602, 0.815]); at k = 50, median ρ = 0.797 (IQR [0.670, 0.864]); at k = 100, ρ = 1.000 exact in all 232 groups. The Laplacian inversion therefore produces a substantially different ranking from raw signed divergence at sparse k (median ρ ≈ 0.73–0.80), with the difference contributing strict Top-1 dominance for Hodge φ; at the K_N limit, the Laplacian inversion preserves rank order exactly (this follows from the algebraic structure of the K_N Laplacian: L_{K_N} = N · (I − 11^T/N) has L⁺ acting as a constant on the zero-mean subspace, which preserves the ranking induced by div(f) up to a non-rank-changing affine map).

The Hodge framework's distinctive Top-1-accuracy contribution within the sparse-graph regime is therefore specifically the Laplacian smoothing operation, which converts the locally noisy k-NN divergence into a global gradient potential via random-walk-like averaging. On dense graphs, the contribution of Laplacian smoothing to Top-1 accuracy is zero, and the Hodge framework's value shifts to its interpretive and theoretical components: gradient-curl orthogonal decomposition (main text Results §3), the analytical gradient-fraction baseline GF₀ = 2/[3(1 + CV²)] (main text Results §1 and Supplementary Note 4), the harmonic-component interpretation, and the topology-residual 3φ framework that licenses disease-specific structural interpretation (main text Results §2 and Supplementary Note 5).

##### S6.6 Failure modes, related work, and epistemic boundary.

Roughly 34% of Norman perturbation groups fail at Top-1 identification under every scorer in the structural family. Failure-mode analysis identifies four categories: diffuse effectors that distribute the perturbation signal broadly across the candidate set without a dominant source (e.g., MAPK1); indirect effectors that act through intermediate genes not in the candidate set (e.g., ARID1A); non-transcriptional effectors that change protein-level activity without a measurable mRNA covariance footprint (e.g., ATL1); and epistatically masked effectors in double-gene perturbations (e.g., AHR in AHR_KLF1_ double knockdowns). These categories define the operational boundary of intervention-consistent source recovery from covariance-response geometry.

Mathematical relationship to existing methods: Hodge decomposition on sparse pairwise comparison graphs is not mathematically unprecedented; the HodgeRank framework (Jiang, Lim, Yao & Ye 2011) establishes the use of Hodge decomposition to recover a global ranking from sparse pairwise preference data, with gradient, curl, and harmonic components interpreted respectively as ranking, intransitive inconsistency, and topological obstruction. Hodge-theoretic ideas have also entered single-cell biology in the trajectory-inference setting (PHLOWER; Cheng et al. 2025), and Riemannian / SPD covariance geometries have been applied to neurodegenerative single-cell transcriptomics (Choi et al. 2026 for Parkinson's disease). The present work draws on these foundations and is novel at a distinct methodological layer: the demonstration that experimentally imposed perturbation sources are recoverable from the off-diagonal structure of a control-whitened covariance-response field, that this recovery is mediated by a structural-aggregation family of which Hodge φ is one member, that Hodge-specific Laplacian smoothing contributes additional accuracy on sparse graphs, and that the residualisation of this validated readout against healthy co-expression topology enables disease-specific structural interpretation in sALS.

Epistemic positioning: the benchmarks in this Note establish intervention-consistent causal-source recovery — when a perturbation source is experimentally imposed, it is recovered as a structural source in the post-perturbation covariance-response field. They do not establish Pearl/Rubin causal-effect estimation. The framework does not output directed acyclic graphs, average treatment effects, or counterfactual quantities. Applied to sALS, the same validated readout identifies structural upstreamness after topology regression (main text Results §3); the resulting hypotheses about disease cascade architecture (brain-side circuit-distal oligodendrocyte upstream / motor-neuron sink; NEMF/CATylation as candidate intermediate node) are testable structural inferences, not direct evidence of chronological or molecular causation.

##### S6.7 Sources and reproducibility.

All benchmarks described in this Note were executed on the canonical IDS R2b pipeline (TP10K + log1p normalisation; Ledoit–Wolf SPD covariance estimation with eigenvalue clamping at 10⁻¹⁰; symmetric ZCA whitening W = V D⁻¹/² V^T^ from control cells; log-Euclidean tangent-space mapping; bootstrap n = 100 with replacement, seed = 42, control subsample = min(max(n_pert_, 100), n_ctrl_); per-perturbation Top-1 / Top-5 / Top-10 / median rank evaluation). Code is available at the project repository (github.com/akacola2006/scrnaseq-hodge-pipeline). The structural-family, variance-decomposition, k-NN dual-mode, and signed-divergence benchmarks reuse the identical data loader, candidate-gene universe, whitening matrix, and bootstrap protocol; they differ only in the final aggregation operator applied to the whitened Δ matrix. The Norman data are accessed via the scPerturb harmonised deposition (Peidli et al. 2024) from the original Norman et al. 2019 K562 CRISPR Perturb-seq dataset (GSE133344). Per-perturbation-group result tables (Top-1 / Top-5 / Top-10 / median rank) and per-pair McNemar / Spearman statistics are available in the project repository under the external benchmark directory.

**Supplementary Note 7: Structural causal upstreamness from covariance-response geometry**

***7.1 Scope and non-claims***

This note does not claim that graph centrality is causality. It proves a conditional statement: in a baseline-normalised covariance-response field generated by local dissipative propagation, and subject to finite-propagation admissibility, response-field hubness identifies the intervention source. In observational disease data the corresponding claim is weaker and operational: topology-residualised response-field hubness defines *structural upstreamness*, not Pearl/Rubin causal-effect estimation.

The intended hierarchy of progressively constrained readouts is

static covariance hubness
  < response-field hubness
    < baseline-normalised response-field hubness
      < finite-propagation-admissible response-field hubness
        < topology-residualised structural upstreamness.

Only the last layer licenses disease-specific biological interpretation. Throughout this Note, *finite-propagation admissibility* is used in preference to the term “light cone”: the point is not to impose Lorentzian geometry on transcriptomic data, but to require that a candidate upstream source explain downstream response within an admissible propagation domain. Section 7.6 develops this admissibility condition; Section 7.8 applies the resulting hierarchy to oligodendrocytes, translation/ribosome pathways, and NEMF using language calibrated to the strongest defensible claim. Section 7.10 contains the reviewer-facing summary.

This Note is positioned as a self-contained mathematical justification for the empirical results in the main text. Theorems A–C of the underlying chain (tangent stress displacement; canonical row engagement; unique magnitude-preserving directed graphisation) are proved in Supplementary Notes 1 and 3; this Note recapitulates their statements and develops three further results specific to the source-recovery interpretation: Theorem 7.1 (intervention-consistent source recovery), Theorem 7.2 (source recovery under finite-propagation admissibility), and Corollary 7.3 (the integrated structural-source statement). The unified statement is presented as a *corollary* rather than a theorem, because it is the operational summary of the preceding results rather than a new mathematical claim.

***7.2 Response-field setting***

Let *V* = {1, …, *n*} be a set of nodes (gene, pathway, cell type, or module, depending on the analysis). Let Σ₀ denote a baseline covariance state and Σ₁ a perturbed, disease, or later-state covariance state. In the SPD/log-Euclidean formulation, define *X*ᵥᵥ = log Σᵥᵥ and the covariance-response tangent

Δ = *X*₁ − *X*₀ ∈ ℝⁿˣⁿ, Δᵢⱼ = Δⱼᵢ.

Δ is a *tangent-space representation* of how covariance geometry changes relative to a baseline, control, perturbation, or adjacent pseudotime window. In perturbation data, Δ may be constructed after control whitening; in observational disease data, Δ is a log-Euclidean transition field between covariance windows. The diagonal Δᵢᵢ encodes marginal variance response; the structural-source readout considered here uses primarily the off-diagonal geometry:

Δ^off_ij = Δᵢⱼ (*i* ≠ *j*), Δ^off_ii = 0.

**Tangent-response origin.** Under the log-Euclidean metric on SPD(*n*), for a stress functional *S̃*(*X*) = *S*(exp *X*), the natural-gradient flow *Ẋ* = −∇*S̃*(*X*) admits the Lyapunov dissipation d*S̃*/d*t* = −‖∇*S̃*‖²_F ≤ 0 (Supplementary Note 1, Theorem 1). For a finite window, Δᵥᵥ = *X*ᵥᵥ₊₁ − *X*ᵥᵥ = −ηᵥᵥ ∇*S̃*(*X*ᵥᵥ) + O(ηᵥᵥ²), so Δ is the first-order observable tangent displacement of a stress-response dynamics. Static covariance Σ alone is symmetric and non-ordered (Σᵢⱼ = Σⱼᵢ); the source-recovery interpretation rests on covariance *change*, not covariance itself.

***7.3 Node engagement and response-field hubness***

For each node *i*, define off-diagonal response-field hubness in *L*²:

*h*ᵢ = (Σ_{*j*≠*i*} |Δᵢⱼ|²)^(1/2) = ‖Δ^off_{*i*,·}‖₂.

The *L*¹ analogue is *h*ᵢ^(1) = Σ_{*j*≠*i*} |Δᵢⱼ|. Among maps Sym(*n*) → ℝⁿ_≥0 satisfying (i) permutation equivariance *e*(*P*Δ*P*ᵀ) = *P* · *e*(Δ), (ii) row locality (*eᵢ* depends only on row *i*), (iii) positive homogeneity *eᵢ*(λΔ) = |λ| · *eᵢ*(Δ), (iv) quadratic additivity on disjoint row supports, and (v) exact Frobenius-energy partition Σᵢ *eᵢ*(Δ)² = ‖Δ‖²_F, the unique scalarisation is the row *L*² norm *eᵢ*(Δ) = ‖Δᵢ,·‖₂ (Supplementary Note 3, §3.2). The proof reduces under (i)–(iv) to a row functional *g* with *g*(λ*u*)² = λ² *g*(*u*)² and disjoint-support additivity *g*(*u*)² = Σ_k *g*(*u*_k *e*_k)²; energy fidelity (v) on single-entry matrices forces *g*(*a* *e*_k)² = *a*², hence *g*(*u*)² = Σ_k *u*_k². The row *L*² norm is therefore not an arbitrary choice but the unique local, energy-faithful, permutation-equivariant scalarisation of log-Euclidean covariance-response energy at node *i*.

The essential quantity is not marginal expression or marginal variance, but the amount of off-diagonal covariance response aggregated around node *i*. On *K*_N, multiple structural-aggregation operators (*L*¹ row sum, PageRank, eigenvector centrality, source-oriented Hodge φ) yield empirically rank-equivalent rankings of *h*ᵢ (Supplementary Note 6, §S6.2); the choice of aggregation is secondary to the underlying response-energy ordering.

***7.4 Hodge source potential on K_N***

Given Δ and *h*ᵢ = ‖Δᵢ,·‖₂, define the class of antisymmetric edge flows preserving pairwise response magnitudes:

𝒜(Δ) = { *g* ∈ ℝⁿˣⁿ_skew : |*g*ᵢⱼ| = |Δᵢⱼ| for all *i* < *j* }.

Under the alignment objective *J*(*g*) = Σ_{*i*<*j*} *g*ᵢⱼ (*h*ᵢ − *h*ⱼ), and the generic non-tie condition *h*ᵢ ≠ *h*ⱼ on every edge with |Δᵢⱼ| > 0, the unique maximiser of *J* over 𝒜(Δ) is

*f**ᵢⱼ = |Δᵢⱼ| · sign(*h*ᵢ − *h*ⱼ)

(Supplementary Note 3, Theorem 3.3). The objective decouples additively over edges; for each edge the magnitude is fixed and the optimal sign equals sign(*h*ᵢ − *h*ⱼ), and the generic non-tie condition gives global uniqueness. Because *h*ᵢ = η ‖[∇*S̃*(*X*)]ᵢ,·‖₂ + O(η²) under the natural-gradient dynamics of Section 7.2, *f** is the unique magnitude-preserving antisymmetric graphisation maximally aligned with first-order nodewise stress-response energy: *direction is not read from the symmetric entry Δᵢⱼ itself, but from the global response-energy ordering induced by hᵢ*.

The discrete Hodge decomposition writes *f** = *d*φ_raw + δψ + *h*, where *d*φ_raw is the gradient component (potential-induced), δψ the curl component, and *h* the harmonic component. The scalar potential satisfies the discrete Poisson equation *L* φ_raw = div(*f**).

On the complete graph *K*_N the harmonic component vanishes and the Laplacian pseudoinverse has the closed form *L*⁺ = (1/*N*)(*I* − *J*/*N*). Hence

φ_raw,i = (1/*N*) div_i(*f**) = (1/*N*) Σ_{*j*≠*i*} *f**ᵢⱼ.

**Source orientation.** The sign of φ_raw depends on the incidence convention chosen on the graph. To remove this ambiguity, all reported potentials are *source-oriented*: φ ≡ ±φ_raw, chosen so that larger φ denotes more source-like (upstream) position. Under this convention, the cell-type-level condition-displacement formula and the gene-level rankings reported in the main text are uniformly source-oriented. Source orientation is what licenses “upstream”/“downstream” terminology for φ-ranked nodes.

***7.5 Intervention-consistent source recovery***

We give the result in two steps: a model-free margin proposition (Proposition 7.0), then a dissipative-source theorem (Theorem 7.1) that supplies a concrete margin.

**Proposition 7.0** (margin-based source recovery, model-free). *Let s ∈ {1, …, n} be an externally imposed perturbation source and let* Δ⁽ˢ⁾ *be the corresponding control-whitened covariance-response field. Let Sᵢ(*Δ) *denote a structural-aggregation score (e.g. Hodge φ, L¹ row sum, PageRank, eigenvector centrality). Suppose the true source has a separated off-diagonal response footprint,* Sₛ(Δ⁽ˢ⁾) > max_{i≠s} Sᵢ(Δ⁽ˢ⁾) + *m, for some margin m > 0, and the bootstrap-mean estimation error satisfies* max_i |*Ŝᵢ − Sᵢ*| < m/2. *Then* ŝ = argmax_i *Ŝᵢ* = *s*.

The proposition is purely a margin statement; it commits to no propagation model. The Norman Perturb-seq benchmarks (Supplementary Note 6) empirically test the fraction of real CRISPR perturbations satisfying its premise. Theorem 7.1 below derives *m* = αγ as a structural margin from a dissipative local-source model and shows that the source uniquely maximises off-diagonal response-field aggregation, not merely that some margin happens to exist.

A dissipative local-source response model justifies reading φ as a structural source readout. Assume a local source *s* generates a dominant propagated response mode *r* = (*r*₁, …, *r*ₙ), *rⱼ* ≥ 0, with first-order response-field model

Δ = α · Off(*rr*ᵀ) + *E*, α > 0,

where Off(·) zeros the diagonal and *E* absorbs residual noise, finite-sample error, higher-order dynamics, and hidden confounding. A stable propagation model *r* = (*I* − *A*)⁻¹ *e*_s with ρ(*A*) < 1 produces this form: downstream amplitudes are attenuated and the source has the largest local amplitude, *r*_s > *r*ᵢ for *i* ≠ *s*.

**Theorem 7.1** (intervention-consistent source recovery). *Let* Δ = α · Off(*rr*ᵀ) + *E* *with* α > 0, *rⱼ* ≥ 0, *and true source* *s*. *Assume* (A1) *source dominance:* *r*_s > *rᵢ* *for all i ≠ s; (A2) multi-target propagation:* Σ_{*j*∉{*s*,*i*}} *rⱼ*² > 0 *for every i ≠ s; (A3) noise bound:* max_i ‖*Eᵢ,·*‖₂ ≤ η. *Define the noiseless row-engagement score gᵢ = rᵢ (*Σ_{*j*≠*i*} *rⱼ*²)^(1/2) *and the source margin* γ = min_{*i*≠*s*} (*g*_s − *gᵢ*). *Then* γ > 0, *and if* η < αγ/2, *the observed response-field hubness uniquely identifies the source:* *h*_s > *hᵢ* *for all i ≠ s. Under the source-oriented Hodge construction (Section 7.4), the conclusion transfers to* φ: φ_s > φᵢ *for all i ≠ s*.

*Proof sketch.* From *g*ᵢ² = *r*ᵢ² (*R*² − *r*ᵢ²) with *R*² = Σⱼ *rⱼ*², we have *g*_s² − *g*ᵢ² = (*r*_s² − *r*ᵢ²) Σ_{*j*∉{*s*,*i*}} *rⱼ*² > 0 by (A1)–(A2), so γ > 0. The reverse triangle inequality gives |*h*ᵢ − α*gᵢ*| ≤ η, hence *h*_s − *hᵢ* ≥ α(*g*_s − *gᵢ*) − 2η ≥ αγ − 2η > 0 when η < αγ/2. The Hodge conclusion follows from the *K*_N divergence relation and order-preservation of the weighted flow. ∎

**Probabilistic consistency.** Under *hᵢ* = α*gᵢ* + εᵢ with centred sub-Gaussian errors of scale σ² and effective margin Γ = αγ, a sub-Gaussian tail and union bound give Pr(*ŝ* ≠ *s*) ≤ (*n* − 1) exp(−*c* Γ²/σ²) for *ŝ* = argmax_i *hᵢ*, *c* > 0 universal. This is the matching probabilistic statement of Theorem 7.1.

**Meaning.** Under a dissipative local-source model, the source is not merely correlated with many nodes; it is the unique maximiser of off-diagonal response-field aggregation. This is the mathematical basis for interpreting response-field hubness as structural causal upstreamness. The interpretation is conditional on (A1)–(A3); the Norman Perturb-seq benchmarks (Supplementary Note 6) empirically test the fraction of real CRISPR perturbations that satisfy these conditions.

***7.6 Finite-propagation admissibility***

Theorem 7.1 establishes when a source maximises response-field hubness. Centrality alone, however, is not enough for a causal reading: a source candidate must also be able to *explain* downstream responses through finite-speed propagation. We use the term *finite-propagation admissibility* rather than a literal Lorentzian light cone. The point is not to impose relativistic geometry on transcriptomic data, but to require that a candidate upstream source explain downstream response within an admissible propagation domain.

**Definition (future cone).** Let *d*_G(*i*, *j*) be a structural distance on a graph or manifold of nodes (graph distance in a co-expression or regulatory graph, diffusion distance, anatomical/circuit distance for cell-type compartments, or a learned manifold distance). Let θᵢ be a response-time coordinate for node *i*; in pseudotime or snapshot data, an operational estimate is

θᵢ = (Σᵥᵥ tᵥᵥ *hᵢ*(*w*)) / (Σᵥᵥ *hᵢ*(*w*)),

i.e. the *h*-weighted mean activation pseudotime of node *i*. Given an effective propagation speed *c* and slack ε_θ, define the source future cone

𝒞_c^+(*i*) = { *j* : θⱼ ≥ θᵢ − ε_θ *and* *d*_G(*i*, *j*) ≤ *c* (θⱼ − θᵢ) + *c* ε_θ }.

A candidate source *s* is finite-propagation-admissible for downstream node *j* only if *j* ∈ 𝒞_c^+(*s*).

**Cone-restricted hubness.** Define a cone mask *M*^(c)_ij = 𝟙{*j* ∈ 𝒞_c^+(*i*)} (a soft kernel version is also valid; see the source markdown for details). The cone-restricted hubness is

*h*ᵢ^c = (Σ_{*j*≠*i*} *M*^(c)_ij |Δᵢⱼ|²)^(1/2).

**Theorem 7.2** (source recovery under finite-propagation admissibility). *Under the model of Theorem 7.1, assume* (C1) *finite-propagation support: rⱼ > 0, j ≠ s* ⇒ *j* ∈ 𝒞_c^+(*s*); (C2) *conic source margin* γ_c = min_{*i*≠*s*} (*g*_s^c − *gᵢ*^c) > 0 *with gᵢ*^c = *rᵢ* (Σ_{*j*≠*i*} *M*^(c)*ij rⱼ²)^(1/2); (C3) cone-restricted noise bound max_i (Σ*{*j*≠*i*} *M*^(c)_ij |*Eᵢⱼ*|²)^(1/2) ≤ η_c. *If* η_c < α γ_c / 2, *then h*_s^c > *hᵢ*^c *for all i ≠ s*.

*Proof.* Identical to Theorem 7.1 after replacing the full row norm with the masked row norm. The masked noiseless field satisfies ‖*M*^(c)*i,· ⊙ Δ⁰*{*i*,·}‖₂ = α *gᵢ*^c, the margin assumption gives *g*_s^c − *gᵢ*^c ≥ γ_c, and the reverse triangle inequality completes the bound. ∎

**Interpretation.** The admissibility theorem adds the rigorous content of the cone analogy: *hubness is not causal upstreamness unless the hub can reach the downstream response through an admissible finite-propagation cone*. The cone-restricted hubness *hᵢ*^c can be inserted into the same Hodge construction (Section 7.4) to yield a cone-restricted source potential; this is the cone-Hodge potential. In the present snapshot ALS data, *c* and ε_θ are not separately identifiable, so the finite-propagation condition functions as an *admissibility principle*: it constrains which causal narratives are compatible with the response geometry, rather than fitting a physical propagation-speed model.

***7.7 Disease-specific residualisation***

Raw φ ranking on *K*_N is dominated by healthy co-expression topology: in the sALS cohort, 82–92 % of φ variance in glia reflects the weighted node strength of each gene in the healthy network. Disease-specific structural information therefore does not reside in raw φ position but in the *residual* after topology regression — the 3φ framework, in which φ_disease is regressed on φ_static via a cubic polynomial fit and per-gene residual *z*-scores identify rewiring (*z* > +2) and collapse (*z* < −2) (Methods §6.15).

The structural rationale for residualisation can be stated at the field level. Define the *non-integrable residual* of an edge flow as

*r*_G(*f*) = (*I* − *P*_{grad,G}) *f* = δψ + *h*.

Equivalently, the residual class [*f*]*G ∈ C¹(G) / im d is the quotient by integrable gradient deformation. Under coarse-graining maps C^1*{ℓ→L}: *C*¹(*G*_ℓ) → *C*¹(*G*_L) satisfying gradient admissibility *C*^1_{ℓ→L}(im *d*_ℓ) ⊆ im *d*_L, the macroscopic non-integrable residual depends only on the microscopic non-integrable residual:

*R*_L *C*^1_{ℓ→L} *f*_ℓ = *R*_L *C*^1_{ℓ→L} *R*_ℓ *f*_ℓ.

Integrable gradient-like deformation is annihilated under admissible coarse-graining, while the non-integrable residual class is the object that survives scale change. This is a structural — not statistical — argument: it motivates residualisation as the operation required to separate topology-dominated integrable structure from disease-specific structural displacement, and supports the 3φ framework as a mathematically natural rather than ad hoc choice. It does not prove that any individual residual gene (e.g. NEMF) is causal for ALS; the residual structure is necessary for disease interpretation but not sufficient for causal proof.

***7.8 Consequence for sALS***

Combining Theorem 7.1, Theorem 7.2, and the residualisation rationale of Section 7.7 yields safe operational language for the three principal sALS findings. In each case the strongest defensible claim is *intervention-consistent structural upstreamness*, not biological initiation of disease.

**Oligodendrocytes.** Oligodendrocytes occupy an intervention-consistent, topology-preserved structural source position in the sALS covariance-response architecture (Results §3, §7). They retain healthy manifold structure while sitting at the source side of the disease response geometry; under Theorem 7.1, this is the same structural position occupied by experimentally imposed intervention sources in the Norman benchmarks (Supplementary Note 6). The defensible reading is *a central glial upstream-like compartment, not a definitive causal origin*; the avoided reading is “oligodendrocytes are the causal origin of sALS.”

**Cytoplasmic translation / ribosome.** Cytoplasmic translation is the only cross-cell-type programme showing reproducible topology-residualised upstream enrichment after the 3φ residual transformation (Results §5; 9–10 / 10 cell types under matched-null *z*-scores). This is consistent with a *structural stress core* in the response-field sense: translational programmes remain structurally upstream after residualisation against healthy topology, identifying translational stress as the strongest cross-cell-type structural upstream programme detected by the framework. The defensible reading is *topology-residualised structural upstream programme and high-priority perturbation target*; the avoided reading is “translation dysfunction causes sALS.”

**NEMF / CATylation.** NEMF shows cross-cell-type disease-specific collapse of co-expression coherence (*z* < −2 in 7/10 cell types; Results §6) despite largely unchanged mRNA expression. In response-field language, NEMF is a *structural failure node* rather than necessarily the earliest *source* node: it may sit downstream of translational stress but upstream of TDP-43 pathology, as a bottleneck in ribosome-associated quality-control coherence. The defensible reading is *NEMF/CATylation network decoherence is a candidate structural intermediate linking translational stress to motor-neuron TDP-43 vulnerability*; the avoided reading is “NEMF loss causes TDP-43 pathology in sALS.”

These three reading rules — *upstream-like preserved compartment* for oligodendrocytes, *structural stress core* for translation, *structural intermediate* for NEMF — are the strongest mathematically defensible biological claims supported by the response-field source-recovery framework on observational sALS data.

***7.9 Relation to IDS Core Theory***

The relationship to the IDS Core Theory framework can be stated sharply. The present source-recovery note concerns the *gradient/source component* of a covariance-response field, whereas IDS Core Theory identifies the *non-integrable Hodge residual class* (curl + harmonic) as the object transported under admissible coarse-graining. These viewpoints are complementary rather than contradictory: the gradient component provides source/sink semantics at a fixed observational scale, while the residual class controls what survives reparametrisation and scale change. Disease-specific interpretation therefore requires both — source-oriented gradient potential for upstream localisation, and topology/residual controls (the 3φ framework, the curl/harmonic projection, sparse-graph residuals) to avoid confusing baseline hubness with disease-specific structural organisation. Combining finite-propagation admissibility (Section 7.6) with the non-integrable residual class via *causal-cone-admissible coarse-graining* (gradient admissibility *and* preservation of cone reachability under coarse-graining) is the cleanest way to connect finite-propagation source recovery, Hodge residuals, and disease response geometry across scales without invoking physical Lorentzian spacetime.

***7.10 Reviewer-facing summary***

The chain reads:

1. Static covariance is symmetric and non-causal (Section 7.2).
2. A perturbation or disease transition produces a log-Euclidean covariance-response tangent Δ; under natural-gradient stress dynamics, Δ is the first-order observable tangent displacement (Section 7.2, Supplementary Note 1).
3. The unique canonical nodewise response energy is the row *L*² norm *hᵢ* = ‖Δᵢ,·‖₂ (Section 7.3, Supplementary Note 3).
4. The unique magnitude-preserving directed graphisation aligned with this engagement is *f**ᵢⱼ = |Δᵢⱼ| · sign(*hᵢ* − *hⱼ*); Hodge decomposition turns it into a source/sink potential (Section 7.4).
5. Under source dominance, multi-target propagation, and a noise bound, the response-field hubness uniquely identifies the intervention source (Theorem 7.1, Section 7.5).
6. Under additional finite-propagation admissibility, the cone-restricted hubness identifies the source (Theorem 7.2, Section 7.6).
7. Disease-specific structural information requires topology residualisation (Section 7.7; the 3φ framework).

**Corollary 7.3** (integrated structural-source statement). *Let* Δ *be a baseline-normalised covariance-response field on nodes V, and assume jointly (i) the tangent-response condition (Section 7.2); (ii) the local dissipative source model of Theorem 7.1 with source-dominance and positive source margin; (iii) the noise-margin condition* η < αγ/2; *(iv) finite-propagation admissibility in the sense of Theorem 7.2 (downstream response inside the source future cone, or cone-restricted hubness with positive conic margin); (v) the source-oriented Hodge construction of Section 7.4; and (vi) topology residualisation against healthy co-expression structure (3φ framework, Section 7.7). Then the maximal residual response-field source score recovers s as the structural causal source in the operational sense*

*ŝ* = argmax_i [ φ^c_disease,i − *m̂*^c(φ^c_static,i) ] = *s*,

*provided the total residual / topology / noise error is below the combined source margin. Under sub-Gaussian errors, recovery obeys an exponential margin bound* Pr(*ŝ* ≠ *s*) ≤ (*n* − 1) exp(−*c* Γ²_eff / σ²_eff), *where* Γ_eff *is the post-cone, post-topology source margin*.

This Corollary is the operational summary of Theorems 7.1–7.2 together with the residualisation rationale of Section 7.7; it is stated as a corollary rather than a standalone theorem because each ingredient is proved above. Its content does *not* assert that observational sALS data prove the ultimate biological cause of disease. It asserts that *if* a disease-associated node occupies the same baseline-normalised, finite-propagation-admissible, topology-residualised response-field position that known CRISPR intervention sources occupy in benchmark perturbations (Supplementary Note 6), *then* it is mathematically justified to call that node an *intervention-consistent structural causal source*.

**Short reviewer answer.** *Is this causal?* It is intervention-consistent on perturbation data and structurally upstream-like on observational disease data; it does *not* estimate Pearl/Rubin causal effects, and does *not* output directed acyclic graphs, average treatment effects, or counterfactual quantities. Applied to sALS, the framework identifies oligodendrocytes as an *upstream-like preserved compartment*, cytoplasmic translation as a *topology-residualised structural stress core*, and NEMF/CATylation as a *candidate structural intermediate* — falsifiable structural inferences about disease cascade architecture, not direct evidence of chronological or molecular causation.

### Supplementary Discussion S1: ATM Brake Loss and the NVU Trigger Hypothesis

This Supplementary Discussion presents findings from an independent analytical pipeline applied to the same NYGC ALS Consortium dataset used in the main paper. It is included as a working hypothesis rather than as a primary finding of this manuscript, because the evidence base is more limited than that for the cell-type and gene-level structural results in the main text: sample sizes for vascular cell types are small (hundreds of cells), the causal inference methods employed are observational, and no interventional experiments have been performed. All causal language in this section denotes candidate mechanisms and working hypotheses rather than established relationships. The convergence of two independent pipelines on overlapping conclusions (oligodendrocyte upstream positioning) provides cross-methodological support but does not substitute for experimental validation. The ATM brake loss / NVU trigger candidate has not been independently validated in the cross-cohort meta-analysis reported in main-paper Results §7 (which used the seven common broad cell types across cohorts, not the vascular subtypes analysed here).

#### S1.1 Module-level causal analysis: independent pipeline (original §9)

##### Pipeline overview

Parts I and II analysed the NYGC ALS Consortium snRNA-seq data (primary motor cortex (BA4); 54 donors total, of which 24 were retained for primary analysis: 14 sALS + 10 PN; see Part I Methods) through a gene-level Hodge-decomposition framework applied to 10 major cell types. Part III applies an entirely separate analytical pipeline to the same underlying dataset but includes the full donor panel and extends to 43 cell types (including vascular populations not represented in Part I's 10-cell-type analysis) and 23 predefined functional modules.

The module-level pipeline proceeds as follows:

- *Module scoring.* For each of 43 cell types, expression scores are computed for 23 functional modules: Angiogenesis, Apoptosis, Autophagy, Calcium Signaling, Cell Cycle, Complement, Cytoskeleton, DNA Repair, ECM, ER Stress, Epigenetic, Growth Factors, Inflammation, Ion Transport, Metabolism, Mitochondria, Myelination, Oxidative Stress, Protein Homeostasis, RNA Processing, Synaptic, Transcription, and lncRNA.
- *Diffusion Pseudotime (PT_dpt).* A continuous disease-progression axis is constructed via diffusion pseudotime on the 23-module feature space (standardised module scores as input, k = 30 nearest-neighbour graph, Gaussian kernel, eigenvectors 2—6 of the normalised Laplacian). The root cell is defined as the cell with the lowest total stress score. PT_dpt is constructed to be stress-independent: the variance in PT_dpt explained by aggregate stress is only R-squared = 1.9%, and PT_dpt shows near-zero correlation with the stress-based pseudotime axis PT_imes (R-squared = 62.5% stress dependence). This separation ensures that ordering along PT_dpt reflects structural disease progression rather than a trivial recapitulation of stress magnitude.
- *Module-level causal inference.* Causal ordering among modules within and across cell types is estimated using LiNGAM (Linear Non-Gaussian Acyclic Model) and complementary time-lag analyses. These methods identify candidate directional relationships from observational data; they do not establish causality in the interventionist sense.

This pipeline is methodologically independent of the Hodge-decomposition approach in Part I. The two pipelines share the same raw data but differ in feature representation (gene-level φ scores vs. module-level aggregate scores), pseudotime construction (SPD manifold vs. diffusion pseudotime), and analytical framework (Hodge decomposition vs. LiNGAM-based causal estimation). Where the two pipelines converge on consistent findings, the convergence functions as a form of cross-methodological validation.

##### NVU-origin candidate pattern

The module-level pipeline revealed a consistent pattern in which vascular cell types of the neurovascular unit (NVU) exhibited early changes along the disease-progression axis. Key observations include:

In the Phase 8 time-lag analysis, upper motor neuron changes preceded glial changes across all 23 functional modules, a pattern consistent with top-down propagation but not distinguishing between neuronal vulnerability and upstream vascular perturbation. In Phases 9-11, when vascular cell types (capillary endothelium, pericytes) were incorporated into the causal hierarchy, they consistently appeared at early positions along the disease-progression axis. A recurrent pattern emerged in which capillary endothelial changes preceded pericyte changes across multiple functional modules, consistent with (but not proof of) a capillary-origin, pericyte-propagation sequence.

The capillary-first pattern emerged in module-level causal estimates: capillary endothelium occupied the earliest position among vascular cell types for DNA Repair, RNA Processing, and Oxidative Stress modules. Pericyte changes in these same modules appeared at later pseudotime positions. This ordering is consistent with a candidate model in which vascular perturbation initiates in the endothelial compartment and propagates to mural cells, though the observational data cannot exclude simultaneous or reverse-direction effects.

##### Oligodendrocyte positioning across pipelines

A point of convergence between the Part I and Part III pipelines concerns oligodendrocytes:

- In Part I, oligodendrocytes were identified as the cell type with the highest structural precedence (φ-based upstream ranking) among the 10 major neuronal and glial cell types analysed.
- In Part III, the module-level causal analysis similarly placed oligodendrocytes in an upstream position relative to other glial and neuronal populations.
- However, Part III additionally analysed vascular cell types (capillary endothelium, pericytes) that were not included in the Part I analysis. When these vascular populations were incorporated, capillary endothelium occupied a position upstream of oligodendrocytes in several module-level causal estimates.

The absence of vascular cell types from Part I reflects a methodological constraint rather than a biological conclusion: the Part I analysis focused on the 10 principal neuronal and glial cell types in the primary motor cortex (BA4) snRNA-seq data. Capillary endothelial cells and pericytes, being relatively rare in cortical snRNA-seq preparations, were analysed separately in the Part III pipeline. The candidate ordering (capillary endothelium upstream of oligodendrocytes upstream of other glia) is a working hypothesis that requires validation in datasets where all cell types can be jointly analysed.

#### S1.2 ATM pathway decline in capillary endothelium (original §10)

##### Identification criteria

To evaluate candidate early vascular signals within capillary endothelium, we applied three criteria designed to distinguish robust early signals from downstream or inconsistent changes. These criteria are necessary but not sufficient conditions for causal involvement; satisfying all three identifies a candidate mechanism warranting experimental follow-up, not a proven cause.

The first criterion is temporal ordering: the candidate mechanism must show detectable changes at earlier disease-progression stages than its proposed downstream consequences. Operationally, this requires that the candidate's expression changes precede downstream module changes along the PT_dpt axis, and that the candidate shows altered expression in the earliest pathological stage (TDP-43-negative cases in the GSE212630 dlPFC dataset; Wang et al. 2023).

The second criterion is cross-context consistency: the direction of change must be consistent across independent observation conditions. We require a sign consistency score of at least 80% across combinations of datasets (NYGC ALS Consortium and GSE212630), comparison types (case-control and pseudotime trajectory), cell types (capillary and pericyte), and analytical levels (cell-level and donor-level). The 80% threshold is set a priori to be stringent enough to exclude noisy candidates while acknowledging that biological heterogeneity may produce occasional discordance.

The third criterion is pathway coherence: the candidate must show coordinated directional change across the genes constituting its biological pathway, not merely a single-gene effect. We require at least 70% of pathway member genes to change in the same direction, a threshold that balances the expectation of coordinated pathway regulation against the reality that individual genes may be subject to compensatory or context-dependent regulation.

##### ATM pathway evaluation

The ATM-mediated DNA damage response (DDR) pathway emerged as the leading candidate early vascular signal in capillary endothelium. Its performance against the three identification criteria is summarised below:

**Table 15. Criteria-based evaluation of the ATM pathway candidate.**

| Criterion | Result | Assessment |
| --- | --- | --- |
| Temporal ordering | ATM pathway genes decline in TDP-43-negative (earliest-stage) cases in GSE212630; decline at low PT_dpt values in the NYGC dataset | Satisfied |
| Cross-context consistency | Consistent downregulation across 2 datasets, multiple comparison axes, and 2 vascular cell types: 8/8 observation conditions show the same direction (100%) | Satisfied |
| Pathway coherence | 8 of 9 ATM pathway genes (89%) show concordant downregulation | Satisfied |
| Statistical significance | ATM: p = 0.018 (ALS vs. Control, donor-level); ATM: p = 0.003 (PT_dpt trajectory, Wilcoxon) | Significant |
| Cross-dataset concordance | 83% direction match between GSE212630 and NYGC Phase 13 across 18 genes (ATM + R-loop pathways) | Concordant |

The ATM pathway satisfied all three candidate criteria and additionally showed statistically significant individual gene effects and cross-dataset concordance. On the basis of these results, ATM brake loss in capillary endothelium emerged as the leading candidate early vascular signal in this dataset. This designation reflects the observational evidence; it does not constitute proof of causality.

##### Gene-level evidence

The table below presents gene-level evidence for ATM pathway involvement in capillary endothelium. Sample sizes: Capillary, 881 cells total (785 with PT_dpt assignment); ALS 17 donors, Control 15 donors. All p-values are from the original analyses and have not been adjusted for multiple comparisons within this table; the reader should interpret individual gene-level significance in the context of the pathway-level pattern rather than as independent confirmatory tests.

**Table 16. ATM pathway gene-level evidence in capillary endothelium.**

| Gene | GSE212630 direction | Phase 13 ALS vs. Control direction (p-value) | Phase 13 PT_dpt correlation direction (p-value) | Consensus direction |
| --- | --- | --- | --- | --- |
| ATM | Down | Down (p = 0.018*) | Down (p = 0.003*) | Down |
| NBN | Down | Down (p = 0.238) | Down (p = 0.048*) | Down |
| XRCC5 | Down | Down (p = 0.126) | Down (p = 0.008*) | Down |
| XRCC6 | Down | Down (p = 0.970) | Down (p = 0.040*) | Down |
| MRE11 | Down | Down (p = 0.378) | Down (p = 0.287) | Down |
| RAD50 | Up | Up (p = 0.985) | Down (p = 0.304) | Mixed |
| TP53BP1 | Down | Down (p = 0.268) | Down (p = 0.411) | Down |
| BRCA1 | Down | Down (p = 0.165) | Down (p = 0.625) | Down |

Of the 8 genes with unambiguous consensus direction, 7 of 7 show downregulation (ATM, NBN, XRCC5, XRCC6, MRE11, TP53BP1, BRCA1). RAD50 shows mixed directionality, with upregulation in the case-control comparison but downregulation along the PT_dpt trajectory. CHEK2 was excluded from this table due to incomplete cross-dataset coverage (not available in PT_dpt analysis), though it showed downregulation in Phase 13 ALS vs. Control (logFC = -0.47).

Four genes reached statistical significance in the PT_dpt trajectory analysis (ATM, NBN, XRCC5, XRCC6; all p < 0.05), and ATM itself reached significance in the ALS vs. Control donor-level comparison (p = 0.018). The convergence of pathway-level coherence with individual gene-level significance supports the candidacy of ATM brake loss as a coordinated pathway-level phenomenon rather than a single-gene artefact.

A parallel analysis of 9 R-loop surveillance genes (SFPQ, UPF1, XRN2, DIS3, EXOSC10, RNASEH1, RNASEH2A, SETX, DDX5) showed 7/9 genes with downward direction in ALS vs. Control (pathway coherence 78%), with XRN2 (p = 0.043) and DDX5 (p = 0.047) individually significant. This is consistent with the working hypothesis that ATM brake loss may contribute to R-loop accumulation through impaired double-strand break repair, though the directional relationship between these two pathways cannot be established from observational data alone.

##### Comparison with competing candidates

To assess the specificity of the ATM pathway finding, we evaluated alternative candidate early vascular signals using the same three criteria:

**Table 17. Candidate early vascular signals ranked by criterion satisfaction.**

| Rank | Candidate | Temporal ordering | Cross-context consistency | Pathway coherence | Overall assessment |
| --- | --- | --- | --- | --- | --- |
| 1 | ATM pathway | Satisfied | 100% | 89% (8/9 genes) | Leading candidate |
| 2 | R-loop surveillance | Satisfied | 85% | 78% (7/9 genes) | Candidate (possibly downstream of ATM) |
| 3 | ATR pathway | Partially satisfied | 60% | 50% | State variable, not origin candidate |
| 4 | DAMP signalling | Not satisfied | Low | N/A | Result indicator, not origin candidate |

The ATM pathway outperformed all competing candidates across all three criteria. R-loop surveillance ranked second and may represent a downstream consequence of ATM brake loss rather than an independent origin, a possibility consistent with the known biological relationship between ATM-mediated DDR and R-loop resolution, but not demonstrable from these data. The ATR pathway showed inconsistent directionality (60% consistency), compatible with its known role as a state-responsive checkpoint rather than an initiating event. DAMP signalling failed the temporal ordering criterion entirely: DAMP levels showed strong quality-control dependence in single-cell data, and residualisation for technical covariates substantially attenuated DAMP-disease associations, suggesting that DAMP elevation reflects downstream cellular damage rather than an upstream trigger.

##### Working hypothesis: Mechanistic Model Candidate 1 (MMC-1)

On the basis of the evidence presented in Sections 10.1—10.4, we propose the following working hypothesis for the earliest detectable molecular events in ALS-associated neurovascular pathology. We designate this Mechanistic Model Candidate 1 (MMC-1) to emphasise its status as a falsifiable hypothesis rather than an established mechanism.

The MMC-1 cascade, as a working hypothesis, proceeds as follows:

ATM brake loss in capillary endothelium (candidate origin)

v

Delayed double-strand break (DSB) response

v

R-loop accumulation (DNA-RNA hybrids)

v

RNA surveillance failure

v

Propagation to downstream cell types

Each arrow in this cascade represents a candidate directional relationship supported by the temporal ordering of expression changes along the disease-progression axis. The arrows do not represent proven causal relationships. Specifically:

- The ordering of ATM decline before R-loop surveillance decline is consistent with ATM brake loss contributing to R-loop accumulation, but the reverse direction (R-loop accumulation causing ATM pathway exhaustion) cannot be excluded from observational data.
- The propagation from capillary endothelium to pericytes is inferred from the consistent capillary-first pattern observed across multiple modules (Supplementary Discussion S1, §S1.2), but simultaneous perturbation of both cell types by a common upstream cause remains possible.
- The connection between vascular perturbation and broader neurodegeneration (propagation to downstream cell types) is the most speculative element of MMC-1 and is supported primarily by the temporal ordering in the module-level causal hierarchy rather than by direct mechanistic evidence.

Falsification criteria for MMC-1 include: (i) demonstration that ATM pathway function is intact in capillary endothelium of ALS patients using orthogonal methods (e.g., immunohistochemistry, functional DDR assays); (ii) demonstration that R-loop accumulation precedes ATM pathway changes in a time-resolved model system; (iii) failure of ATM inhibition in endothelial cells to produce downstream R-loop or RNA surveillance phenotypes in vitro.

##### Sample size constraints and statistical power

The vascular cell type analyses in Part III are based on substantially smaller sample sizes than the neuronal and glial analyses in Part I:

**Table 18. Sample sizes for vascular cell type analyses.**

| Cell type | Total cells | Cells with PT_dpt | ALS donors | Control donors |
| --- | --- | --- | --- | --- |
| Capillary endothelium | 881 | 785 | 17 | 15 |
| Pericyte | 511 | 461 | 17 | 16 |

For comparison, the Part I analyses utilised 24 retained donors (14 sALS + 10 PN) across 10 major cell types, with typically several thousand cells per cell type. The vascular cell analyses in Part III therefore operate with approximately one-fifth to one-tenth the cellular sample size and fewer donors.

These sample sizes impose several constraints on interpretation:

- With 881 capillary cells and 32 total donors, the analysis is adequately powered to detect large effect sizes (Cohen's d > 0.8) but may miss moderate effects. The significant p-values reported (e.g., ATM p = 0.018, XRCC5 p = 0.008) should be interpreted in light of this power limitation: non-significant genes may harbour real effects that the current sample size cannot detect.
- The donor-level comparisons (17 ALS vs. 15 Control for capillary; 17 ALS vs. 16 Control for pericyte) have limited degrees of freedom. Wilcoxon rank-sum tests with n = 32 total can detect large effects but are underpowered for the moderate effects typical of complex disease genomics.
- The gene-level p-values in Table 16 are not corrected for multiple comparisons. While the pathway-level coherence (89% concordant direction) provides a form of internal replication that mitigates the multiple comparison concern, the individual gene-level p-values should not be interpreted as definitive evidence of differential expression.
- Diffusion pseudotime estimates for 785 capillary cells carry substantial uncertainty, particularly at the extremes of the trajectory. The correlation between gene expression and PT_dpt may be inflated or attenuated by pseudotime estimation error.
- The vascular cell data derive from a single brain region (primary motor cortex (BA4)) in a single cohort (NYGC ALS Consortium). The cross-dataset validation with GSE212630 (dlPFC, BA9, C9orf72 ALS/FTD; Wang et al. 2023) provides some evidence of cross-region, cross-disease reproducibility, but generalisability to sALS motor cortex and other brain regions (e.g. spinal cord) remains to be tested. The 83% concordance rate means that 17% of genes showed discordant directions, which may reflect genuine regional heterogeneity or noise.

These limitations do not invalidate the ATM brake loss candidacy: the convergence of multiple lines of evidence across datasets, cell types, and analytical approaches provides meaningful support for the hypothesis. However, they mandate that ATM brake loss be treated as a candidate requiring independent validation rather than an established finding. Larger vascular cell datasets, ideally from multiple cohorts and brain regions, are needed to confirm or refute this hypothesis.

#### S1.3 Limitations of the NVU/ATM analysis (original §11)

The following limitations apply specifically to the analyses and interpretations presented in Part III. These should be considered in addition to the general limitations discussed in Part I.

- The capillary endothelium (881 cells) and pericyte (511 cells) analyses are based on cell counts one to two orders of magnitude smaller than those typical of the major neuronal and glial populations in the same dataset. This limits statistical power, increases vulnerability to outlier effects, and constrains the complexity of models that can be reliably estimated. The ATM brake loss hypothesis, while consistent across multiple analytical approaches, rests on a cellular evidence base that is limited by single-cell genomics standards.
- The 23 functional modules used in the module-level causal analysis were defined prior to analysis on the basis of known biological pathways. This approach has the advantage of biological interpretability but the disadvantage of potentially missing disease-relevant gene programmes that do not correspond to canonical pathways. A data-driven module discovery approach (e.g., WGCNA or topic modelling) might identify additional or alternative functional units relevant to disease progression. The current findings are conditional on the chosen module definitions.
- The LiNGAM-based causal estimates and time-lag analyses used in Part III assume specific statistical properties (non-Gaussianity of residuals for LiNGAM; stationarity for time-lag analysis) that may not hold in single-cell transcriptomic data. More fundamentally, all causal inference from observational data is subject to the possibility of unmeasured confounders. The causal orderings reported here (including the capillary-first pattern and the ATM-before-R-loop sequence) are candidates consistent with the data, not proven causal relationships. LiNGAM-based estimates are known to be optimistic (i.e., to infer directional relationships with higher confidence than warranted) when sample sizes are small relative to the number of variables, and this concern applies to the vascular cell analyses with hundreds of cells and 23 module dimensions.
- The primary dataset (NYGC ALS Consortium) derives from primary motor cortex (BA4), while the cross-validation dataset (GSE212630; Wang et al. 2023) derives from dorsolateral prefrontal cortex (dlPFC, BA9) in C9orf72 ALS/FTD. The 83% cross-dataset direction concordance is encouraging, but the 17% discordance may reflect genuine biological differences between brain regions and disease subtypes rather than noise. Motor cortex and prefrontal cortex differ in cellular composition, vulnerability to ALS pathology, and vascular architecture. The ATM brake loss candidate hypothesis may not generalise equally to all brain regions.
- Even if ATM brake loss in capillary endothelium is confirmed as an early event in ALS pathogenesis, the cause of this ATM decline remains unknown. Possibilities include genetic variation in ATM pathway genes (rare variants or common polymorphisms), epigenetic silencing (e.g., promoter methylation of ATM or its regulators), environmental exposures affecting DDR capacity, or age-related decline in DDR function that is accelerated in ALS. The current data cannot distinguish among these possibilities. Identifying the upstream cause of ATM brake loss, should it be validated as a genuine early event, will require genetic, epigenetic, and environmental studies that are beyond the scope of transcriptomic analysis.
- All findings in Part III are based on mRNA expression levels. mRNA abundance is an imperfect proxy for protein levels and an even more imperfect proxy for protein function. ATM kinase activity, for example, is regulated by post-translational modifications (autophosphorylation, acetylation) that are invisible to transcriptomic analysis. The observation that ATM mRNA is reduced in ALS capillary endothelium is consistent with reduced ATM function but does not demonstrate it. Protein-level (immunohistochemistry, Western blot) and functional (DDR activity assays) validation in human brain vascular tissue is required.
- The capillary endothelial cells analysed here were identified by computational cell type annotation. If ALS alters the subtype composition of capillary endothelium (e.g., selective loss of ATM-high endothelial subtypes), the observed ATM decline could reflect compositional change rather than per-cell downregulation. Single-cell resolution partially mitigates this concern, but subtypes below the resolution of the clustering algorithm could contribute to the signal.

*Part III presents ATM brake loss in capillary endothelium as the leading candidate early vascular signal identified by the module-level causal pipeline. This designation is based on satisfaction of three predefined criteria (temporal ordering, cross-context consistency, pathway coherence) and cross-dataset concordance of 83%. The hypothesis is falsifiable and requires validation in independent cohorts, at the protein level, and ultimately through interventional experiments. The integration of Part III findings with the Hodge-decomposition results of Parts I—II is undertaken in Part IV.*

### Supplementary Discussion S2: Hypothetical Integration — A Multi-Layer Cascade Model for sALS

This Supplementary Discussion is the most speculative section of the supplementary material. It attempts a hypothetical integration of three independent lines of evidence: the cell-type and gene-level structural findings reported in the main paper (cytoplasmic-translation core and NEMF coupling collapse) and the candidate ATM/NVU pathway described in Supplementary Discussion S1, each of which rests on a different level of empirical support. The cascade model proposed below is an optional interpretive framework: readers may accept the main-paper findings while rejecting this integrated model without contradiction. Every connecting step is annotated with its evidential basis, and steps that rely on inference rather than data are marked explicitly. The model is presented because it generates falsifiable predictions (§S2.6), not because we regard it as established.

#### S2.1 Three independent lines of evidence (original §12.1)

This section synthesises findings from Parts I—III. Before proposing any integration, we summarise each Part's principal conclusions and their epistemic standing.

*Part I (Hodge instrument; high confidence: instrument externally validated).* Translation-associated programmes are structurally upstream in 9/9 cell types, oligodendrocytes are the sole upstream cell type (*φ* = 0.900), and disease structure decomposes into an irreversible translation-resource gradient on cyclic synaptic curl. Results are robust to gene-selection method (Spearman ρ = 0.992).

*Part II (Gene-level φ and the RQC vulnerability hypothesis; moderate confidence: exploratory, primary discovery in NYGC with predominantly negative cross-cohort support for the §8.9 NEMF finding in Part V, literature-consistent). Gene-level φ analysis reveals cell-type-specific upstream profiles centred on proteostasis and lipid metabolism in oligodendrocytes and on calcium homeostasis and axonal transport in neurons (Results §5). Among known ALS-causative genes, TBK1 and SQSTM1 rank most upstream while TARDBP ranks downstream in oligodendrocytes, consistent with TDP-43 as a convergence point. The RQC system is a candidate vulnerability node: no demand-responsive upregulation, self-limiting negative feedback, and competition with the ISR for shared substrates (Results §6).*

*Part III (ATM brake loss and the NVU trigger; limited confidence: small sample, cross-dataset validated candidate).* An independent module-level causal analysis identifies ATM-mediated DNA damage response decline in capillary endothelium as the top-ranked causal candidate (8/9 pathway genes concordant; cross-dataset agreement 83%; ATM *p* = 0.018). Capillary changes precede pericyte involvement, suggesting a neurovascular-unit origin for the cascade, constrained by small vascular cell numbers.

#### S2.2 The connecting mechanism: R-loop to ribosome stalling (original §12.2)

The three lines of evidence described above were obtained through independent analytical pipelines. Whether these three lines of evidence can be connected into a coherent sequence is the central question of this section. We propose that the following molecular logic provides a plausible, though not directly demonstrated, bridge between the NVU trigger (Part III) and ribosome quality control overload (Part II).

The proposed connection proceeds through four steps. First, ATM brake loss permits R-loop accumulation: ATM kinase is required for efficient resolution of R-loops (three-stranded DNA:RNA hybrid structures) at sites of transcription-replication conflict, and when ATM activity declines, R-loops persist and expand (Crossley et al., *Mol Cell*, 2019; Shanbhag et al., *Cell*, 2010); Part III data are consistent with early ATM pathway decline in capillary endothelium, raising the possibility that R-loop burden increases in NVU cells. Second, persistent R-loops disrupt normal transcription termination, producing read-through transcripts and mRNAs with abnormal secondary structures (Skourti-Stathaki and Proudfoot, *Genes Dev*, 2014), which are known substrates for ribosome stalling during translation. Third, ribosomes that stall on aberrant mRNA are normally rescued by the RQC pathway (Brandman and Hegde, *Nat Rev Mol Cell Biol*, 2016), but as described in Part II, RQC lacks demand-responsive upregulation and possesses self-limiting negative feedback, so that if stalling frequency exceeds a threshold, RQC capacity may be saturated. Fourth, oligodendrocytes and neurons impose the highest absolute translational demands in the central nervous system, so that even a modest increase in stalling frequency could exceed their fixed RQC capacity.

In this framework, MAM acts as a downstream recipient rather than an upstream driver. MAM-associated genes (TBK1, SQSTM1, VAPB, SPTLC1, ATG14) occupy co-expression topology hubs (raw φ upstream) but show no disease-specific residual shift (z ≈ 0; Results §6), which rules out MAM capacity saturation as an upstream driver. Truncated peptides generated by CATylation failure (Layer 1) may nonetheless accumulate at the ER membrane/MAM platform, causing functional contamination while co-expression structure remains intact. In this model, MAM is a downstream recipient of translational quality control failure, consistent with the Plessis-Belair et al. (2024) pathway in which NEMF dysfunction leads to Importin-β sequestration at the ER membrane.

A critical caveat applies to this chain: the R-loop-to-stalling bridge is logically plausible but *not directly supported by data in this study*. Part I φ analysis does not include NVU cell types. Part III ATM analysis does not measure RQC-related endpoints. We present this connection as an inferred hypothesis, not as an empirical finding.

#### S2.3 Proposed cascade: a five-layer model (original §12.3)

Drawing on the three independent datasets and the inferred R-loop-to-stalling bridge, we propose the following multi-layer cascade as a working model for sporadic ALS pathogenesis. Each layer is annotated with its evidential basis.

*Layer 0: NVU trigger (supported by Part III data).* ATM pathway activity declines in capillary endothelium, the earliest cell type affected in the module-level causal analysis. This decline is consistent with impaired double-strand break (DSB) repair, permitting R-loop accumulation and downstream RNA surveillance failure. The capillary-to-pericyte propagation pattern suggests that the initial perturbation originates within the neurovascular unit.

*Evidential basis: Part III, Sections 10.1-10.4: ATM pathway satisfies order, consistency, and pathway coherence criteria in capillary endothelium; cross-dataset validated against GSE212630 (Wang et al. 2023). Limited by small cell numbers.*

↓ *[Inferred connection: R-loop accumulation generates mRNAs with abnormal secondary structure, increasing ribosome stalling frequency (Supplementary Discussion S2, §S2.2). This step is not directly measured in this study.]*

*Layer 1: NEMF-associated CATylation-bottleneck hypothesis and collision-response initiation signature (NEMF co-expression loss supported by Part II 3φ residual data; mechanistic interpretation drawn from literature). NEMF co-expression coherence is selectively disrupted (z &lt; −2 in 7/10 cell types; Results §6), consistent with a specific bottleneck or coordination fracture at the CATylation step of RQC, while other components (LTN1, ZNF598, ABCE1) remain co-expression-coherent. Concurrently, collision-response ISR kinases GCN2 (5/10 CTs) and PKR (3/10 CTs) lose co-expression coherence, while PERK and ISR effectors are preserved. This pattern is consistent with preserved collision-sensor-associated co-expression but impaired initiation of collision-responsive signalling. Truncated peptides accumulate at the ER membrane, and chaperone networks activate compensatorily (4/10 CTs, translation-independent; Results §6).*

*Evidential basis:* Part II, Results §6: RQC design vulnerabilities documented from published literature (no positive feedback: Carneiro et al., *Sci Rep*, 2024; negative feedback via TCF25/LTN1: Brandman et al., *Cell*, 2012; RQC-ISR competition: Yan and Zaher, *Mol Cell*, 2021). φ data show chaperones CHORDC1 and ST13 as structurally most upstream in oligodendrocytes; ISR sensor EIF2AK4 positioned upstream (Part II, Results §6). The RQC overload model is a hypothesis consistent with these observations.

↓

*Layer 2: RNA sequestration (supported by Part II data). Published stress-granule biology provides a plausible mechanism for RNA sequestration of stalled mRNAs (G3BP2, CAPRIN1, and TIAL1 are canonical stress granule components); upon granule disassembly, sequestered mRNAs may re-enter translation, stall again, and generate additional aberrant peptides, potentially creating a self-reinforcing cycle. The present φ landscape does not directly establish this loop.*

*Evidential basis:* Part II, Results §6: Loop 2 of the triple feedback model. φ positions of stress granule components are observational; the re-stalling cycle is inferred from known stress granule biology (Protter and Parker, *Trends Cell Biol*, 2016).

↓

*Layer 3: TDP-43/FUS convergence (supported by Part I and Part II data).* Sustained stress granule accumulation promotes aberrant recruitment of TDP-43 and FUS. Liquid-liquid phase separation shifts toward solid-like aggregation. RNA sequestration worsens as aggregated RBPs lose normal function. TARDBP occupies a structurally downstream position (26.5 percentile in oligodendrocytes; Part II, Results §5), consistent with TDP-43 pathology representing a convergence point rather than an initiating event.

*Evidential basis:* Part II, Results §5: ALS-causative gene topological hub hierarchy places TARDBP downstream of TBK1, SQSTM1, FUS, and C9orf72 (note: this reflects co-expression topology, not disease-specific ordering; see Results §5 residual analysis). Part II, Results §6: NEMF CATylation failure leads to truncated peptide accumulation, which via the Plessis-Belair et al. (2024) pathway produces TDP-43 nuclear loss. Part I: curl component captures cyclic dynamics consistent with feedback processes involving RNA-binding proteins. The interpretation of TDP-43 as convergence point is a hypothesis consistent with both the topological hierarchy and the NEMF → TDP-43 mechanistic pathway.

↓

*Layer 4: Cell-type-specific collapse (supported by Part I and Part II data).* Each cell type's most vulnerable programme fails according to its specific upstream landscape:

- *Oligodendrocytes:* sphingolipid and myelin metabolism collapse (TECR, PSAP structurally upstream in Part II; oligodendrocytes identified as sole upstream cell type with φ = 0.900 in Part I).
- *Neurons (L4/6):* calcium homeostasis and axonal transport fail (MICU1, MICU3, KIF5C structurally upstream in Part II).
- *All cell types:* translation-resource stress remains structurally coherent across cell types (translation/ribosome programmes are the gradient source in 8/10 cell types by mean φ ranking; Part I, Results §2), while downstream quality-control coupling fails at the NEMF/CATylation bottleneck.

*Evidential basis:* Part I, Results §2: cell type φ rankings and gradient/curl decomposition. Part II, Results §6: cell type-specific upstream gene profiles. The interpretation of these as &quot;collapse programmes&quot; is inferential.

#### S2.4 Why oligodendrocytes may be affected first (original §12.4)

The identification of oligodendrocytes as the sole structurally upstream cell type (Part I, φ = 0.900) raises the question of why this cell type would be particularly vulnerable. Several considerations (none individually conclusive) converge on a consistent picture:

First, oligodendrocytes maintain among the highest translational loads of any cell type in the central nervous system, producing massive quantities of myelin membrane proteins (PLP1, MBP, MOG) and the lipid biosynthetic enzymes required for myelin assembly. This high baseline translation rate implies a correspondingly high absolute frequency of stochastic ribosome stalling events.

Second, oligodendrocytes are anatomically positioned within the neurovascular unit, directly exposed to perturbations originating in capillary endothelium. If Layer 0 (ATM brake loss) generates signals that propagate through the NVU, oligodendrocytes would be among the first parenchymal cells affected.

Third, the φ = 0.900 value for oligodendrocytes in Part I may reflect this combination of high translational demand and early NVU exposure, positioning oligodendrocytes as what the cascade model would predict to be the first parenchymal cell type to manifest detectable co-expression changes.

These considerations are individually speculative and collectively constitute a plausibility argument rather than a mechanistic demonstration.

#### S2.5 Familial-sporadic unification: entry points into a shared cascade (original §12.5)

If the multi-layer cascade model is approximately correct, it suggests a framework in which different ALS-associated mutations and sporadic disease may represent different entry points into a shared downstream pathway. Table 19 summarises this hypothetical mapping.

**Table 19. Hypothetical mapping of ALS genetic and sporadic forms onto cascade layers.**

| Genetic form / Cause | Primary layer affected | Proposed mechanism of entry | Supporting evidence |
| --- | --- | --- | --- |
| C9orf72 repeat expansion | Layer 1 (RQC overload) | Dipeptide repeat proteins (DPRs, especially poly-GR) stall ribosomes at the exit tunnel, directly activating RQC; RQC insufficiency leads to CAT-tailing product accumulation | Poly-GR ribosome stalling (Hartmann et al., *PNAS*, 2018); NEMF/LTN1 as genetic modifiers of DPR toxicity (Tseng et al., 2024) |
| FUS mutations | Layer 2 (RNA sequestration) | Mutant FUS disrupts stress granule dynamics, impairing normal mRNA triage and promoting aberrant RNA sequestration | FUS phase separation (Patel et al., *Cell*, 2015) |
| TARDBP mutations | Layer 3 (TDP-43 convergence) | Mutant TDP-43 accelerates liquid-to-solid phase transition, directly worsening RNA sequestration and feedback into Layers 1—2 | TDP-43 phase separation (Conicella et al., *Structure*, 2016) |
| VCP mutations | Layer 1 (RQC overload) | VCP/p97 is required for extraction of RQC substrates from stalled ribosomes; VCP loss impairs clearance of ubiquitylated stall products | VCP in RQC (Verma et al., *eLife*, 2013) |
| SOD1 mutations | Layer 1 (RQC overload) | Oxidative stress from misfolded SOD1 increases global ribosome stalling frequency through oxidative mRNA damage | Oxidative RNA damage and translation (Tanaka et al., *Nucleic Acids Res*, 2007) |
| TBK1 mutations | Layer 1—2 (RQC/autophagy) | TBK1 is required for selective autophagy of protein aggregates; loss impairs clearance of RQC-generated aberrant peptides | TBK1 φ position: 5.7 percentile in oligodendrocytes (Part II) |
| Sporadic ALS | Layer 0—1 | Age-dependent NVU deterioration (Layer 0) erodes R-loop surveillance capacity. Gradual accumulation of stalling events exceeds NEMF CATylation capacity (Layer 1), leading to collision response initiation failure in the seventh or eighth decade. Truncated peptides accumulate at the ER membrane/MAM, causing downstream functional contamination | No single mutation required; cumulative age-related decline in translational quality control |

This table is explicitly hypothetical. The mapping is constructed post hoc to be consistent with known biology and with the φ hierarchy observed in Part II. It does not constitute independent evidence for the cascade model but rather illustrates the model's internal consistency with established genetics. Several entries in the "proposed mechanism" column draw on published literature but have not been tested in the specific context of this cascade.

#### S2.6 Testable predictions from the integrated model (original §12.6)

A model that cannot be falsified has no scientific value. We therefore derive five predictions from the cascade model, each of which is specific enough to be tested with existing or near-term technologies and, if refuted, would undermine the corresponding layer of the model.

*Prediction 1 (Layer 0: pre-symptomatic vascular changes).* If the NVU trigger hypothesis is correct, capillary endothelial cells in pre-symptomatic ALS mutation carriers should show reduced ATM pathway activity and elevated markers of unresolved DSBs (e.g., persistent gamma-H2AX foci) *before* motor neuron loss is detectable.

*Technology:* Spatial transcriptomics or immunohistochemistry on pre-symptomatic C9orf72 carrier brain tissue. *Falsification criterion:* If ATM pathway activity is normal in pre-symptomatic NVU cells across multiple carriers, Layer 0 as formulated would be refuted.

*Prediction 2 (Layers 0—1 bridge: R-loop and stalling co-occurrence).* If R-loop accumulation mechanistically increases ribosome stalling, then paired DRIP-seq (R-loop mapping) and disome profiling (ribosome collision mapping) on the same samples should show a positive correlation between R-loop burden and stalling frequency at overlapping genomic loci.

*Technology:* DRIP-seq + Ribo-seq/disome profiling on ALS versus control post-mortem tissue or iPSC-derived motor neurons. *Falsification criterion:* If R-loop burden and ribosome stalling frequency are uncorrelated or negatively correlated, the proposed R-loop-to-stalling bridge would be refuted.

*Prediction 3 (Layer 1: NEMF CATylation rescue).* If NEMF CATylation is the rate-limiting bottleneck, then NEMF overexpression (not LTN1) in ALS model systems should reduce CATylation burden and may attenuate downstream TDP-43 pathology, while LTN1 overexpression alone should have minimal effect (because its substrate supply via NEMF is disrupted).

*Technology:* AAV-mediated LTN1/NEMF overexpression in SOD1-G93A or C9orf72 BAC transgenic mice. *Falsification criterion:* If RQC augmentation produces no delay in motor phenotype onset across multiple model systems, Layer 1 as the critical bottleneck would be undermined.

*Prediction 4 (Layers 1—2: pre-symptomatic oligodendrocyte stress).* If oligodendrocytes are the first parenchymal cell type affected, pre-symptomatic tissue should show elevated CAT-tailing products (RQC failure markers) and truncated peptides specifically in oligodendrocytes, before such markers appear in motor neurons.

*Technology:* Cell-type-specific mass spectrometry (e.g., TRAP-MS) or proximity labelling in pre-symptomatic mutation carrier tissue. *Falsification criterion:* If RQC failure markers appear first in motor neurons rather than oligodendrocytes, the oligodendrocyte-first model (and by extension the φ = 0.900 interpretation) would require revision.

*Prediction 5 (Layers 1—3: CSF biomarkers).* If the cascade model reflects the temporal order of disease, cerebrospinal fluid (CSF) from early-stage ALS patients should contain elevated levels of RQC-associated truncated peptides and CAT-tailing products, potentially before TDP-43 fragments become detectable. The temporal order of biomarker appearance in longitudinal CSF samples should recapitulate the layer order: RQC substrates (Layer 1) before stress granule markers (Layer 2) before TDP-43 fragments (Layer 3).

*Technology:* Longitudinal CSF proteomics in pre-symptomatic mutation carriers followed through conversion. *Falsification criterion:* If TDP-43 fragments consistently appear in CSF before RQC-related truncated peptides, the proposed layer ordering would be contradicted.

#### S2.7 Final limitations and epistemic boundaries (original §13)

##### Limitations of the integrated model

The cascade model proposed in Supplementary Discussion S2 is subject to the following limitations, which collectively place it at the lowest confidence level within this paper.

- The Layer 0 to Layer 1 connection is inferential, not empirical. The bridge between ATM brake loss (Part III) and RQC overload (Part II) passes through R-loop accumulation and increased ribosome stalling, a sequence that is mechanistically plausible but not measured in any dataset analysed in this study. Part I φ analysis does not include neurovascular cell types. Part III module-level analysis does not assess RQC-related molecular endpoints. The two analytical pipelines were not designed to be connected, and the proposed connection is a post hoc interpretive act.
- The triple feedback loop model fills gaps in the φ landscape with hypothetical structure. φ measures structural precedence within the covariance geometry of gene co-expression change. Systems that fail silently (such as RQC, whose components maintain near-constant expression while becoming functionally insufficient) fall outside what φ can detect. The triple feedback loop (RQC overload, RNA sequestration, TDP-43 convergence) is therefore partly constructed to explain what φ *cannot* see, which is an inherently less constrained exercise than interpreting what φ directly measures.
- All Part I–III findings derive from a single primary cohort (NYGC ALS Consortium); the general limits this imposes are described in Discussion. Cross-cohort partial validation (Part V) addresses this for the Part I cell-type-level structural architecture: the Oligo-preserved / ET-sink condition-displacement φ pattern is 3-way taxonomy-robust across NYGC + Takeuchi + Ruf2026 (Results §7, 17), and the Part II §8.9 NEMF gene-level finding shows predominantly negative direction across all three cohorts (with breadth reproduced in Ruf2026 but not Takeuchi) under the same 3φ poly3-residual instrument (Results §7). However, several cross-cohort claims (Microglia preservation, Astrocyte upstream, donor-level PT-C distribution as ET-sink corroboration, dying-back temporal propagation) did not generalise and were not retained after locked sensitivity analyses. Most importantly, the Part III ATM brake loss / NVU trigger finding has NOT been independently validated in the cross-cohort meta (the meta-analysis used the seven common broad CTs across cohorts, not the vascular subtypes that Part III analyses); for the integrated model, the Part III leg therefore retains its single-cohort discovery status. The cascade as a whole remains a working hypothesis until Part III is independently replicated and the NVU-to-RQC bridge (Layer 0 → Layer 1) is empirically tested.
- Post-mortem tissue captures disease end-state, not progression. All analysed tissue represents the terminal phase of disease. The φ instrument infers structural precedence from cross-sectional data, and pseudotime ordering reconstructs a hypothetical progression axis, but neither directly observes the temporal unfolding of disease. Pre-symptomatic validation (Predictions 1 and 4) is required to determine whether the inferred ordering reflects actual disease chronology.
- φ remains structural precedence, not causation. This point, developed in Sections 4.4 and 5, applies with particular force to the integrated model: the cascade's layer ordering is derived from φ rankings and is consistent with causal priority but does not constitute causal evidence. Interventional experiments (Prediction 3) are needed to test causal claims.
- The familial-sporadic unification table is post hoc. Table 19 maps known ALS mutations onto cascade layers in a manner consistent with published molecular biology. This mapping was constructed after the cascade model was formulated and does not provide independent support for the model. It illustrates internal consistency, not predictive power.
- Cell-type coverage is incomplete. Part I analyses ten cell types from cortical snRNA-seq; Part III analyses neurovascular cells from the same dataset but through a separate pipeline. Microglia, astrocytes, and peripheral immune cells (all implicated in ALS by other studies) are not positioned within the cascade as primary actors, which may reflect analytical limitations rather than biological irrelevance.

##### Independence of empirical findings from the integrated model

A central design principle of this paper is that each Part's findings are independently valid regardless of whether the integrated model in Part IV is correct. We make this modularity explicit.

The Hodge-decomposition instrument of Part I stands independently. Its validity rests on external benchmarks: recovery of known causal structure in Perturb-seq data (Norman et al., 2019), spatial CRISPR screens (Shen et al., 2026), and temporal ground truth (Chao et al., 2026, TimeVault). These validations do not depend on any ALS-specific biological claim. If the cascade model is entirely wrong, the instrument remains an externally benchmarked framework for inferring structural upstream organisation from static snapshot data.

The sALS biological findings of Part I also stand independently. The observations that translation/ribosome programmes are structurally upstream in 9/9 cell types, that oligodendrocytes are the sole upstream cell type (φ = 0.900), and that disease structure decomposes into irreversible gradient and cyclic curl components are descriptive findings about the covariance geometry of sALS gene expression. They do not require the cascade model to be meaningful. Even if the mechanistic interpretation is incorrect, the structural pattern itself is a reproducible observation (LODO 24/24 stable).

The gene-level φ landscape of Part II stands independently as well. The cell type-specific upstream gene profiles (chaperones and sphingolipid enzymes in oligodendrocytes; ganglioside metabolism and calcium regulators in L4/6 neurons) and the ALS-causative gene hierarchy (TBK1 upstream, TARDBP downstream) are observational results of the φ analysis. They do not depend on the RQC vulnerability hypothesis. The hypothesis provides one possible mechanistic interpretation, but the φ rankings themselves are data.

The ATM brake loss finding of Part III stands independently too. ATM pathway decline in capillary endothelium, satisfying pre-specified order, consistency, and pathway coherence criteria, is a finding of the module-level causal analysis pipeline. Its status as the top-ranked causal candidate does not depend on whether it connects to downstream RQC overload. Even if the NVU-to-RQC bridge is incorrect, ATM brake loss in capillary endothelium remains a causal candidate worthy of independent investigation.

In summary, the cascade model is an optional interpretive layer. Its refutation would not invalidate the empirical contributions of Parts I—III. The paper is structured so that readers may accept any subset of its findings without logical contradiction.

**Cross-cohort modularity (Part V). Part V's cross-cohort meta-analysis adds another modular layer. The Part V cardinal claim (3-way taxonomy-robust ET-sink + Oligo-preserved structural architecture; cross-cohort-supported, predominantly negative NEMF signal) stands on its own as a structural-displacement validation independent of the Part II RQC vulnerability hypothesis and the Part III–IV cascade model. If the cascade model is incorrect, the cross-cohort structural architecture remains valid as an empirical observation about three independent BA4/MCX sALS cohorts. Conversely, if the cross-cohort generalisation were to fail (e.g., in a fourth larger cohort), the Part I–II NYGC findings would still stand as a discovery cohort observation, and the Part III ATM/NVU finding would still stand as a single-cohort module-level causal candidate. Each layer fails or survives independently, and the paper reports the boundaries of validation honestly at each layer.**

##### Closing statement on epistemic boundaries

This paper has attempted to maintain a consistent epistemic gradient: from externally benchmarked instrument (Part I) through exploratory analysis (Part II) and limited-sample causal candidacy (Part III) to speculative integration (Part IV). We close by restating the boundaries that this gradient imposes.

Concretely, the instrument identifies structural precedence, not causes; the gene-level φ landscape reveals which programmes are positioned upstream in co-expression change dynamics, not which genes initiate disease; the ATM brake loss finding identifies a causal candidate satisfying observational criteria, not a proven cause of ALS; and the cascade model proposes a sequence in which these observations could be connected, not the mechanism of ALS.

The contribution of this work is a set of structured observations, an externally benchmarked analytical instrument, and a falsifiable hypothesis. The observations constrain future mechanistic models. The instrument is applicable to other diseases and datasets. The hypothesis generates predictions that can be tested, and if the predictions fail, the observations and the instrument remain.
